# Evaluating the Utilities of Foundation Models in Single-cell Data Analysis

**DOI:** 10.1101/2023.09.08.555192

**Authors:** Tianyu Liu, Kexing Li, Yuge Wang, Hongyu Li, Hongyu Zhao

**Affiliations:** Interdepartmental Program in Computational Biology & Bioinformatics, Yale University, CT, USA.; Department of Biostatistics, Yale University, CT, USA.

**Keywords:** single-cell data, large language model, foundation model, deep learning, benchmark

## Abstract

Foundation Models (FMs) have made significant strides in both industrial and scientific domains. In this paper, we evaluate the performance of FMs for single-cell sequencing data analysis through comprehensive experiments across eight downstream tasks pertinent to single-cell data. Overall, the top FMs include scGPT, Geneformer, and CellPLM by considering model performances and user accessibility among ten single-cell FMs. However, by comparing these FMs with task-specific methods, we found that single-cell FMs may not consistently excel than task-specific methods in all tasks, which challenges the necessity of developing foundation models for single-cell analysis. In addition, we evaluated the effects of hyper-parameters, initial settings, and stability for training single-cell FMs based on a proposed **scEval** framework, and provide guidelines for pre-training and fine-tuning, to enhance the performances of single-cell FMs. Our work summarizes the current state of single-cell FMs, points to their constraints and avenues for future development, and offers a freely available evaluation pipeline to benchmark new models and improve method development.

## 1 Introduction

Single-cell sequencing technologies offer high-throughout observations into complex biological systems at the cell level with multimodal data [1, 2]. They help elucidate disease mechanisms and potential treatments [3–5]. In line with the central dogma of molecular biology, these technologies enable the characterization of various molecules, such as DNA (e.g. scDNA-seq) [6], RNA (e.g. scRNA-seq) [7, 8], and proteins (e.g. Citeseq) [9]. Furthermore, single-cell sequencing can facilitate epigenetic studies, including chromatin accessibility (e.g. scATAC-seq) [10, 11] and methylation [12, 13]. These technologies have been rated as among the most impactful ones in recent years [14, 15].

With the development of single-cell technologies, expansive single-cell datasets have been collected, and they present challenges in integration, interpretation, and downstream utilization. Therefore, there is a need to build a Foundation Model (FM) which can benefit from prior knowledge for single-cell data analysis. In a similar motivation, natural language processing (NLP) also boasts extensive datasets, where FMs such as pre-trained Large Language Models (LLMs) have shown great success in addressing NLP tasks or multimodal tasks [16]. Numerous LLMs, including GPT-4 [17] and LLaMA [18], excel at diverse language-related tasks such as question answering and sentence generation, which has received widespread attention from both the AI community and society [19]. Moreover, these LLMs showcase impressive performance in zero-shot learning, thereby enabling them to address tasks beyond their original training scope, such as solving mathematical problems [20]. Similarly, a good FM in single-cell data analysis should also handle multiple tasks with a unified framework.

Indeed, notable parallels exist between studies based on single-cell sequencing data and those in NLP. Both fields leverage advanced technologies, such as transformer architectures, which have proven effective in handling a variety of tasks within each domain [21–24]. Similarly, both disciplines emphasize the importance of analyzing intra-data and inter-data relationships, which in the context of single-cell analysis might involve interactions among genes or cells [23, 25–27]. Moreover, the concept of representation learning is central to both fields: effective embeddings of cells and genes can be developed using techniques analogous to those used for generating token embeddings in NLP [28]. Lastly, the success of both fields hinges on access to high-quality databases [16, 29], underscoring the need for careful data selection during model training. These synergies suggest that integrating methodologies from NLP may significantly advance the analysis and interpretation of single-cell data.

While FMs have seen marked success in the realms of DNA analysis [30] and biomedical NLP [31, 32], their application in single-cell research remains largely uncharted. There is a limited number of robust pre-trained models (known as single-cell FMs) capable of managing multiple tasks in single-cell research. Some single-cell FMs focus on cell-type annotation or gene function prediction, including scBERT [33], CellLM [34], and Geneformer [35], while others aim to create a foundation model in this area that can handle multiple tasks, including scGPT [36], scFoundation [37], tGPT [38], GeneCompass [39], SCimilarity [40], UCE [41], and CellPLM [42]. Details of these models can be found in Appendices E and F. Furthermore, no studies to date have comprehensively evaluated the utility of these models and provided guidance for model training. Little has been done to compare NLP-focused LLMs with those used for single-cell research to gain insight into scaling laws and zero-shot (or few-shots) learning abilities of the latter.

Here we investigate the overlap of the pre-training stages of different single-cell FMs, and present a framework for assessing various single-cell FMs and tasks (shown in Figure 1 (a)), termed as Single-cell Large Language Model Evaluation (**scEval**), shown in Figure 1 (b). Using scEval, we not only compare different single-cell FMs across various datasets and tasks as a horizontal comparison but also identify critical parameters and strategies for the fine-tuning process of specific models as a vertical comparison. We also examine the potential contributions of model scaling of single-cell FMs, substantiating that the latter also possesses distinctive abilities. To help the audience better understand FMs, we provide a glossary summary of common terms in Artificial Intelligence (AI) and Machine Learning (ML) in Supplementary file 1.

**Fig. 1:**
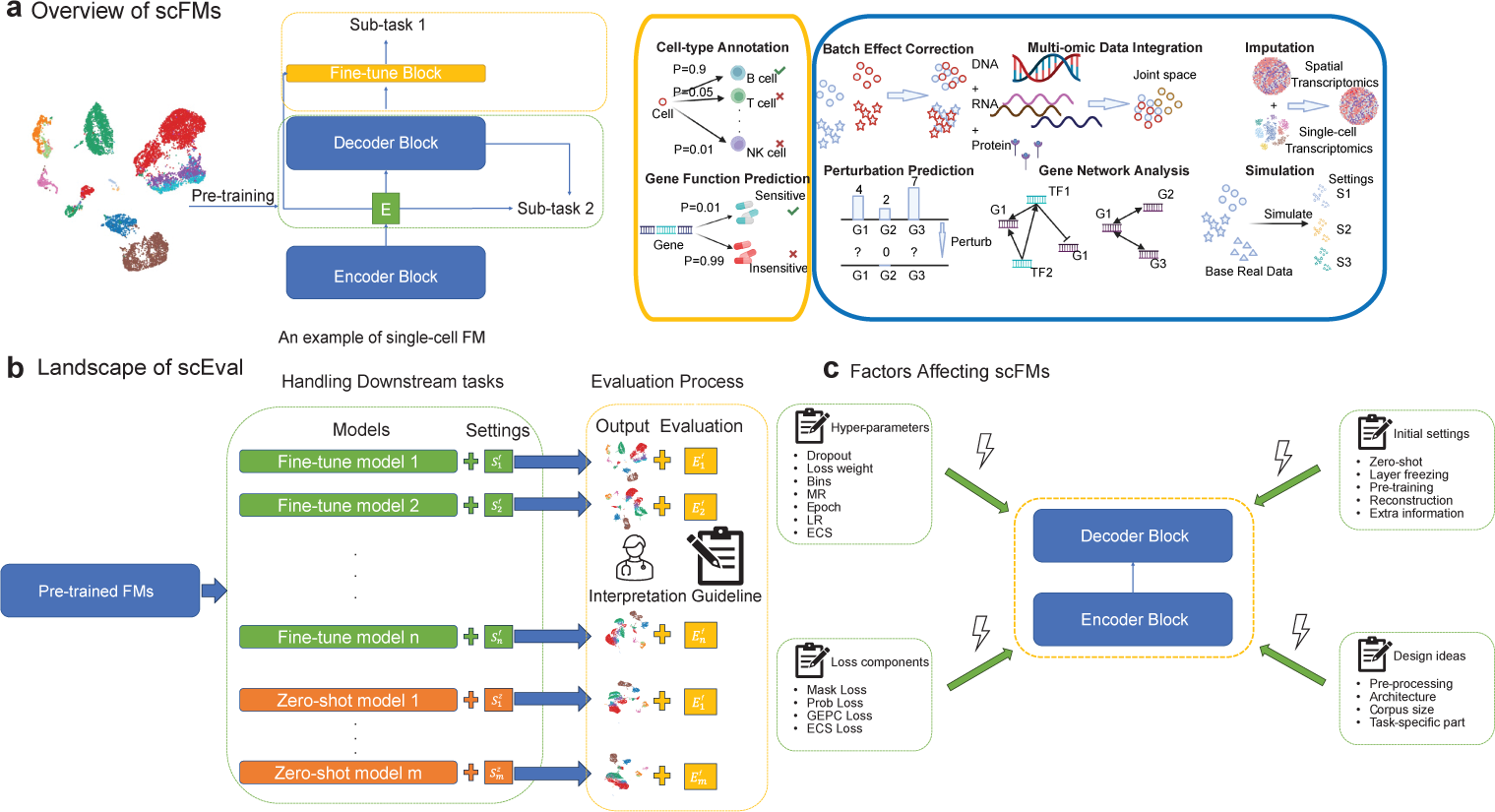
Overview of single-cell FMs, landscape of scEval and factors affecting single-cell FMs. (a): Overview of single-cell FMs describing the typical structure of FMs and general tasks for single-cell data analysis. The right two blocks represent two types of downstream tasks. Yellow block: Sub-task 1, including Cell-type Annotation and Gene Function Prediction (top to bottom). Blue block: Sub-task 2, including Batch Effect Correction, Multi-omic Data Integration, Imputation (From left to right, top row), Perturbation Prediction, Gene Network Analysis, and Simulation (From left to right, bottom row). (b): The landscape of scEval shows the workflow of our systematic evaluation. Here we consider models based on fine-tuning design with settings *S^f^*_1_ to *S^f^_n_*, and they have corresponding evaluation pipelines *E^f^*_1_ to *E^f^_n_*. We also consider models based on zero-shot-learning design with settings *S^z^*_1_ to *S^z^_m_*, and they have corresponding evaluation pipelines *E^z^*_1_ to *E^z^_m_*. The reason for using the above notation to represent the process of evaluation is that different models can perform different tasks, while different tasks have different evaluation scenarios. (c): Factors which can affect the performance of single-cell FMs. These factors are summarized based on different designs of single-cell FMs. The known factors can be classified into four different types. Details of hyper-parameters can be found in Appendix A. Details of initial settings can be found in Appendix B.

## 2 Results

### Overview of our evaluations

We evaluated the performance and user accessibility of nine open-source single-cell FMs (scGPT, Geneformer, scBERT, CellLM, tGPT, SCimilarity, CellPLM, UCE, and scFoundation) and one closed-source single-cell FM (GeneCompass) by assessing their outputs on eight tasks with 29 datasets. We did not evaluate all the models with all the datasets with the reasons provided in Supplementary files 2 and 3, but tried our best to implement the missing functions for evaluation. The tasks that can be performed for different models as well as the overall ranks are summarized in Tables 1 (a) and (b). We also compared their performance with state-of-the-art (SOTA) task-specific methods. Our workflow can be summarized in Extend Data Figure 1. The top three methods include scGPT, CellPLM, and Gene-former by considering both usability and performance. For each task, we evaluate single-cell FMs based on their default settings for a fair comparison. Moreover, we discuss the effect of different parameter settings on model performance and investigate the contribution of different loss functions of scGPT and initial settings by ablation tests. We also consider the contribution of model scales to the performance of FMs. Finally, we evaluate the stability and usability of different single-cell FMs and make recommendations for preferred models. The detailed design of scEval is explained in the Methods section. To ensure fairness, we selected metrics for different tasks based on their benchmarking analysis, and we fixed all models to be from versions before July 1, 2024.

**Tab. 1:**
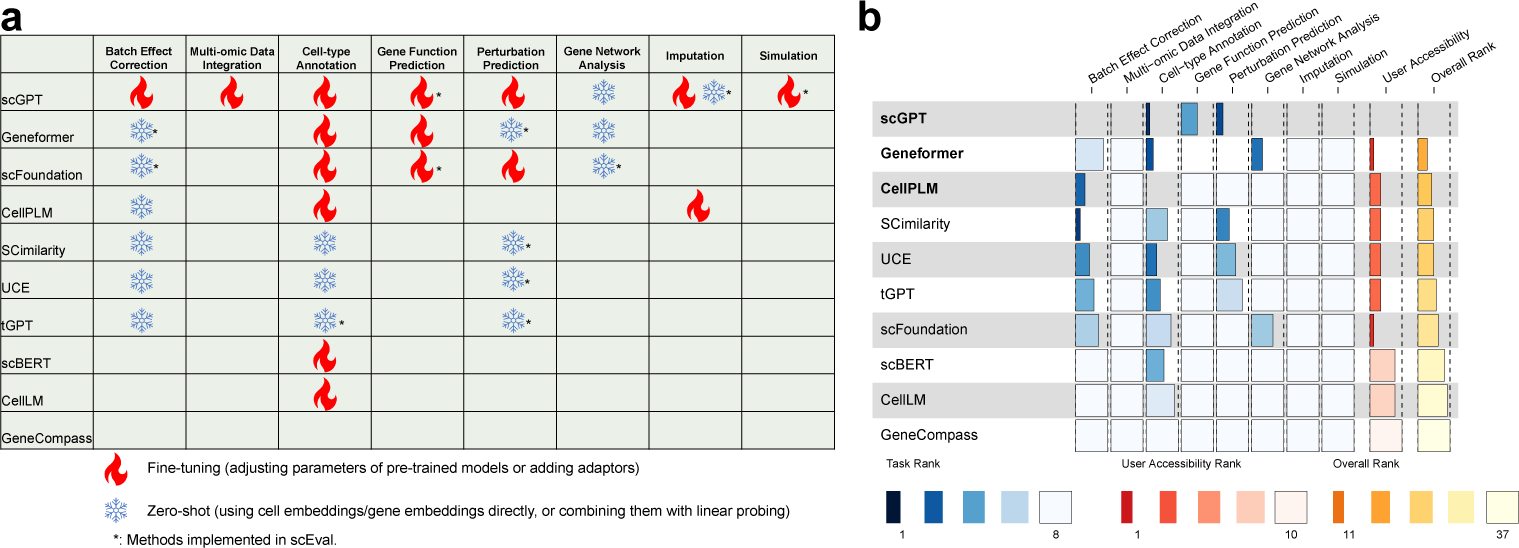
Overall comparisons of model capacity and performance. (a) Evaluations for the capacity of different single-cell FMs. We utilized different signs to denote the default settings we tested in scEval. Details are explained in the table legend. The icons are inspired by [43]. (b) Evaluations for both performance and usability of different single-cell FMs. We record the ranks of different methods across different tasks and user accessibility evaluations and compute the final overall rank. A lower rank means a better model. The top three methods are boldfaced.

### Comparisons of pre-training datasets for different single-cell FMs

In this section, we investigated and compared the pre-training steps of different single-cell FMs from three dimensions, including the scale of pre-training datasets, the diversity of pre-training datasets, and the major human tissues or organs overlap of pretraining datasets. Based on Extended Data Figure 2, we found that GeneCompass had the largest pre-training datasets, followed by scFoundation. However, GeneCompass is closed-source. Interestingly, the pre-training scales of UCE, scGPT, Geneformer, SCimilarity, and tGPT are comparable, which implies that having 20-30 million may be acceptable for constructing pre-training datasets. Moreover, SCimilarity had the largest diversity in the datasets conditioned on diseases compared with other methods. Most of the single-cell FMs chose to include cancer cells in their pre-training process, which was possibly due to many single-cell studies focused on cancer [44]. UCE had the largest diversity in the datasets from different species, while the rest of the single-cell FMs focused more on human. In addition, few models considered the use of multi-omic data information, except tGPT, CellPLM, and UCE. However, in the pre-training process, UCE and tGPT omitted the extra information (for example, the spatial location of spatial transcriptomic data) provided by these datasets and treated the data the same as scRNA-seq data. Using different pre-training pipelines for multiomic datasets might improve the performance of current models on addressing related tasks. In Extended Data Figure 3, we computed the overlap of major human tissues or organs used for pre-training across different models. Geneformer had the largest diversity in this comparison because we treat it as a reference method. UCE had the largest overlap compared with Geneformer. scBERT and CellLM shared major human tissues or organs because they both used PanglaoDB [45] for pre-training. Interestingly, scGPT did not use many types of tissues for pre-training. Considering its relatively better performance, including as many types of datasets as possible may not necessarily lead to better performance, and data ablation analysis is needed for the pre-training step.

Furthermore, we analyzed the relationship between these statistics and model performance. Here, we considered two major tasks, batch effect correction, and cell-type annotation, for our comparison. We compared four FMs for batch effect correction and seven FMs for cell-type annotation, thus offering statistical meaning to perform further investigation. For each task, we computed the Spearman correlation under three settings, including performance vs. pre-training data scale, performance vs. the number of major tissues for pre-training, and performance vs. the number of model parameters. However, all the six correlations computed based on the comparisons did not show statistical significance (p-value *>* 0.05). Therefore, the relationship between the performance of single-cell FMs on downstream tasks and their pre-training settings might be affected by many factors, including pre-training strategies, data-cleaning pipelines, rules of training-testing datasets splitting, and others. Therefore, we focus on task-driven analysis and offer guidelines for model development based on their performances on different tasks.

Finally, we include a table that represents whether our evaluation datasets are used for the pre-training of single-cell FMs with known pre-training information in Supplementary file 2. This table shows that most of the datasets were not included by all single-cell FMs, so we had a sparse table.

### Evaluation based on cell-perspective tasks shows that single-cell FMs can reduce batch effect and annotate cell types

#### Batch Effect Correction

For this task, we intend to reduce batch effect of scRNA-seq datasets. We considered scGPT, tGPT, UCE, SCimilarity, CellPLM, Geneformer, scFoundation, Harmony [47], and ResPAN [48] for this task. We also provided a detailed analysis of the influence of various hyper-parameters on the performance of scGPT on batch effect correction. The description of the evaluation metric *S_final_* can be found in Appendix D. We computed *S_final_* based on the weighted average of metrics for evaluating the level of batch effect removal and the metrics for evaluating the level of biological variance conservation. As shown in Figure 2 (a), scGPT outperformed ResPAN in one of the 11 datasets and outperformed Harmony in two of the 11 datasets, while Harmony had an overall best performance. In the comparison of FMs, scGPT v1 had the best performance, and scGPT v1 outperformed the scGPT model on average, raising the issue of the need for increasing the size of pre-training datasets for this task. Moreover, only Harmony and ResPAN achieved better *S_final_* compared with the raw datasets on average. Therefore, the ability of FMs to remove the batch effect is limited. Moreover, these FMs had worse performance in reducing the batch effect for large-scale datasets. One reason is that their biology conservation scores were lower than those of raw datasets (shown in Extended Data Figure 4). UCE and scFoundation could not handle large-scale datasets due to running errors, which also raised problems in model accessibility. Extended Data Figures 6 and 7 show the Uniform Manifold Approximation and Projection (UMAP) [49] plots for the raw data and the scGPT results. We could still observe the batch effect in the output of scGPT for Pancrm, HumanPBMC, MCA, MHSP, DC, Lung atlas, Immune atlas, and Heart atlas datasets. We also evaluated the performances of single-cell FMs and task-specific methods for preserving the trajectory information during the batch effect correction process, shown in Extended Data Figure 5, and the metric used here is trajectory score. Based on this figure, Harmony still had the best performance, followed by scGPT.

**Fig. 2:**
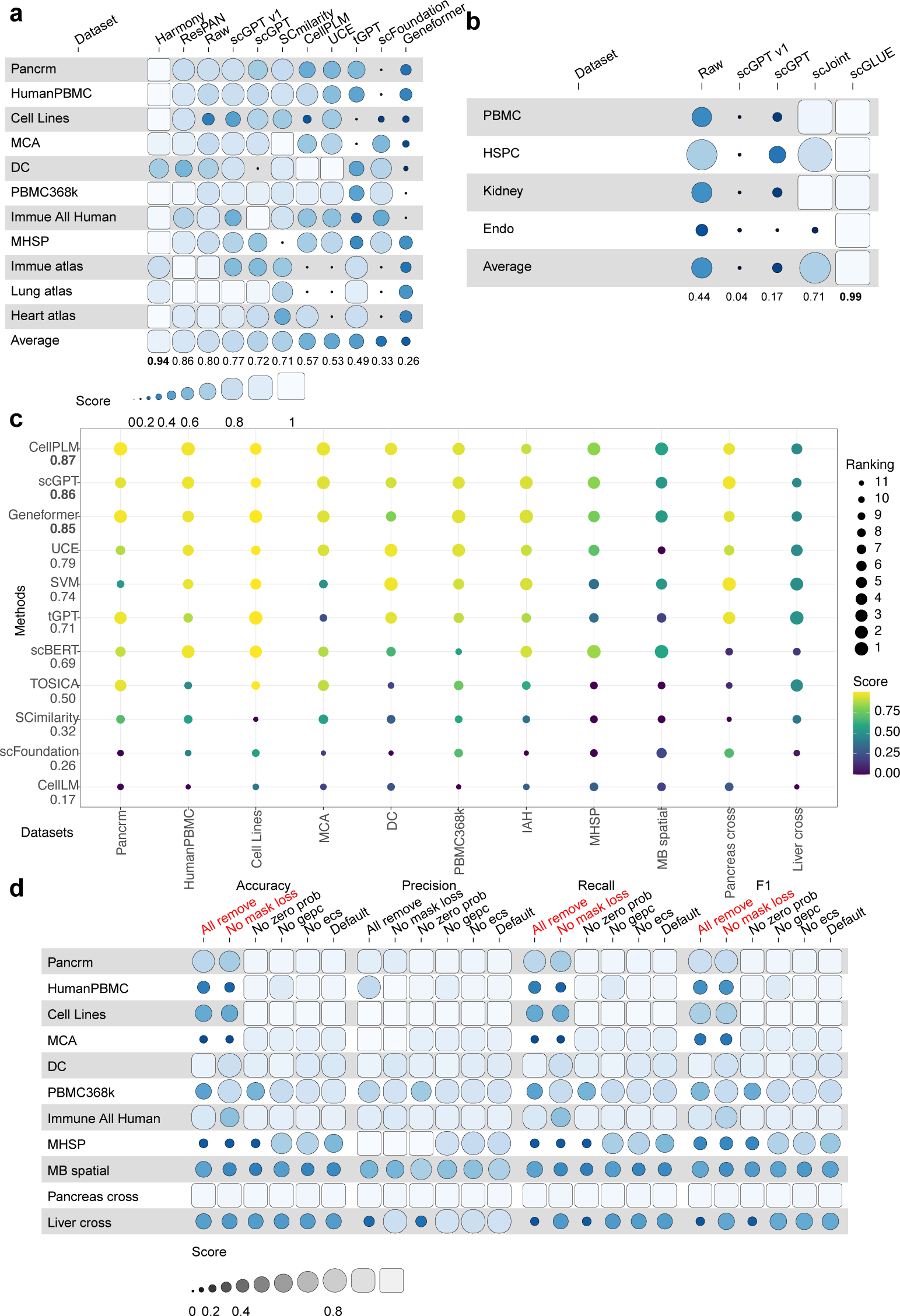
Experimental results of single-cell FMs and benchmarking methods for cell-Extended Data Figure 9 (a) presents the comparison of scores across different initial settings for the batch effect correction task using scGPT. We can see that scGPT is capable of performing zero-shot learning in batch effect correction. For the Cell Lines dataset, the zero-shot learning approach even achieved the highest score, indicating that it may be an effective method for certain datasets. Moreover, pretraining significantly contributes to the performance of scGPT in the batch effect correction task. Without pre-training, the model’s performance notably decreased. Using cross-entropy as the loss function for gene expression reconstruction yielded better results than the mean square error (MSE) loss for most datasets. Freezing weights is not crucial for batch effect correction. Interestingly, the encoder structure appears to play a more important role in the training process, as freezing the encoder layer led to a larger decrease in the score. Incorporating the cell type information as a human label into the training process enhanced performance for most datasets. These observations suggest that each dataset may require unique tuning, underscoring the importance of adaptable methodologies in single-cell RNA-seq data analysis.

We provide a detailed analysis of the impact of various hyper-parameters on the performance of scGPT in batch effect correction based on Extended Data Figure 8. A smaller learning rate tended to lead to better performance across all datasets. The optimal number of training epochs varied across datasets, with a larger number of epochs being beneficial for most datasets. This result contradicts recent research advocating for a single-epoch training approach [50], suggesting that the optimal number of epochs might be context-dependent. Increasing the number of bins is generally associated with an increase in the final score. The impact of the mask ratio and dropout rate on model performance is unclear, suggesting that further investigation is needed to understand their influence. These observations may improve the application of scGPT for batch effect correction in single-cell data analysis and may also inform fine-tuning of other similar models.

Extended Data Figure 9 (b) shows the performance metrics versus the choices of different optimizers for the batch effect correction. Adam and AdamW [51] had comparable performance in this task, while SGD [52], Sophia-G [53] (a novel optimizer which is designed for training FMs) and Lion [54] (an optimized version of Adam) were worse. Therefore, the optimizers in the Adam family are preferred.

Extended Data Figure 9 (c) illustrates the impact of different loss function components on the performance of batch effect correction using scGPT. Here we consider Mask Loss (It means we mask the gene expression levels of some genes and reconstruct them to compute the loss), Prob Loss (i.e. we predict whether some genes are expressed in some cells to compute the loss), GEPC Loss (i.e. we predict the gene expression levels based on the cell embeddings to compute the loss), ECS Loss (i.e. we bring similar cells closer together and push different cells farther apart in the embedding space to compute the loss), and one task-specific loss function, known as the gradient reverse loss (i.e. we reverse the gradient of batch label classifier to perform adversarial training). We compared the scGPT without certain loss functions with its default mode to perform ablation tests. Using all components of the loss function did not always yield the best results, with the exceptions of the Pancrm, MCA, and MHSP datasets. Using only the gradient reverse loss function resulted in the worst performance. The GEPC Loss seemed to play a crucial role in the performance of the batch effect correction task. These results suggest the need for a careful composition of the loss function when training single-cell FMs for batch effect correction, with each loss function component contributing differently to model performance.

#### Multi-omic Data Integration

For this task, we intend to integrate unpaired scRNA-seq datasets with scATAC-seq datasets. We considered scGPT, scJoint [55], and scGLUE [56] for evaluation. We assessed the integration quality through the same score as Batch Effect Correction because these two tasks have similar targets, that is, integrating datasets from different domains while preserving biological variation. The results presented in Figure 2 (b) summarize the evaluation results for the multi-omics integration task. Overall, scGLUE surpassed scGPT and achieved the best performance. In addition, the Extended Data Figure 10 shows that the results of scGPT did not have better biology conservation scores than the raw condition across all the datasets. Therefore, there exists a performance gap between scGPT and task-specific models. Based on our analysis, scGPT still performed better than scGPT v1, which implied that larger pre-training datasets might help integrate multi-omic datasets. We also observed batch effect for the results of scGPT based on UMAP plots shown in Extended Data Figure 11. In conclusion, single-cell FMs did not show an advantage in handling datasets for multi-omic data integration.

We illustrate the effect of initial settings in Extended Data Figure 9 (d). Different from the case with batch effect correction, the cross-entropy loss function led to worse performance compared to the MSE loss for this task. Interestingly, pre-training did not significantly affect the performance for this task since training from scratch also had similar performance. The encoder part of the single-cell FM played a more important role than the decoder since we observed a larger performance drop by only freezing the encoder part. Including cell types or human labels in the training process proved beneficial, likely providing the model with more precise and useful information for the task. The zero-shot learning approach did not perform as well for this task as it did for batch effect correction. Therefore, we need more consideration for the design of single-cell FMs for this task.

We illustrate the evaluation metrics versus different parameter settings in Extended Data Figure 12. scGPT did not perform well on this task as shown by the low score (below 0.5) even if we tried to search for better hyper-parameters. Certain hyperparameters affected the training process. A smaller weight for the loss function, a larger dropout rate, and more epochs improved the model’s performance. The number of bins and mask ratio did not exhibit monotonicity, making it difficult to draw conclusions. Setting a high learning rate and ECS weight decreased the performance of scGPT. Moreover, based on Extended Data Figure 9 (e), the optimizers in the Adam family are still preferred because of better performance, which is similar to what we found for the batch effect correction task. Since the patterns discovered in the multi-omic data integration task are not exactly the same as what we discovered in the batch effect correction task, we believed that specific design focusing on scATAC-seq data is required.

#### Cell-type Annotation

For this task, we intend to annotate cell types for scRNA-seq datasets. We considered all open-source foundation models, TOSICA [23], and SVM_rej_ [57, 58] for this task. We assessed the performance of different single-cell FMs in assigning cell types based on the four metrics (Accuracy, Precision, Recall, and F1 score) discussed in Appendix D.2. These metrics are widely used in the evaluation for cell-type annotation. The UMAPs for the raw data and scGPT are shown in Extended Data Figures 13 and 14. Since these plots were generated based on the same cell embeddings, they could be used as visual assessment for the observed and predicted distributions of cell types. Figure 2 (c) displays the Accuracy for different settings for these models. On average, models with pre-training performed better than those without pre-training. However, CellLM and scFoundation did not perform well across all the datasets, mainly because these two methods had running errors for Pancrm, HumanPBMC, PBMC368k, and Liver cross datasets, which raised the issue of the reliability and usability of these models again. SCimilarity also did not perform well in this task, which might be due to their pre-training process which contained too many sub-cell types and only utilized 10X sequencing data [40]. Moreover, for the intra-dataset prediction task, CellPLM, scGPT, and Geneformer were comparable although they had different pre-training settings. For the inter-dataset prediction task, CellPLM, scGPT, Geneformer, tGPT and UCE were better than scBERT. Therefore, different single-cell FMs also had large divergences in performance, but CellPLM, scGPT, and Geneformer had good performances across different datasets.

In Extended Data Figure 15, we compared the performance of models with different hyper-parameter settings. Higher loss weight, learning rate, ECS threshold, mask ratio, and smaller epochs tended to lead to worse performance of scGPT. There was little correlation between the number of bins and the performance of scGPT. We observed the consistency in the performance of different single-cell FMs under the condition of altering their shared hyper-parameters. For Geneformer and scBERT, a lower learning rate and higher epochs also tended to lead to better performance.

We also considered different initial settings for model training. The first setting is *Freeze all*, where we froze all the weights of pre-trained layers. The second setting is *Default*, where we used the default fine-tuning settings. The third setting is *From scratch*, where we did not use the pre-trained weights. Extended Data Figure 16 (a) shows the score versus initial settings across different datasets. Here we considered scGPT and scBERT. We omitted Geneformer because it requires pre-training weights as input. It can be seen that pre-training always improved results for scGPT, especially for the cross-dataset conditions. However, there was a little benefit of pre-training for scBERT. For both cases, freezing the pre-training layers and preventing them from being involved in the fine-tuning process was not recommended. In some cases, the finetuning performance of such freezing was worse than training from scratch. Transfer learning for different species is possible because, for the MCA dataset, pre-training based on human data can help predict cell types for the mouse. For the same type of GPU, the training process of scGPT was faster than scBERT and Geneformer, with more GPU memory usage, according to Extended Data Figure 16 (b).

In Extended Data Figures 17 (a) and (b), we explored the capacity of using cell embeddings from scGPT and Geneformer to annotate cell types (the zero-shot learning mode) and compared the results with the fine-tuning mode. We found that on average the fine-tuning mode surpassed the zero-shot learning mode for annotating cell types (p-value=1.9e-3 for scGPT, p-value=1.9e-3 for Geneformer, based on the Wilcoxon rank-sum test). Therefore, it is still necessary to consider fine-tuning scFMs for celltype annotation tasks.

In Extended Data Figure 18, we froze the front layers or scGPT (from 0 to 11) to investigate the relation between the number of freezing layers and annotation accuracy. We found that freezing layers had comparable or even slightly better performance than full-parameter fine-tuning. Although the metrics under different numbers of freezing layers are close and high, we still found that freezing five layers can lead the best performance. Therefore, we can freeze parts of layers for cell-type annotation based on single-cell FMs.

In Extended Data Figure 19, we show the performance of CellPLM, Geneformer, scGPT, and scBERT based on different optimizers across four datasets. Overall, Adam, AdamW, and Lion were comparable. Sophia-G was worse than them but better than SGD, and they were both unstable. Moreover, Geneformer did not support Lion and Sophia-G as optimizers, and thus the optimizers in Adam family are more preferred in fine-tuning single-cell foundation models.

Moreover, we explored the contribution of different loss function components towards the cell-type Annotation task based on ablation tests, and the results are shown in Figure 2 (d). Here we included three extra metrics, and details can be found in Appendix D. Based on the Accuracy of Figure 2 (d), the inclusion of mask loss is important. Moreover, the default setting is generally good across different tasks. Based on precision and recall of Figure 2 (d), the effect of different loss function terms had less effect on precision and more effect on recall. Such a difference could affect the final F1 score. Removing the GEPC loss function terms improved the cell type prediction for the DC, MHSP, and MB Spatial datasets, and did not affect the prediction performance for the other datasets.

Therefore, most single-cell FMs can handle the cell-type annotation task with suitable pre-training data and model structure, but investigation for the pre-training framework is needed to understand their specific performance differences across all the datasets.

### Evaluation based on gene-perspective tasks shows that single-cell FMs can handle tasks related to functions of genes

#### Gene Function Prediction

For this task, we intend to predict the functions of genes. We considered Geneformer, scGPT, scFoundation, and *vanilla* NN based on raw expression data and *vanilla* NN based on Gene2vec [59] for this task. scFoundation met running errors while generating the gene embeddings for datasets used in this task. We splitted the genes for training/testing and evaluated the results with the same four metrics as the Cell-type Annotation task because they are both classification problems. The results are shown in Figure 3 (a). On average, Geneformer and scGPT performed well in this task. Moreover, the accuracy scores of scGPT and *vanilla* NN based on Gene2vec were comparable, while there was a performance gap between single-cell FMs and *Vanilla* NN based on raw data. Therefore, using initial gene embeddings with prior information is meaningful for single-cell FMs.

**Fig. 3:**
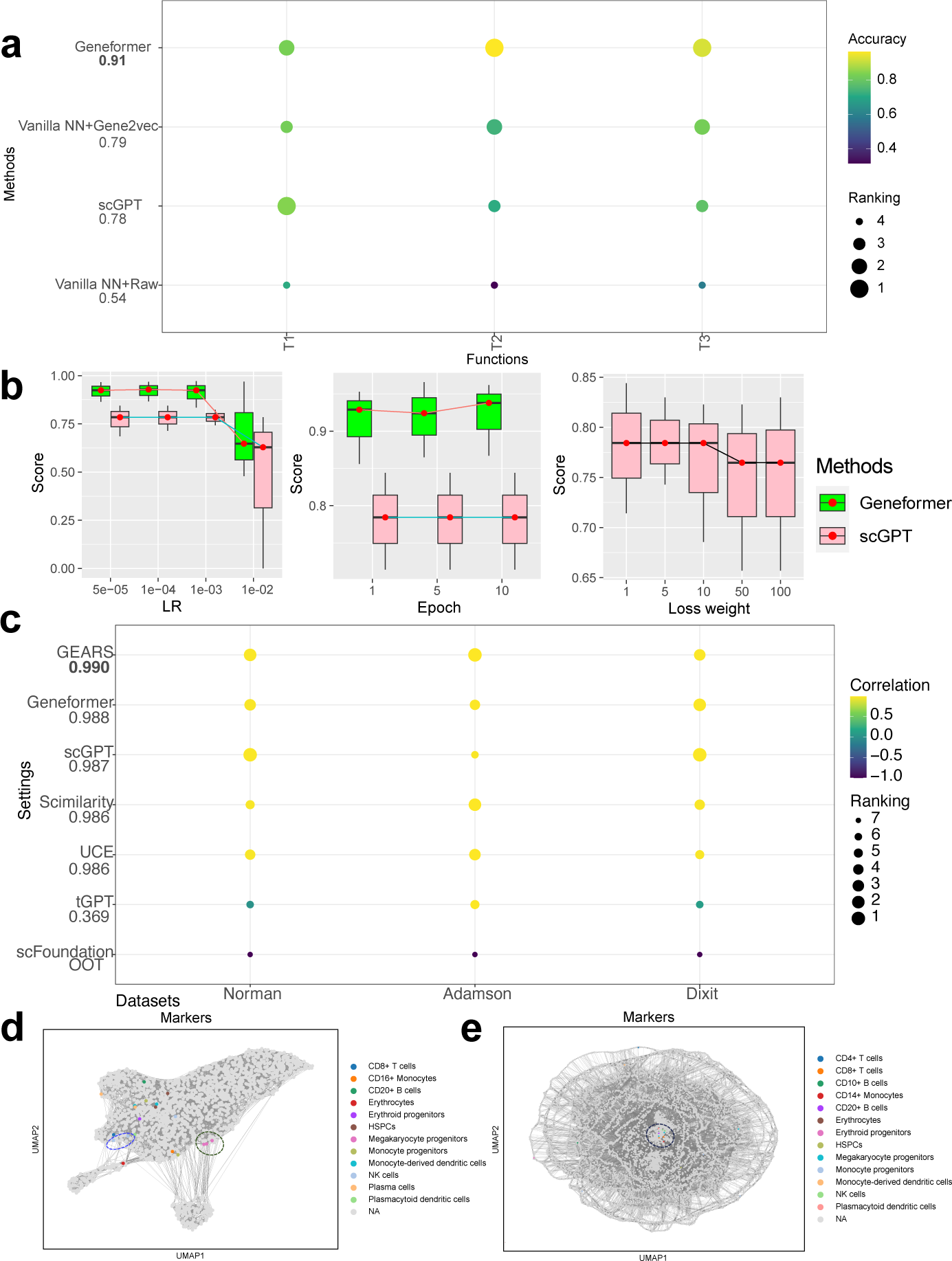
Experimental results of single-cell FMs and benchmarking methods for genelevel tasks. (a): Comparisons among Geneformer, scGPT, and *vanilla* NN in the Gene Function Prediction task. (b): The effect of hyper-parameters including Loss weight, Bins, and Learning rate for scGPT and Geneformer in the Gene Function Prediction task. (c): Correlation metrics of different methods. A higher correlation means lower rank and better performance. The numbers corresponding to the settings represent the average value across two datasets. (d): Dataset-level gene embeddings from scGPT colored by the marker genes of different cell types. (e): Dataset-level gene embeddings from Geneformer colored by the marker genes of different cell types.

Figure 3 (b) and Extended Data Figure 20 show the accuracy of different hyperparameter settings. Smaller learning rate and loss weight tended to have more accurate results. Geneformer was more sensitive to Epoch compared to scGPT. For scGPT, pretraining contributed more than fine-tuning in this task as increasing epochs did not affect the model performance. Only tuning the number of bins, mask ratio, dropout rate, and ECS threshold did not affect the prediction results.

In Extended Data Figure 21 (a), we considered different initial settings for model training. It can be seen that pre-training always improved performances of scGPT. Meanwhile, freezing the whole model did not affect the performance of scGPT.

Extended Data Figure 21 (b) shows the performance of scGPT based on different optimizers. Adam and AdamW were comparable, while Lion was worse than them but better than SGD and Sophia-G.

Extended Data Figure 21 (c) shows the ablation test results of scGPT for this task. There was no significant difference by comparing the default setting and those without certain components. Therefore, the task-specific loss function is the most important design for this task.

#### Perturbation Prediction

For this task, we intend to predict the gene expression levels under perturbations. We considered scGPT, Geneformer, tGPT, SCimilarity, UCE, scFoundation, and GEARS [60] for this task. We implemented a linear-regression-based model to predict perturbed gene expression levels from cell embeddings for those foundation models without this function. We used Mean Pearson Correlation (MPC) and Mean Squared Errors (MSE) as the metric to evaluate the performances across all genes, and details can be found in Appendix D. Therefore, we can assess the prediction performance at the gene level for both perturbed genes and unperturbed genes. The datasets include two perturbation conditions: a single-gene perturbation and a double-gene perturbation. Based on our experiments shown in Figure 3 (c) and Extended Data Figure 22, most of the foundation models (except tGPT) and GEARS were comparable in this task. Moreover, the running time of scFoundation for this task was too long to finish, which suggests the challenges of deploying FMs in solving this task. Since GEARS has robust performance, the requirement of developing FMs for handling such task is not clear.

Extended Data Figure 23 also summarizes results for different initial settings of scGPT for the Norman, Adamson, and Dixit datasets. The default setting performed best for these datasets across different settings. This indicates that the initial configuration of scGPT works well for this task. The performance was comparable between training from scratch and training from pre-trained weights. Freezing the decoder part of the model performed better than freezing the encoder part, which implied that the encoder part is important for perturbation prediction. Interestingly, the ability of zero-shot learning of scGPT towards this task was not suitable. Therefore, the contribution of using non-perturbed data as pre-training datasets for perturbation prediction related to perturb-seq datasets is not significant. We may need to investigate more types of perturbation, including drug-level conditions or disease-level conditions, to check the importance of introducing prior information for perturbation analysis.

Regarding the effect of hyper-parameters, Extended Data Figure 24 shows that scGPT is sensitive to adjusting the learning rate and epochs. Decreasing learning rate and increasing the number of epochs improved MPC. Higher learning rate caused a running error for scGPT. We did not identify patterns for other hyper-parameters. Extended Data Figure 25 (a) shows that AdamW, Sophia-G, Adam, and Lion had comparable performance for scGPT in perturbation prediction. SGD could significantly reduce the performance of scGPT. Therefore, the use of single-cell FMs for perturbation prediction does not require a complicated design.

#### Gene Network Analysis

For this task, we intend to evaluate the quality of gene networks inferred from gene embeddings. We considered scGPT, Geneformer, and scFoundation in this section. Firstly, we clarified the difference between Gene Regulatory Networks (GRNs) and Gene Co-expression Networks (GCNs), and the networks we evaluated here were GCNs. In general, GRNs characterize causal relationships among genes. The GRNs defined in scGPT are in fact GCNs, because the construction process is based on embedding similarity. For the inference of GCNs, two types of them can be defined: Type 1 GCN (Tissue-specific GCN) and Type 2 GCN (Cell-type specific GCN). We used the Immune Human Atlas dataset to evaluate the performance of inferring these two types of GCNs. The known information including marker genes [61], cell types [61], and Reactome pathways [62] was utilized to evaluate the performance of scGPT on the GCN inferences.

Type 1 GCN is generated by applying the single-cell FM to the entire dataset under a zero-shot learning framework to create gene embeddings. The similarity is then computed to infer gene-gene relationships based on these embeddings. The quality of the GCN is evaluated based on the relationships between marker genes for different cell types.

Type 2 GCN is created by applying the single-cell FM to generate cell-type-specific gene embeddings under the zero-shot learning framework, and the same type of similarity is used to infer gene-gene relationships based on these embeddings [36]. The quality of this GCN is evaluated based on the Gene Ontology Enrichment Analysis (GOEA) [63] for specific gene sets. These GCNs can provide valuable insights into the understanding of gene interactions and regulation in specific tissues or cell types, which could have broad applications in biology and medicine [64, 65].

In the analysis of the Immune Human Atlas dataset, we initially focused on Type 1 GCN, with the results presented in Figures 3 (d) and (e). The neighboring relationships within this dataset are colored according to the distribution of marker genes. We collected marker genes based on the source paper of this dataset [61] and filtered them to have marker genes with similar expression patterns for each cell type. As the gene-gene relationship was determined based on k-nearest neighbors, it can be viewed as a form of gene co-expression relationship. From Figure 3 (d), only marker genes from two cell types showed the co-embedded and isolated relationship. They are Monocyte-derived dendritic cells and Megakaryocyte progenitors. Figure 3 (e) shows that the embeddings from Geneformer for maker genes were all co-embedded. Therefore, the embeddings of scGPT are better than the embeddings of Geneformer on preserving the cell-type-specific information. Extended Data Figures 26 (a) and (b), on the other hand, represent the cluster labels based on the Leiden clustering method [66]. These clusters can be interpreted as groups of genes that share common functions, or “gene co-function clusters”. We first analyzed the cluster information from the gene embeddings from scGPT. For marker genes from other cell types, some of them are in different clusters shown in these two figures, and some genes are coembedded with other cell types’ marker genes. There are two isolated groups (9 and 12), but no marker genes are identified in these two groups. For the gene embeddings from Geneformer, we found that most of the marker genes from different cell types are embedded in the same cluster, while the rest of clusters did not contain much cell-type-related information. Therefore, the clustering result of scGPT was also more biologically meaningful than Geneformer, possibly due to the number of selected genes. Geneformer generated all genes for embeddings by default, while scGPT only focused on highly variable genes.

We also evaluated and explored GCNs based on the human immunology system quantitatively, which is known for its complexity due to interactions among various cell types. The original analysis of GCN from [36] focused on HLA genes and CD genes. We did not analyze HLA genes because they are highly polymorphic and thus carry a higher risk for having errors of reads [67–70], so the network result may not be reliable. Therefore, we selected genes co-embedded with CD3 genes, and utilized the embeddings of these genes to compute the GCNs. The results of scGPT and Geneformer for CD3-related genes are shown in Extended Data Figures 26 (g) and (h). The value above the edges represents the strength of correlation. The threshold of strong correlation here is set as 0.4. A publicly accessible database containing pathway information-the Immune System R-HSA-168256 pathway from the Reactome 2022 database [62] -was used as a reference for validation. The GCN was constructed based on the correlation between the two genes. For the CD3-related genes, there was nearly no overlap (7/1943) for scGPT, Geneformer (8/1943), and scFoundation (0/1943), indicating poor inference of GCN. We also visualize the detailed pathway information from scGPT and Geneformer in Extended Data Figures 27 (a) and (b), and we find that there is no overlap between the results from the two models. Moreover, for all the pathways involved in the selected gene set, we computed the ratio between the number of significant pathways and all pathways, and recorded the results in Extended Data Figure 26 (f). We found that gene embeddings from scGPT had a higher ratio, suggesting a better quality of scGPT output.

These analyses highlight how scGPT can be used to create gene embeddings to reveal important patterns of co-expression and co-functionality between different genes, offering insights into potential gene-gene interactions and regulatory relationships.

Extended Data Figures 26 (c) and (d) focus on gene embeddings categorized by cell types from scGPT, while Extended Data Figure 26 (e) shows the cell-type-specific gene embeddings from Geneformer. By treating cell types as observed labels, we also computed Normalized Mutual Information (NMI) and Adjusted Rand Index (ARI) for gene embeddings from different methods. Extended Data Figures 26 (c) and (d) show that gene embeddings of scGPT (NMI=0.049, ARI=0.035) from different cell types tended to be co-embedded and there was no apparent difference. The distribution of the remaining genes on the UMAP results was relatively random, and we also observed a random distribution of gene embeddings from Geneformer (NMI=0, ARI=0). Furthermore, the NMI and ARI scores of two types of gene embeddings are also low. One reason could be that the quality of gene embeddings was unsatisfactory. Since scGPT and Geneformer adopt the zero-shot learning scheme embeddings, it does not incorporate cell-type-specific information for a specific dataset. The other reason might be that the complex biological network in the human immune system makes the communication between cell-cell or gene-gene difficult to decompose [71, 72]. Additional analysis is needed for generating gene embeddings. This analysis illustrates the complexity of cellular functionality and the difficulty of clearly defining these relationships based on gene embeddings. Despite these challenges, the scGPT model still demonstrates its potential in identifying functional similarities between different cell types.

In conclusion, our results highlight the importance of critical evaluation and cross-referencing in the development of GCNs inference, as well as the potential and limitations of using single-cell FMs for this purpose.

### Evaluation based on Imputation and Simulation Analysis shows that further improvement of single-cell FMs is needed

#### Imputation

For this task, we intend to impute gene expression profiles of different datasets. We considered imputation for two different types of datasets, known as scRNA-seq datasets and spatial transcriptomic datasets. To impute scRNA-seq datasets, we intend to evaluate the performances of scFMs for filling technical zeros, which contain ∼ 20,000 genes. To impute spatial transcriptomics, we intend to evaluate the performances of scFMs for filling ∼ 20,000 unobserved genes, which are observed in scRNA-seq data. We compared scGPT and Tangram [73] in this task. The evaluation metrics for this task based on clustering and correlation are provided in Appendix D. The evaluation of clustering performance and gene expression correlation can assess the preservation of biological variation after imputing. The imputation results for scRNA-seq are summarized in Extended Data Figure 29 (a), which implies that the imputation function of scGPT for scRNA-seq data introduced more noise into the original sequencing data, suggesting the unreliability of the decoder’s output. According to Extended Data Figure 29 (b), scGPT performed well in the spatial transcriptomic data imputation task compared to the SOTA spatial imputation method, Tangram [73, 74]. Based on the evaluation of correlation and significance proportion, the imputation results of scGPT are better than the results of Tangram. Moreover, the scores of these two metrics based on the zero-shot learning version were even better than the pre-training version with scRNA-seq data. However, based on the results of the average bio score evaluation, the raw data had better scores. This could be caused by the sources of the spatial clustering labels, which were generated from gene expression clusters rather than expert annotation. Such methods could introduce bias before and after imputation.

Extended Data Figure 30 shows the results of Differentially Expressed Genes (DEGs) based on pre-imputation data and post-imputation data. Results in Extended Data Figure 30 (a) showed that scRNA-seq imputation was not reliable because the expression patterns of all genes were similar after imputation based on scGPT. However, based on Extended Data Figure 30 (b), no mitochondrial (MT) genes were included in the DEGs after imputation based on scGPT. These genes were identified as DEGs in the raw dataset. A high proportion of MT genes is indicative of low-quality data, which means cells with such patterns are apoptotic or lysing [75]. Therefore, the MT genes should be omitted in the downstream analysis by filtering[76, 77]. Moreover, based on Extended Data Figures 30 (b) and (c), the DEG patterns after imputation based on scGPT and Tangram are similar. Thus, scGPT has the potential to produce biologically meaningful imputation for spatial transcriptomic datasets.

#### Simulation Analysis

For this task, we intend to simulate synthetic scRNA-seq datasets. We considered scGPT, Splatter [78], and scDesign3 [79] for this task. We evaluated the output of scGPT against the output of scDesign3 and Splatter. For conditions incorporating batch effects, we employed the same metrics used in the evaluation of batch effect correction. In scenarios without batch effects, our metrics are primarily focused on assessing the preservation of biological information. As shown in Extended Data Figures 29 (c)-(e), scDesign3 outperformed scGPT and Splatter across two conditions of the simulation task. However, scGPT performed better than Splatter for simulating datasets without batch effect. In particular, scDesign3 had a more pronounced superiority in generating simulation data without batch effects, in comparison to scGPT. This is consistent with the results shown in Extended Data Figures 29 (d), (e) and Extended Data Figure 31. The gene-gene correlation from scDesign3 was also more similar to the gene-gene correlation of the raw data. The gene-gene correlation from scGPT has null value due to missing the gene expression certain genes, which is a problem caused by the decoder outputs from scGPT. Therefore, the simulation task needs to be improved for single-cell FMs as reference-based simulators. In addition, we present UMAPs of the output produced by different methods in Extended Data Figures 32 and 33 that illustrate the advantage of scDesign3. The embeddings of scGPT with the no batch effect settings tended to preserve the batch effect, while the embeddings with batch effect tended to remove the batch effect. The embeddings of Splatter with no batch effect also mixed different cell-type information.

### Model scaling contributes to single-cell FMs but their stability need to be improved

#### Model Scaling Analysis

In this section, we explored the contributions of model scaling based on CellPLM, scBERT, Geneformer, and scGPT. These methods have finetuned versions and do not suffer from running errors. The scaling law implies that models with large-scale parameter size tend to have better performances for certain tasks [80–82]. Here we considered three scenarios to investigate the model scaling: cross-data cell type prediction, cross-species analysis, and spatial transcriptomic analysis.

Inspired by [37], we compared models with large parameter size of pre-training to models of small parameter size. For the task of cross-data cell type prediction, we compared other scFMs with *Vanilla* neural networks (NNs) to identify any contributions of model scaling. For the task of cross-species analysis, we compared other scFMs with expert model SATURN [83]. As for the task of spatial data batch effect correction, we examined the statistics derived from our correction evaluation to verify our hypothesis and the expert methods include ResPAN and Novae [84]. Novae is a foundation model pre-trained with spatial transcriptomics data and thus it is a meaningful baseline to evaluate the contribution of transferring knowledge from single-cell transcriptomics to analyze spatial transcriptomics.

In the first scenario, our anticipated improvement from model scaling would be a significant improvement in prediction accuracy for single-cell FMs compared to *vanilla* NNs of a smaller size. Figure 4 (a) provides an overview of different model sizes. From Figure 4 (b), it is evident that scGPT and Geneformer outperformed *vanilla* NNs in terms of accuracy in cross-data scenarios, suggesting the contribution of model scale in the cross-data cell-type annotation task occurred for models which contain parameters larger than 10 million. The low accuracy of scBERT may also be caused by its default fine-tuning setting, pre-trained model weights, or pre-training datasets.

**Fig. 4:**
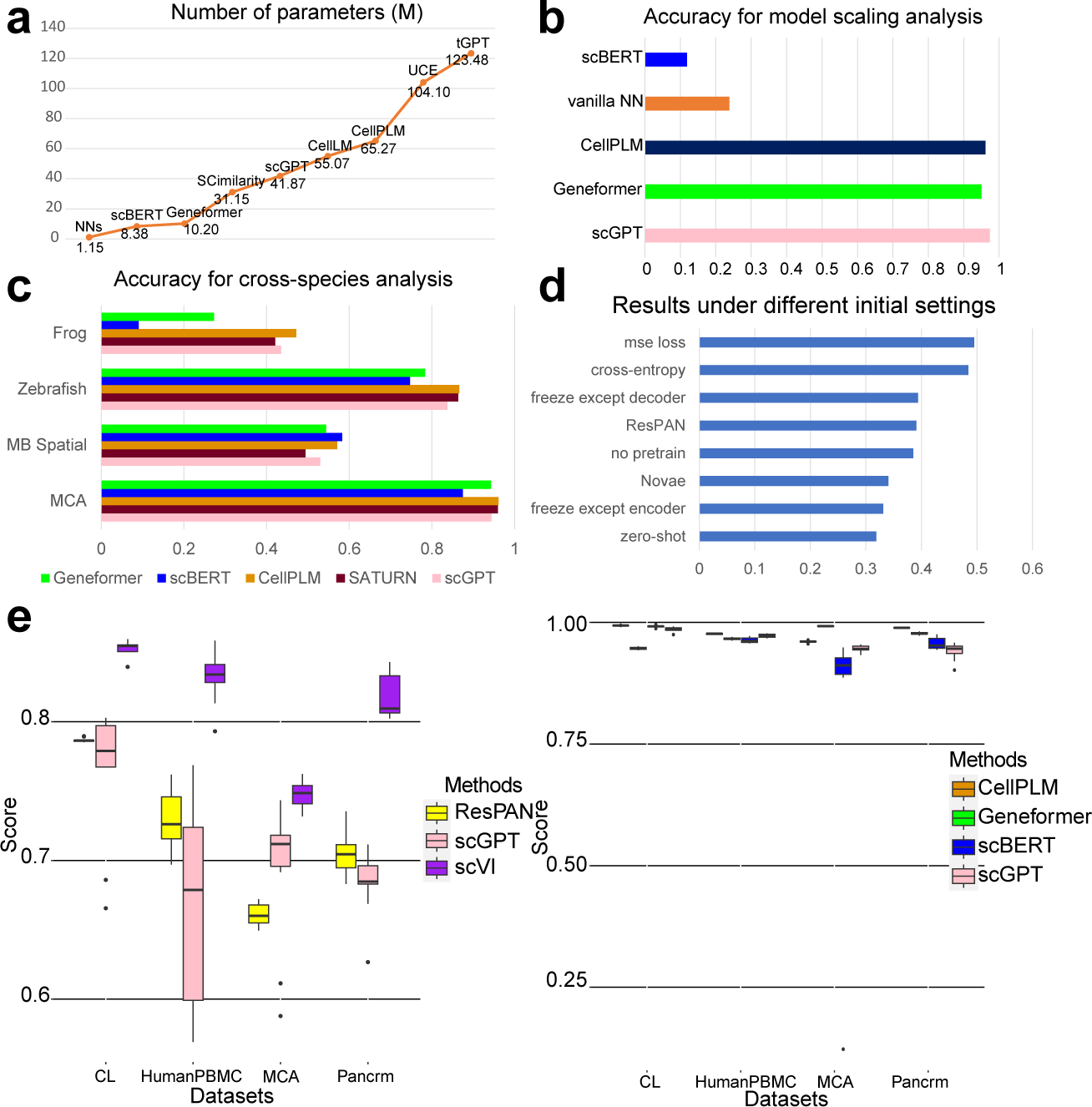
Different comparison groups for model scaling analysis and stability analysis. (a): The model scale of different methods, defined by the number of parameters (unit: Million). (b): Accuracy of FMs and *vanilla* NN in Cell-type Annotation task. The dataset here is the Pancreas cross dataset. (c): Accuracy of FMs and SATURN in Cell-type Annotation task across species. (d): Overall score comparison including ResPAN, Novae and different settings of scGPT. The dataset here is the human spatial transcriptomic dataset. (e): Different batch correction scores of different models based on changing random seeds (left) and different average classification scores of different models based on changing random seeds (right). The bold black line represents the median value, while the length of each box can be interpreted as the variance level.

In the second scenario, the desired contribution of model scaling mirrors that of the first task. Nonetheless, there was no enhancement in performance from Figure 4 (c). The cross-species cell-type annotation task is difficult, also suggested by [83, 85] as a representative task in analyzing cross-species information transition. Figure 4 (c) shows that SATURN is a strong baseline and it outperforms scFMs such as Geneformer and scBERT, while is comparable with scGPT. However, CellPLM can still outperform SATURN in most of the datasets used for cross-species analysis. Therefore, single-cell foundation models including CellPLM and scGPT showed advantages in this task.

In the third scenario, we considered two possible tasks. Firstly, in line with the batch effect correction task, we postulated that using Human scRNA-seq data for pretraining could aid in batch effect correction for human spatial transcriptomic data. Secondly, resonating with the cell-type annotation task, we hypothesized that pretraining on Human scRNA-seq data might assist in cell-type annotation for mouse spatial transcriptomic data. Figure 4 (c) shows that we did not detect any contributions of model scaling in the MB Spatial data annotation task. Figure 4 (d) suggests the contribution of model scaling for batch effect correction. scGPT under different loss functions all outperform Novae, while Novae performs better than the zero-shot mode of scGPT in batch effect correction. Therefore, including single-cell transcriptomics and spatial transcriptomics data jointly for model training can contribute to spatial transcriptomics analysis in batch effect correction. The fine-tuning process appeared beneficial in reducing the batch effect inherent in the spatial data, whereas scenarios with model freezing except decoder yielded subpar results. The performance of scGPT in the integration of spatial data was better than that of ResPAN. Therefore, we observed the contribution of model scaling in the batch effect correction for spatial transcriptomic data.

#### Stability Analysis

To analyze the stability of single-cell FMs, we selected Batch Effect Correction and Cell-type Annotation as two representative tasks and varied the seeds from 0 to 9 of single-cell FMs to investigate the model stability. All of the methods used in this section are fixed with their default hyper-parameters. These two tasks are the main tasks in single-cell data analysis and have solid metrics for evaluation. Ideally, the results of different single-cell FMs should not vary substantially across different datasets. We also considered stability for other benchmarking tools. Our experiment results summarized in Figure 4 (e) showed that the stability for singlecell FMs is task-specific.

Based on Figure 4 (e), the variance of scVI [86] and ResPAN was generally lower than that of scGPT, and scVI and ResPAN also had a higher score on average. Therefore, single-cell FMs were not as stable as SOTA deep-learning-based methods for the Batch Effect Correction task.

Figure 4 (f) suggests that the variance of Geneformer and CellPLM was generally smaller than scGPT and scBERT. All four models had high median average scores. Moreover, the variance of scBERT was relatively large in the experiments based on the MCA dataset, which implies that single-cell FMs might fail under certain random seeds.

## 3 Discussion

In this paper, we have evaluated the performance of ten single-cell FMs on eight different tasks for single-cell data analysis. The ranks of these single-cell FMs are shown in Tables 1 (a) and (b), where we not only considered the broader functions of the models, but also their usability. Based on our evaluation results, open-source models have higher ranks than closed-source models. Open-source models with tutorials are more friendly to researchers, and these models also receive a number of stars and likes based on Extended Data Figure 34. Tutorials associated with these models also enhance their accessibility to researchers.

Based on our experimental results, the performance and contribution of single-cell FMs are task-specific. For the Batch Effect Correction and Multi-omic Data Integration tasks, single-cell FMs did not outperform task-specific SOTA methods. Moreover, the correction results of atlas-level datasets were worse than those of the raw datasets. For the Cell-type Annotation task, we found that CellPLM, scGPT, and Geneformer had comparable performance. Moreover, these methods were better than the other models in this task. For the Gene Function Prediction task, Geneformer also performed well and better than classical models for prediction. We also showed that the pre-trained model can have better performance in this task. For the Perturbation Prediction task, the function of model pre-training is not apparent, and GEARS is comparable with other single-cell FMs in this task, questioning the need of the necessity for developing single-cell FMs to understand perturbations. For the Gene Network Analysis task, we did not observe the great contribution of gene embeddings from single-cell FMs for either tissue-specific or cell-type-specific analysis. Constructing the cell-type-specific gene networks is still challenging for scFMs reflected by the low clustering scores, and thus it is meaningful to explore methods for improving the function of network discovery. For the Simulation task, scGPT also did not perform very well. scDesign3, as a task-specific method, is better for simulating reference-based scRNAseq datasets. For the Imputation task, we showed that scGPT did not perform well in the scRNA-seq imputation task, but it outperformed Tangram in the spatial transcriptomic data imputation task both under zero-shot learning and fine-tuning frameworks. Such a finding suggests that single-cell FMs can transfer knowledge across different omic data. Furthermore, we observed the contributions of large-scale parameters for single-cell FMs in specific tasks. For the cross-data and cross-species Cell-type Annotation tasks, the performance of single-cell FMs is much better than the performance of baselines with smaller parameter sizes. Moreover, scGPT can also be used to analyze spatial transcriptomic data for batch effect correction, which has attracted much attention in recent years [87]. In our stability analysis, we found that single-cell FMs were not very stable in the Batch Effect Correction task. Moreover, in the evaluation for the Cell-type Annotation task, we found that different single-cell FMs had different variance levels. The variance under different random seeds was driven by datasets. Considering the problems of stability, we still have difficulty refining it into a toolbox (for example, Seurat [88] or Scanpy [77]) with various functions.

Therefore, by considering the definition of a foundation model, there are evident limitations in their construction and training steps. A more comprehensive understanding of such FMs is necessary to address these issues. In this manuscript, we tried different approaches to discover insights from model components and training settings to possibly enhance the performances of single-cell FMs. Our conclusions, combined with the recommendations of models for different tasks, are summarized in Table 2. Considering the difference in resources required for zero-shot learning and fine-tuning, as well as the difference in model effectiveness, we suggest trying to apply the model’s zero-shot learning mode to obtain the embeddings required for the task at first, and then consider fine-tuning model for the given task if the performance of zero-shot learning mode is not satisfactory and we can obtain a good task-specific loss function. We also explored more advanced parameter-efficient fine-tuning (PEFT) frameworks such as LoRA [89] for scGPT in Batch Effect Correction and Cell-type Annotation. Extended Data Figure 35 shows that LoRA slowed down the fine-tuning process and did not improve related scores. Thus, the training of single-cell FMs is more nuanced than that of general FMs. While it is important to consider the similarities between these two model types, we must also consider the differences rooted in the datasets and domain-specific knowledge.

**Tab. 2:** Insights from the benchmarking results of scEval. We organize the table by tasks and model settings.

| Topic | Summary | Recommendation |
| --- | --- | --- |
| Batch Effect Correction | Overall, all scFMs performed worse than task-specific method. | Selecting task-specific method (Harmony) as a starting point. |
| Multi-omic Data Integration | Overall, all scFMs performed worse than task-specific methods. | Selecting task-specific method (GLUE) as a starting point, and including multimodal information in the pre-training stage of scFMs. |
| Cell-type Annotation | Overall, finetuned scFMs performed well in this task. | Exploring scGPT, CellPLM and Geneformer for annotating cell types, and we encourage researchers to include cell states in the pre-training stage. |
| Gene Function Prediction | Overall, finetuned scFMs performed well in this task. | Exploring Geneformer for predicting gene functions. |
| Perturbation Prediction | Overall, the performances of scFMs are close to task-specific methods. | Selecting task-specific method as a starting point, and including perturbation information in the pre-training stage of scFMs. Moreover, exploring the prediction based on zero-shot cell embeddings is also attractive. |
| Gene Network Analysis | scFMs are not good enough for modelling gene interaction networks. | Starting from gene expression profiles rather than gene embeddings for network inference and analysis. |
| Imputation | scFMs help on imputing spatial transcriptomics, while do not perform well in imputing scRNA-seq data. | Exploring the ability of scGPT for imputing spatial transcriptomics with a reference-free design. |
| Simulation | scFMs performed worse than task-specific methods for simulating scRNA-seq datasets. | Selecting task-specific method (scDesign3) as a starting point. |
| Scaling Law | scFMs followed scaling law for cell-type annotation. | Investigating the setting of biological questions before enlarging model size. |
| Stability | scFMs performed inconsistently in batch effect correction, while performed consistently well in cell-type annotation. | Exploring different strategies for pre-training scFMs, and developing functions from cell embeddings or gene embeddings. |
| Training Strategies | Smaller learning rate (e.g. $1e-4$) can improve model performance. During the finetuning process, freezing part of models can also improve efficiency. | Using small learning rate and exploring Parameter-Efficient Finetuning methods. |
| Optimizers | Optimizers in Adam family perform better based on our evaluation. | Using Adam or AdamW for model training. |
| Hard-Ware Requirement | We need at least one GPU with over 40 GB MEM for finetuning. | Using NVIDIA A100/A40/A6000 for deployment. |
| Model Development | scFMs vary greatly in the extent to which they are open source and the number of features they have by default. | Investigating scGPT, Geneformer and CellPLM for scFM development and deployment. |

The application of FMs to single-cell data remains a promising avenue of exploration, given its impressive performance in prediction tasks. Since much has been done to optimize general FMs [90–93] (including efficient tuning, model compression, and other research directions), we focus specifically on how to better train and apply single-cell FMs. Here, we discuss several future directions:

In terms of pre-training preparation, we need high-quality data spanning different contexts, such as various cell types, disease states, genders, and even data from different species for pre-training datasets. High-quality pre-training datasets are important for general FMs; otherwise, the performance may be reduced [94, 95]. The qualification of pre-training data can also be verified with the online learning framework [96]. We can evaluate the performance of single-cell FMs on downstream tasks and then decide which datasets to include. With a better pre-training design, we may increase the scale of current models to a billion level. Moreover, the incorporation of biological information is crucial for the success of a FM in biology. Therefore, it is possible to incorporate other biological factors, including GRN and cell-cell interactions (CCI) [26] in the modeling process. With such information, we can also develop domain-specific or tissue-specific FMs for single-cell data analysis. Including extra labels in the pre-training stage might also improve the model’s ability in identifying gene networks under specific context. Exploring the contribution of multimodal data to develop a multimodal FM is also a possible track. For example, incorporating text-based biological information [97] or multi-omic data with new tokens may help us further extend the functions of these FMs.

As for model training, both the pre-training and fine-tuning steps of the existing single-cell FMs need improvement. During model pre-training, we should focus on incorporating biological information or human feedback into the process, as opposed to just relying on the conventional masked token prediction task. For instance, integrating cell-type or disease condition information into the pre-training step is an intriguing approach, and it allows experts to evaluate the quality of model output during the training process, which is also meaningful. Moreover, we should also consider the security or trustworthiness of FM training [98]. Using poisoned training single-cell datasets (for example, wrong data or make-up data) as an attack can test the robustness of single-cell FMs. For model fine-tuning, instruction tuning [99] is a potential direction to explore. In this context, cells could be considered as prompts, as described in scGPT. Another possible direction is to focus on generating unified embeddings for cells/genes and combining them with task-specific models for downstream applications, inspired by [37, 100].

For model evaluations, we need more effective methods to assess results for certain tasks. Our recommendations for researchers to consider as evaluations in developing new single-cell FMs are summarized in Appendix C. For instance, we may include verification from biology experiments to avoid the harm of incorrect output of single-cell FMs (also known as FM hallucinations [101]). Moreover, analyzing the contributions of model scaling of single-cell FMs is also important to explore the breakthrough contribution and significance of a model, though we have not identified a task that can only be addressed by developing single-cell FMs in our evaluation. In addition, we should also account for the pre-training costs (such as training time and power consumption), and avoid developing a model that consumes large resources but has no significant improvement for downstream tasks.

When it comes to task selection, we should first define the tasks rigorously. For example, in the GRN inference task defined by scGPT, we cannot treat a co-expression network based on gene embeddings the same as a gene regulatory network. Also, from scGPT, using attention, we can infer gene-gene correlation strength with direction, but the relation between the correlation of features and the value in the attention map is in debate [102–104]. We should also consider more meaningful, hardcore, and challenging tasks related to single-cell and spatial data for single-cell FMs. These tasks typically need prior information from large-scale transcriptomic data. For instance, tasks such as in-silico treatment analysis [35], complex perturbation analysis in the single-cell levels [105], and spatial data imputation [106] are difficult to handle without prior information, thus provides an opportunity for FMs to make a difference.

In summary, our goal in studying single-cell data-based FMs is to develop a largescale model that is capable of performing multiple tasks with stable and reliable results. Such a model should also be user-friendly, with detailed tutorials and well-maintained websites. Our results indicate that there is much space for improving single-cell FMs. Although our evaluation is subject to the number of current single-cell FMs, the scale of current single-cell FMs, and the strategies of pre-training, we hope that our analysis can provide insights into the best practices and guide the development of future FMs for single-cell data analysis.

## 4 Methods

### Problem definition

We consider a pre-trained FM, denoted as *M*(*x, θ*), which is based on the single-cell dataset *D*. Here, *θ* embodies the set of both model parameters (e.g., network weights) and hyper-parameters (e.g., epochs and learning rate). Different FMs have used distinct pre-training datasets. The model structure for the fine-tuning phase is defined as *M*^′^(*x, θ*^′^). Our objective is to ascertain the optimal set of *θ*^′^ for various sub-tasks. Formally, we denote the loss for task *k* as *L_k_*(·, ·), and use the evaluation dataset *D*_eval_ = {*x_i_, y_i_*}*^n^_i =_* _1_, to assess *L_k_*. Our primary goal is to find:

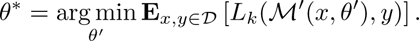

Our second goal is to evaluate the performance of different single-cell FMs, that is, we intend to find:

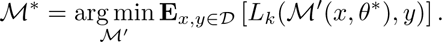

Our third goal is to assess other abilities of single-cell FMs, including: 1. zeroshot learning; 2. model scaling [80–82]; 3. cross-species data analysis; 4. biological mechanism exploration; and 5. stability.

### Parameters and tasks

Most single-cell FMs share the pre-training process. By considering the overlap among various single-cell FMs, we have selected scGPT, scBERT, and Geneformer as representative examples for our analysis. We also highlight the downstream tasks in Figure 1 (a). We focus on eight fine-tuning tasks in total: 1. Batch Effect Correction; 2. Multi-omic Data Integration; 3. Cell-type Annotation; 4. Gene Function Prediction; 5. Perturbation Prediction; 6. Gene Network Analysis; 7. Imputation; and 8. scRNA-seq Simulation. By compiling all the hyper-parameters across different models, we present a list of factors that can affect the model performance categorized by types in Figure 1 (b). The definition of different hyper-parameters is discussed in Appendix A. To analyze the effect of different hyper-parameters, initial settings, and optimizers, we selected representative datasets for different tasks. For Batch Effect Correction and Cell-type Annotation, we selected Pancrm, HumanPBMC, Cell Lines, and MCA because they cover various data types. Pancrm is from Pancreas tissue and has five batches. HumanPBMC is from PBMC and has nine cell types. Cell Lines has two cell types, as a binary label dataset. MCA is from *Mus musculus*. For Multi-omic Data Integration, we included all of the four datasets we used to analyze because their scales cover a large range. For Gene Function Prediction, we included all of the three datasets we used to analyze because they correspond to different sub-tasks. For Perturbation Prediction, we included all of the three datasets we used to analyze because their scales cover a large range and are from different sources.

### Explanations of scEval

Here we introduce a framework known as scEval to evaluate the performance of different single-cell FMs on various tasks. The whole pipeline is highlighted in Figure 1 (a). Since most of the single-cell FMs choose to reconstruct the masked gene expression levels in the pre-training stage, we can use an encoder-decoder structure (similar to an auto-encoder [107]) as well as extra finetuning blocks to unify their architectures. Moreover, we split the eight tasks into two types. For tasks included in sub-task 1, we need an extra fine-tuning block to generate the results. Cell-type Annotation and Gene Function Prediction belong to this task type. For tasks included in sub-task 2, we rely on the outputs of the encoder part and decoder part to generate the results. The rest of the six tasks belong to this task type. The idea of scEval is inspired by benchmarking analysis in both single-cell area [61, 106, 108] and LLM area [109, 110]. According to Figure 1 (b), for each task, we collect the output of single-cell FMs under different settings. Each FM has its specific running settings, and their default modes can be classified into fine-tuning-based models and zero-shot-learning-based models, thus we propose a framework focusing on the outputs of different single-cell FMs to unify the evaluation process. To analyze the factors affecting single-cell FMs, we also adjusted the factors of different FMs and collected their results for evaluation. Moreover, scEval contains different evaluation pipelines for different tasks, and it can also be easily extended to evaluate more tasks by adding more functions.

### Batch Effect Correction

Batch Effect Correction is an essential step following scRNA-seq data pre-processing. It primarily signifies the distribution disparity in scRNA-seq datasets originating from the same tissue, which can be attributed to various factors [111]. The reduction of batch effects is critical not only to allow researchers to discern genuine biological signals but also to facilitate integrated analyses across different studies. The challenge of this task arises from the need to balance the removal of batch signals with the preservation of biological signals. We treat this task as a data integration problem.

For the Batch Effect Correction, the metrics we considered here were inspired by scIB [61], including Normalized Mutual Information (NMI), Adjusted Rand Index (ARI), and Cell-type Average Silhouette Width (cell-type ASW) for the biological conservation score; and batch Average Silhouette Width (batch ASW), Principal Component Regression (PCR), Graph Connectivity (GC) and kBET [112] for the batch effect correction score. We compute the weighted average of these metrics to represent the final batch effect correction score. Details of these metrics can be found in Appendix D.1. Let *S_bio_* donate the average biological conservation metric and *S_batch_* donate the average batch effect correction metric as *S*_batch_, the final model score is:

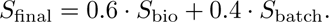

### Multi-omic Data Integration

Multi-omic Data Integration is a key step for multi-omic data analysis [22]. It is akin to an advanced form of batch effect correction. If unpaired multi-omic data are present, the objective is to map different datasets into a shared space for subsequent analysis. If paired multi-omic data are present, the goal is to assess whether the use of multi-omic data can contribute to learning a more comprehensive representation of the data. A significant challenge here is how to align omics at the feature level. For instance, the feature of the scRNA-seq data is a gene, the feature of the scATAC-seq data is a peak, and the feature of the protein data is a protein. The tokenization step can become complex given different modalities. We treat this task as a data integration problem. We used the same metrics for multi-omic data integration as those for Batch Effect Correction.

### Cell-type Annotation

Cell-type Annotation is another key step following singlecell data pre-processing [113]. This step annotates each cell with its accurate cell-type label, which can be achieved through prior knowledge [114] or computational methods [115]. These annotated cell-type labels can provide essential biological information for further downstream analysis, such as cell-type specific network analysis. In addition, drug response prediction [37] or single-cell disease classification [116] can also be treated as a variation of this task. A common approach employed by single-cell FMs in dealing with the cell-type annotation task is to use single-cell datasets for model training and treat the unannotated datasets as testing datasets. The challenge lies in predicting or annotating a set of cells that originate from studies different from the training datasets. Differently, SCimilarity pre-trained the model with cell-type labels to allow query of cell types directly for testing datasets. Moreover, the existence of cells with novel cell types (which are not included in the training datasets) further complicates the problem. We treat this task as a multi-label classification problem. Regarding the choices of metrics for evaluation, we also computed the Pearson correlation coefficients between accuracy and weighted F1-score for the top-tier methods in our evaluation (including scGPT, Geneformer, and CellPLM), the coefficients are all 0.99 with a very small p-value (p-value*<*2.2e-308). Therefore, accuracy is a suitable metric used in our comparison.

In this task, we chose datasets with batch effect in two different cases. The intradataset case allows batch intersection, which means that the training and testing datasets can contain cells from the same batch. Here the total dataset was split into approximately 70% as a training dataset and the rest as a testing dataset. The interdataset case is cross-batch (cross-data) annotation, which means that the training and testing datasets are from different sources. We consider two datasets from the same tissue in this setting. More specifically, we consider the Pancreas cross dataset from the Pancreas, and the Liver cross dataset from the Liver. The main score for evaluation here is accuracy, which is defined as:

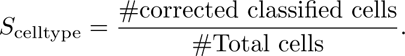

We also consider Precision, Recall and F1 scores in the analysis for ablation test, and details can be found in Appendix D.2. Moreover, except the general comparison, we considered four datasets for the effect of different hyper-parameters, initial settings and optimizers: Pancrm, HumanPBMC, Cell Lines, and MCA, which is from *Mus musculus* rather than *Homo sapiens*. Finally, we investigated the contribution of freezing layers for cell-type annotation by freezing different numbers (from 0 to 11) of forward layers of scGPT.

### Gene Function Prediction

Gene Function Prediction is important to identify the properties of genes across different conditions [35]. There are approximately 20,000 protein-encoding genes for humans [117] and only some are annotated with functions. Accurate prediction of gene function can help us understand and infer the role of genes in biological systems. Here we consider three types of functions for genes. The first one is dosage-sensitive or non-sensitive. Some genes are dosage-sensitive, which means that they are significant in the analysis of Copy Number Variants (CNVs) related to genetic diagnosis. The second one is Bivalent versus non-methylated. Bivalent chromatin structure is important to identify key developmental genes in embryonic stem cells (ESCs). Therefore, identifying bivalently marked genes versus unmethylated genes is important. The third one is Bivalent versus Lys4-only methylated. Lys4-only-methylated genes are also different from bivalently marked genes. We compare the model output with the true gene labels. We treat this task as a binary classification problem. Here we used the same metrics as the cell-type annotation task. We used a public dataset [35] considering only labeled genes in the dataset for prediction and evaluation.

### Perturbation Prediction

Perturbation Prediction [60] is a task based on gene editing and single-cell sequencing technologies. After silencing some genes, we can obtain unperturbed and perturbed gene expression levels by sequencing, which allows us to explore the interactions between genes. A well-known technique is Perturb-seq [118]. In perturbation prediction, we intend to predict the gene expression level after gene editing. Here a model may predict seen gene perturbation in the testing datasets (an easier one) or predict unseen gene perturbation in the testing datasets (a more difficult one). We treat this task as a regression problem. The metric we used here is MPC, and the details can be found in Appendix D.3. In the perturbation prediction task, we construct the paired input-target datasets by selecting the cells with non-control guide identity and then randomly sample cells under the control condition, and then combine them as the training and testing datasets. Our Perturb-seq datasets are from GEARS [60], which contain cells with three conditions: control; one gene perturbation; and two genes perturbation. In the evaluation process, we combined case 2 and case 3. We also compared the MPC across all genes to avoid bias in the gene selection process.

### Gene Network Analysis

Gene Network Analysis is a downstream task for single-cell datasets [119]. The objective is to infer specific gene networks (for example, Gene Regulatory Network (GRN) or Gene Co-expression Network (GCN)) from different datasets. A GRN can assist in understanding the regulatory relationships between genes and predicted perturbation outcomes. The challenge in this task stems from the Granger causal relationship or time-dependent correlation [120]. A GCN can be used to analyze genes with similar functions or uncover the characteristics of genes in some diseases [121]. GCN and GRN are two different tasks because correlation does not imply causal relation [122]. This limitation means that we cannot determine which genes are the “causes” of expression level changes in other genes only based on embeddings similarity or correlation. We treat this task as a network inference problem. In the gene network analysis task, we considered using the overlap between ground truth genes in certain pathways and inferred genes as one metric, and using the ratio of significant pathways related to inferred genes as the other metric. Details can be found in Appendix D.4.

### Imputation

Imputation is a filling task related to missing data. Generally, we have two targets: 1. Perform imputation for scRNA-seq data to reduce data noise and fill in technical zeros with biologically meaningful values [123, 124]. 2. Perform imputation for spatial transcriptomic data because of unseen or unmeasured genes [73, 74]. According to [125], current spatial imputation methods do not show strong performance across different datasets. Using single-cell FMs, we can either use zero-shot learning to impute the unseen genes, or fine-tune our model based on reference scRNA-seq with more genes to perform imputation. We treat this task as a matrix-completion problem. Details of the metrics we used here can be found in Appendix D.5. In the imputation task, we used two public datasets from the mouse tissue to analyze the performance of single-cell FMs. One dataset is a scRNA-seq dataset, and another one is a spatial transcriptomic dataset. For the imputation of the scRNA-seq dataset, we used the output of the model decoder as imputation results. To evaluate this task, we used the biology preservation score from the batch effect correction and compared it to the score from raw data. For the imputation of spatial transcriptomic data, we considered two different settings to perform imputation. The first setting uses scRNA-seq to perform fine-tuning and inference based on spatial transcriptomic data. The second setting uses a zero-shot learning framework to directly perform inference based on spatial transcriptomic data. We consider using the correlation between known raw gene expression and known imputed gene expression as a metric.

### Simulation

scRNA-seq Simulation is a data generation task. Leveraging the generative pre-training process of scGPT, we can generate gene expressions based on real datasets. Since a prevalent issue with scRNA-seq data simulation is the considerable divergence between simulation datasets and real datasets [78], direct generation from real datasets is preferred.

By arranging different sequences of masking genes or altering different seeds, we can generate new simulated scRNA-seq datasets from real ones. By modifying scGPT, we fine-tuned the baseline model based on the reference scRNA-seq dataset with reconstruction loss and utilized the outputs of the decoder part to generate the simulation datasets. Such simulation datasets do not have exactly the same gene expression profiles compared with the reference scRNA-seq dataset, but they should preserve the biological variation information from the reference dataset. We utilized scGPT to simulate datasets with batch effect or without batch effect. To simulate datasets with batch effect, we removed the loss function for removing batch effect in the original pipeline. To simulate datasets without batch effect, we kept the original pipeline for the batch effect correction task but utilized the outputs of the decoder rather than the encoder. We did not incorporate extra information for implementing this function. The quality of our simulation datasets can be evaluated by comparing them with the outputs of current simulation methods. We treat this task as a data-generation problem.

We used the same metrics as Batch Effect Correction for evaluation. It is possible to produce diverse reconstruction outcomes from a single real dataset by varying the random seeds. This feature enables us to create simulated single-cell datasets. Notably, these generated datasets retain the same gene sets as their input counterparts. We generate the simulation results by multiplying the output of the gene expression decoder and Bernoulli decoder. We have the flexibility to generate datasets either with or without batch effects. If we intend to produce datasets with batch effects, the gradient reverse loss function is omitted. Conversely, to generate datasets without batch effects, we either retain this function or utilize single-batch data.

### Model Scaling

[80–82] argue that due to the extensive number of parameters, FMs can manage specific tasks that small-scale models cannot handle. Such an attribute refers to the scaling law. We hypothesize that single-cell FMs may also possess this capability. To test this, we devise different scenarios similar to instances of model scaling to assess the performance of single-cell FMs. Such scenarios include: Cross-dataset Cell-type Annotation, Cross-species Cell-type Annotation, and Spatial Transcriptomic Data Analysis.

### Ablation tests

Given that there is no existing work investigating the significance of different loss function components of single-cell FMs, we conducted a comprehensive analysis of the impact of various loss function components. These include the Masked Gene Expression loss (Mask Loss), Zero Log Probability Loss (Prob Loss), Gene Expression Prediction for Cell Modelling Loss (GEPC Loss), and Elastic Cell Similarity Loss (ECS Loss). We retain the task-specific loss in the fine-tuning process as the baseline condition (All remove). Consequently, our null hypothesis (*H*_0_) is that the removal of the component *C* will not degrade the performance, while the alternative hypothesis (*H*_1_) is that the removal of component *C* will worsen the model’s performance. Comparing the score after eliminating a specific component to the score from the default setting allows us to determine whether to reject the null hypothesis. The test we employed here includes both the paired Students’ t-test and Wilcoxon Rank-sum test. The details of the different loss function components are shown below.

1. Mask Loss: In both pre-training and fine-tuning, we mask the expression levels of some genes, denoted as *M*_mask_, for gene expression prediction. The mask loss is motivated from this setting and works as follows,

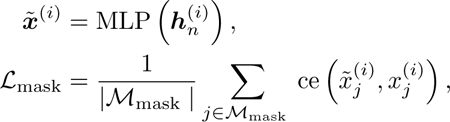

where 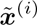 represents the predicted gene expression levels for cell *i*, while 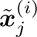 represents the ground truth. ***h****_n_*^(*i*)^ represents the embeddings for cell *i* with *n* genes. MLP means that we use linear Multi-Layer Perceptron (MLP) Neural Networks as the output of single-cell FMs under the setting of this loss function.

2. Prob Loss: Since single-cell data can be treated as count data, we can use the Bernoulli distribution to model the occurrence of the expression in masked genes and use the Maximum Likelihood Estimation (MLE) approach to estimate the parameters of Bernoulli distribution. Such loss function can be used to determine whether the given masked gene position carries zero expression levels or not. Prob Loss works as follows:

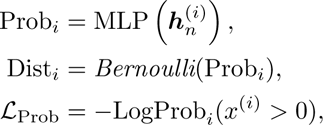

where Prob*_i_* represents the output of the model for the parameter estimation for cell *i*. The estimation is based on one MLP model. Dist*_i_* represents the Bernoulli distribution based on Prob*_i_*. LogProb*_i_* is the log probability based on the relationship between gene expression levels of cell *i* and zero, which can be computed based on Dist*_i_*.

3. GEPC Loss: This loss function is similar to Mask Loss, but now we predict the gene expression levels based on cell embeddings or cell representation. For cell *i* with gene *j*, we create a query vector ***q****_j_* and represent the cell based on ***h***^(*i*)^*c*. We can use the inner product between these two terms to predict gene expression levels. That is,

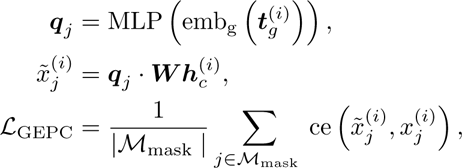

where emb _g_(***t****_g_*^(*i*)^) represents the embeddings of the gene token *g* in cell *i*. We also use one MLP to generate the query embeddings. The following process is similar to the steps for *L*_mask_.

4. ECS Loss: This loss function is used to control the similarity of the embeddings of cells in the same batch, which is defined as:

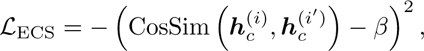

where CosSim represents the cosine similarity function, *i* and *i*^′^ are the indices of the two cells. The idea of this loss function is to ensure the similarity between paired cells is higher than the predefined threshold *β*. Moreover, dissimilar pairs should be more dissimilar, respectively.

### Task-specific fine-tuning process

For the experiments we have in the Results section, we load the pre-training weights based on the requirement of different single-cell FMs. The pre-training weights we used can be found in our GitHub folder. After the fine-tuning process, we recorded the related metrics and conducted more analysis. In the experiments for all tasks, we chose scGPT as a baseline and representative model for the following three reasons. Firstly, scGPT is an open-source single-cell FM with the largest datasets for pre-training, and it is well-defined with detailed tutorials. In addition, the architecture of scGPT is easy to adjust and includes multiple loss function components. The functions of Gene Function Prediction, Imputation, and Simulation for scGPT were designed in scEval for evaluation. The settings of hyper-parameters for these tasks are transferred from the design for cell-type annotation and batch effect correction tasks. Moreover, different single-cell FMs have overlaps and unique terms in the pre-training and fine-tuning framework, but scGPT is the most general one. In each task, we also included task-specific methods as comparisons. Moreover, for Geneformer, scBERT, CellPLM, and CellLM, we also evaluated their performance based on shared hyper-parameters or optimizers with scGPT to verify if our rules found in scGPT can be extended for other single-cell FMs.

## 5 Data availability and reproducibility

We used the resources from the Yale High Performance Center (Yale HPC) to conduct all of the experiments. Our maximum running time for each dataset is 24 hours. The version of GPU we used is NVIDIA A100 (40 GB). The random seed of all experiments is the same as the default setting of the original papers.

The information of datasets and the download link can be found in Appendix G. The codes of **scEval** can be found in https://github.com/HelloWorldLTY/scEval with MIT license.

## 6 Acknowledgements

We appreciate the comments, feedback, and model explanations from Yingxin Lin, Haotian Cui, Christina Theodoris, Rex Ying, Minsheng Hao, Xinming Tu and Malte D. Luecken. We also appreciate the constructive comments from reviewers and editors of this manuscript. This project is supported in part by NIH grants R01 GM134005 and P50CA196530.

## 7 Author contributions

T.L. designed the study with H.Z. and Y.W. T.L. ran the experiments with K.L. H.L. and T.L. designed the website. T.L., K.L. and H.Z. wrote the manuscript. H.Z. supervised this work.

## A Hyper-parameters

We first define different hyper-parameters used for hyper-parameters analysis.

1. Dropout: The dropout rate is the probability of making the neural unit inactive or closed. If a unit is closed, it will not participate in the training process but will take part in the inference process. The design of dropout is to avoid or reduce overfitting. The scope is [0, 0.1, 0.2, 0.3, 0.5].

2. Loss weight: Changing the weights of different loss function components is a well-known approach to increasing the performance of neural networks. Here the loss weight means the weight of the task-specific loss component. For example, for the Batch Effect Correction task, such weight represents the loss weight for the loss of the gradient reverse network. The scope is [1, 5, 10, 50, 100].

3. Bins: To reduce the batch effect during the pre-training or fine-tuning process in the pre-processing step, the gene expression is divided into different intervals and the endpoint values of the intervals are used to replace the original gene expression. The intervals here are the bins. Formally, the binned expression value *x_j_*^(*i*)^ for cell *i* is:

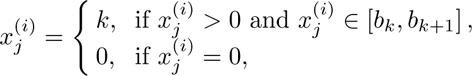

where *k* ∈ {1, 2*, …, B*} and *B* represents the number of bins. The scope is [51, 101, 151, 201, 501, 1001].

4. Mask ratio: In the pre-training and fine-tuning process, we choose to mask a fixed ratio of gene expression levels and try to use our decoder to reconstruct the expression levels of the masked genes. Such a ratio is defined as a mask ratio. The scope is [0.1, 0.3, 0.4, 0.5, 0.7, 0.9].

5. Epoch: Epoch represents the number of steps we use to train our model based on the whole training dataset. The scope is [1, 5, 10].

6. Learning rate: Learning rate is designed for the optimizer, it represents the step size of gradient descent. The scope is [5e-5, 1e-4, 1e-3, 1e-2].

7. ECS threshold: The threshold *β* in the ECS Loss represents the upper bound for the similarity between paired cells. The scope is [0, 0.2, 0.4, 0.6, 0.8, 1.0].

Details of the settings of hyper-parameters we used in the evaluation for all of the methods are summarized in Supplementary file 4.

## B Initial Settings

In this section, we delineate the initial settings for Batch Effect Correction, Multi-omic Data Integration, Cell-type Annotation, and Perturbation Prediction tasks.

1. Zero-shot learning: This refers to using our pre-trained model without any further training for unseen datasets on specific tasks. This setting aids in validating the significance of pre-training.

2. No pre-training: This setting entails starting the fine-tuning process without any pre-trained weights, allowing us to train a *de novo* single-cell FM. This configuration helps ascertain the importance of pre-training.

3. Cross-entropy (default setting): Here, we utilize the cross-entropy loss function for the reconstruction of gene expression levels, which is the default setting in scGPT.

4. Mean Squared Error (MSE) loss: In this setting, we employ the MSE loss function for the reconstruction of gene expression levels.

5. Freeze except encoder: This setting involves freezing the weights of the encoder component during the fine-tuning process.

6. Freeze except decoder: Here, we freeze the weights of the decoder section during the fine-tuning process.

7. Training by reward network: The successful application of Human-feedback Reinforcement Learning (HFRL) to the training process for FMs has been established [126], and we borrow ideas from [89, 127]. We consider cell types as the human label for different cells, akin to the labels of sentences in the NLP area. A reward network is employed to predict the cell types of the provided cells and minimize the loss in this process. This procedure can also be integrated into the pre-training process or prompt-based learning framework. This approach can also be regarded as joint fine-tuning for Batch Effect Correction and Cell-type Annotation.

The results from this analysis provide valuable insights into how to configure scGPT most effectively for this specific task.

## C For developers: How to justify a good foundation model for single-cell analysis?

By summarizing our experimental results, here we list essential factors and criteria for selecting benchmarking datasets for developers of single-cell foundation models.

Regarding important factors, we recommend that developers and researchers consider the following:

- Including basic tasks to demonstrate its functionality. Current single-cell foundation models work well for basic tasks, including cell-type annotation, gene function prediction, and perturbation prediction. These tasks can be utilized as a starting point for developing foundation models.
- Improving model performances under challenging tasks. We demonstrated that single-cell foundation models could have performed better in batch effect correction, imputation, and simulation, which should be considered as hard but important tasks for resolving with new foundation models. For simulation, it is important to investigate the generator used for simulating the gene expression profiles to preserve the correct gene-gene correlation.
- Demonstrating the biological discoveries from foundation models. Training a foundation model has a high cost, and thus the developers may consider novel tasks in single-cell analysis and compare the foundation model with simple baselines and well-defined metrics. If necessary, developers are encouraged to collaborate with biologists for experimental verification.
- Exploring the model capacity for both zero-shot learning and fine-tuning. The ultimate task of a single-cell base model is to be able to understand the underlying biological system on a cellular basis, so it is important to measure the zero-shot learning capacity after model training. We understand that there still exits performance gap between zero-shot learning capacity and fine-tuning capacity for the same task, but we encourage the developers to explore effective approaches to enclose the gap or recommend better designs for the given task.
- Incorporating multimodel data in the training stage to increase capacity. The information from transcriptomic data is limited, and thus, introducing data from other domains might enhance the ability of single-cell foundation models to understand biology.
- Aligning single-cell foundation models with principles from other areas. There are some abilities of Large Language Models (for example, scaling law in various tasks and content generation) that are not observed in single-cell foundation models, so the developers may seek explanations or methods to bridge the gap.
- Ensuring trust and robustness. We demonstrated that single-cell foundation models suffer from unstable problems in specific tasks, and foundation models always risk being attacked. Therefore, developing a safe and robust model should be taken into consideration.
- Enhancing accessibility by open-source and user-friendly designs. We encourage open-science research, and thus, developing fully open-source models is recommended. Moreover, considering the hardware limitation of biologists, developing a model with fewer resources for deployment is encouraged.

Regarding dataset selection, we recommend developers and researchers consider the following:

- Selecting suitable datasets for pre-training is important. Many tasks in single-cell analysis can be handled by methods developed based on cell embeddings or gene embeddings without further fine-tuning, and thus, learning a good representation of cells and genes in the pre-training stage is important. The developers should consider this point in the data collection stage and the training/validation/testing separation stage.
- Evaluating the model with commonly used datasets. Considering the faster development speed in the single-cell analysis area, many techniques are outdated and have limited contribution even after integration. Therefore, evaluating the performances of single-cell foundation models in commonly used datasets (like scRNA-seq from 10X) is encouraged. Moreover, methods with functions for handling spatial transcriptomics should also consider datasets in different resolutions.
- Considering the diversity of datasets. Multiple biological factors can affect the variation of single-cell datasets, including tissue type, disease state, cell type, perturbation, etc. Ensuring and demonstrating that the model can learn information from various data sources are important.
- Avoiding data leakage. It is essential to ensure no intersection between training and testing datasets, and an improper split might inflate model performance in the testing stage.

## D Metrics Information

Here we describe details about the metrics we used in the evaluation process for different tasks.

### D.1 Batch Effect Correction, Multi-omic Data Integration and Simulation

1. Normalized Mutual Information (NMI): NMI is a score to evaluate the performance of biological information preservation. We calculate this score based on computing the mutual information between the optimal Leiden clusters and the known cell type labels and then take the normalization. Therefore, NMI ∈ (0, 1) and higher NMI means better performance.

2. Adjusted Rand Index (ARI): ARI is a score to evaluate the performance of biological information preservation. ARI can measure the agreement between optimal Louvain clusters and cell type labels. ARI ∈ (0, 1) and higher ARI means better performance.

3. Average Silhouette Width (ASW): Here we have cell type ASW (*ASW_cell_*) and batch ASW (*ASW_batch_*). For one cell point, ASW calculates the ratio between the inner cluster distance and the intra cluster distance for this cell. Therefore, higher *ASW_cell_* means better biological information preservation and lower *ASW_batch_* means better batch effect correction. To make them consistent, for *ASW_cell_*, we take the normalization, that is:

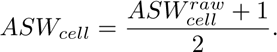

Similarly, for *ASW_batch_*, we take the inverse value of the normalization, that is:

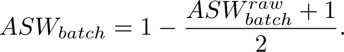

Both of metrics are in (0,1), and a higher score means better model performance. For multi-omic data integration, all the metrics are the same.

4. Graph Connectivity (GC): GC means the connectivity of cells in different cell types. If the batch effect is substantially removed, the connectivity of cells of the same cell type from different batches will have a higher connectivity score based on the k-NN neighbor graph. Therefore, we can calculate the GC score for each cell type and take the average. GC score is in (0,1) and higher means better batch effect correction performance.

5. Principal Component Regression (PCR): PCR is a metric to evaluate the performance of batch effect correction. We calculate the *R*^2^ for a linear regression of the covariate of interest onto each principal component. The variance contribution of the batch effect for all the PCs is based on the sum of the product between the variance of each PC and the *R*^2^ of each PC across all the PCs. Therefore, the score can be represented as:

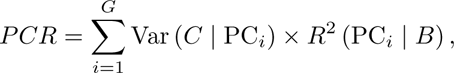

where *G* denotes the number of PCs and *B* denotes the batch information. PCR is in (0,1) and a higher score means better performance.

6. kBET: the kBET algorithm is used to determine if the label composition of the k-nearest-neighbors of a cell is similar to the expected label composition. For the batch label mixture of cells in the same cell types, the proportion of cells from different batches for the neighbors of one cell should match the global level distribution. The kBET score is in (0,1) and higher means better batch effect correction performance.

We computed the *S_bio_* based on the average value of NMI, ARI, and *ASW_cell_*. The clustering scores are computed based on searching the leiden resolution to achieve the best performances based on scIB. We computed the *S_batch_* based on the average value of *ASW_batch_*, GC, PCR, and kBET. *S_final_* = 0.6*S_bio_* + 0.4*S_batch_*. We also investigated the choices of weights for computing the final score. scIB chose to assign a higher weight for *S_bio_* based on the prior information that biological variation plays a more important role in the analysis. To compare the difference between the default setting of weights and other settings of weights, we designed two methods for computing the final rank of different methods across different datasets.

The first method we tried is to adjust the weights of *S_bio_* from 0.5 to 0.9, and then compute the metrics for each method under each dataset based on different weights. For each weight, we could compute the final score and ranked these methods by different datasets. Thus, we finally had five different rank tables. Now we averaged the ranks of different methods across datasets to get the final average rank of each method, and such value was used for our final evaluation. The result is shown in Extended Data Figure 36 (a).

The second method is the same as our original method. The result is shown in Extended Data Figure 36 (b).

Therefore, we only need to check the difference between the first method and the second method to see whether we need to optimize the weights for *S_bio_* and *S_batch_*or we can treat them based on our prior. We compared the average rank results from Extended Data Figure 36, and we found that the top three methods are exactly the same under different methods. Moreover, only the average rank of two different methods was affected by adjusting methods for computing the ranks, and their actual average rank is also very close. Furthermore, we computed the Spearman correlation coefficient between these two rank results, and we found that the correlation is 0.98 with a p-value smaller than 1e-55. Therefore, the results of these two methods are in strong correlation. It means that we do not need to adjust weights for different scores and the original choice of scIB and our manuscript is robust enough to evaluate the performance of different methods in the batch effect correction task.

For specific datasets with trajectory information, we also include the trajectory score to evaluate the conservation of trajectory information after batch effect correction. According to [61], to compute this score, we first calculate the diffusion pseudotime with a known initial cluster and trajectory relationship, and then we compute the Spearman correlation coefficient *s_sp_* between the pseudotime values before and after integration. Therefore, the trajectory score is defined as:

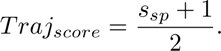

Higher *Traj_score_*means better trajectory conservation performance.

These metrics are also used for evaluating Multi-omic Data Integration. For simulation analysis, we only evaluated the biological information preservation for the simulation output with batch effect setting.

We also considered qualitative assessment. By visualizing the cell embeddings of these tasks using UMAPs, we could evaluate the performance of different methods for removing the batch effect and preserving the biological variation. A dataset without large batch effect should have a pattern that cells from the same cell type but different batches in the same or similar clusters.

### D.2 Cell-type Annotation and Gene Function Prediction

1. Accuracy: We calculate the accuracy based on the ratio between the total number of cells (or genes) being classified correctly and the total number of cells (or genes).

2. Precision, Recall, and F1: For classification results, we calculate true positives (*tp*), false positives (*fp*), true negative (*tn*), and false negatives (*fn*). Precision, Recall, and F1 score are calculated as follows:

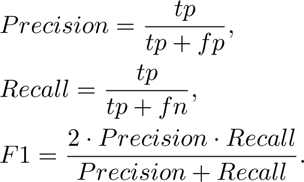

Note that precision, recall, and F1 are calculated based on the weighted average, where the weight of each cell type is determined by the cell number.

For the Gene Function Prediction task, all of the metrics are the same but for genes rather than cells.

### D.3 Perturbation Prediction

We use the Mean Pearson Correlation (MPC) and Mean Squared Errors (MSE) in the validation dataset to evaluate the performance of scGPT across different datasets.

Considering we have *n* genes in total. The Pearson correlation for the predicted gene expression *g*^′^ of gene *i* across all the cells and the ground truth gene expression *g_i_* of gene *i* across all the cells is:

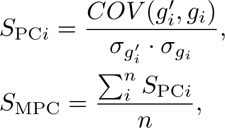

where *COV* (, ) is used to compute the covariance of two vectors, and *σ* is the standard deviation of the given vector. Higher *S*_MPC_ means better performance.

Similarly, we have MSE defined as:

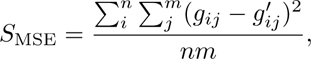

where *m* represents number of cells. Lower *S*_MSE_ means better performance.

### D.4 Gene Network Analysis

Here we extract the genes related to Human Immunology pathway. Moreover, we extract genes that have significant correlations with CD3-genes from our gene embedding dataset. We then measure the Jaccard similarity for these two gene sets *G_path_* and *G_net_* as follows,

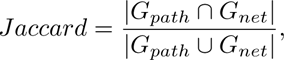

where *G_path_* represents the genes contained in the prior known pathway, and *G_net_* represents the genes contained in the GCN generated by single-cell FMs. The Jaccard similarity measures the similarity between two gene sets to evaluate the quality of GCN inference proposed by single-cell FMs. Higher Jaccard similarity means better performance.

Moreover, we performed the gene enrichment analysis for genes from *G_net_*. We extracted the pathways enriched in these genes and calculated the total number of significant pathways, after which we divided this number by the number of all pathways to get the final ratio. Higher ratio means better performance. The p-value threshold was adjusted based on Bonferroni correction.

We also considered qualitative assessment. By visualizing the gene embeddings of these tasks using UMAPs, we could evaluate the performance of different methods for generating meaningful embeddings by checking the co-embedded conditions for marker genes with similar expression patterns by cell type. Such marker genes from the same cell type should be located in the same cluster, while the difference of marker genes from different cell types should be preserved, in a space from good gene emebddings.

### D.5 Imputation

1. Average bio: The average bio score is calculated based on cell-type NMI and ARI scores using the latent space.

2. Correlation: The correlation score is calculated based on the average correlation between the raw data and imputed data for known common genes. The justification of this metric is based on the assumption that the target of spatial imputation is to predict the expression of missing or unmeasured genes while keeping the distribution of known gene expression levels. We use Pearson correlation here.

3. Significance proportion: The significance proportion corresponds to the proportion of genes with their p-value of correlation smaller than a specific threshold, which is 0.005 in our evaluation. The proportion can be used as a metric to evaluate the similarity between two paired data.

## E Evaluation Methods

Here we describe the details of the implementation of the methods discussed in this manuscript.

Single-cell FMs include:

- tGPT: tGPT is a single-cell FM based on the GPT-2 [128] structure. It utilized large-scale scRNA-seq datasets for pre-training and set the pre-training task as predicting gene expression rankings. The downstream applications of tGPT follow the zero-shot learning framework and include clustering, batch effect correction, and bulk RNA-seq analysis. We evaluate the performance of tGPT for Batch Effect Correction, Cell-type Annotation, and Perturbation Prediction.
- scBERT: scBERT is a pre-training-based single-cell FM focusing on cell-type prediction. It is based on Performer [129] with gene embeddings initialized by Gene2vec [59]. It has six self-attention blocks. The default fine-tuning process of scBERT on downstream datasets is freezing the penultimate layer. scBERT is considered for Cell-type Annotation.
- Geneformer: Geneformer is a single-cell FM using transfer learning to predict cell types and gene functions. The Geneformer tokenization step is done based on ranking gene expression values in single cells following scaling across the whole training dataset. Cells are represented as token strings with genes rankings as tokens. We evaluate the performance of Geneformer for Batch Effect Correction, Cell-type Annotation, Gene Function Prediction, Perturbation Prediction, and Gene Network Analysis. Geneformer is used for the Cell-type Annotation task and the Gene Function Prediction task. Both tasks were performed by fine-tuning on the basis of the published pre-trained Geneformer network. The default hyper-parameters were used for Geneformer fine-tuning. Prior to tokenization, gene names in all datasets were converted to ENSEMBL IDs using python packages *mygene* [130] and *pyensembl* [131], and most of unmatched genes are non-coding RNAs. For consistency with other benchmarked FMs, Geneformer’s workflow in Gene Function Prediction and non-cross Cell-type Annotation was altered to split all labeled genes into the training and the testing datasets using the same manners mentioned in our Methods section, instead of using the cross-validation evaluation method provided by the authors.
- CellLM: CellLM is a single-cell FM using three different pre-training strategies. The pre-training loss function includes: 1. masked gene expression level reconstruction; 2. cell condition discrimination; and 3. self-supervised contrastive learning. Moreover, it incorporated protein-protein interaction networks as prior information during the pre-training process. The downstream tasks of CellLM are all related to Cell-type Annotation. Here we modified CellLM so that it can perform cross-dataset cell-type annotation, by separating the training and the testing datasets.
- scFoundation: scFoundation employs a pre-training methodology similar to BERT and introduces Bayesian down-sampling as a data pre-processing step. The input data of scFoundation also contain target total counts and input total counts as extra information. The downstream tasks of scFoundation include clustering (a function of cell embeddings across all models), drug response prediction (belong to Cell-type Annotation) and Perturbation Prediction. We evaluate the performance of scFoundation for Batch Effect Correction, Cell-type Annotation, Perturbation Prediction, and Gene Network Analysis.
- SCimilarity: SCimilarity declares that it serves as a foundation model for new data querying or searching based on the cell embeddings generated from known large-scale scRNA-seq datasets. It pre-trains a MLP rather than transformer-based models. The downstream tasks of SCimilarity include Batch Effect Correction and Cell-type Annotation. We evaluate the performance of SCimilarity for Batch Effect Correction, Cell-type Annotation, and Perturbation Prediction.
- CellPLM: CellPLM utilizes cells as tokens and also pre-trains a transformer-based model based on both scRNA-seq datasets and spatial transcriptomic datasets. CellPLM also constructs the latent space based on Gaussian Mixture priors and their decoder part accepts the latent variables sampled from this latent distribution. We evaluate the performance of CellPLM for Batch Effect Correction, Cell-type Annotation, and Imputation. The codes for Perturbation Prediction based on CellPLM were not evaluated because they did not release codes for performing this task.
- UCE: UCE utilizes genes as tokens and sorts the gene expressions by the genomic location of genes. UCE pre-trains a transformer-based model augmented by gene tokens from a large protein language model (PLM) based on mega-scale atlas scRNA-seq datasets. UCE is capable of Batch Effect Correction, Cell-type Annotation, and Gene Expression Prediction. We evaluate the performance of UCE for Batch Effect Correction, Cell-type Annotation, and Perturbation Prediction because they did not release codes for performing other tasks.
- GeneCompass: GeneCompass pre-trains a transformer-based model with external knowledge embeddings including Gene Regulatory Network (GRN), Promoter, Gene family, and Gene co-expression. It has an expression decoder and gene ID encoder to perform self-supervised learning. GeneCompass can perform tasks including Celltype Annotation, Perturbation Prediction, Dose Response Prediction, and GRN Inference. We did not evaluate the performance of GeneCompass because it is totally closed-source with no pre-training weights and instructions.

Task-specific methods are SOTA models in their areas based on their description. They include:

- ResPAN: ResPAN is a batch effect correction tool based on Generative Adversarial Network (GAN) [132, 133]. The high-level idea of ResPAN is based on the distribution alignment or domain adaption across data from different batches. Such requirement can be treated as optimal transport, which can be accomplished by training a GAN. ResPAN is used for evaluating the batch effect correction task.
- Harmony: Harmony is a batch effect correction tool starting from PCs based on self-supervised learning and vector correction. Harmony sets up the initial clustering labels for uncorrected data. For cells with the same cluster labels, Harmony performs correction for cell embeddings of each cluster, and then repeats the clustering process until convergence. Harmony is used for evaluating the Batch Effect Correction task.
- scVI: scVI is a batch effect correction tool based on variational inference and variational auto-encoder [134]. scVI encodes the gene expression data with batch information using a neural network and sets the output of the network as parameters for a distribution. Based on such distribution of the latent space, scVI can correct the batch effect in the latent space as well as the original space, as long as we consider the output of the decoder model.
- scJoint: scJoint is a multi-omic data integration tool based on an auto-encoder structure. It contains three steps. In the first step, scJoint performs semi-supervised transfer learning. In the second step, scJoint performs label transfer based on the kNN classification for the joint cell embeddings. In the last step, scJoint starts joint training for the labeled data to optimize the cell embeddings. scJoint is used for evaluating the Multi-omic Data Integration task.
- scGLUE: scGLUE (GLUE) is a multi-omic data integration tool based on VAE and Graph Neural Network (GNN). It contains two parts. For the feature-encoding part, scGLUE utilizes a Knowledge-based guidance graph with GNN to generate feature embeddings. For the cell-encoder part, scGLUE utilizes VAE to generate cell embeddings. The decoder process is constructed by performing inner-product based on the cell embeddings and feature embeddings. scGLUE is used for evaluating the Multi-omic Data Integration task.
- *Vanilla* NNs: This neural network contains three MLP layers with batch normalization and uses Mish [135] as the activation function. *Vanilla* NNs are used for evaluating the Cell-type Annotation task and the Gene Function Prediction task. We trained *Vanilla* NNs based on different input datasets. The learning rate is set as 1e-4, the optimizer is Adam, and the epoch is set as 10. We tuned the best model by splitting training datasets and testing datasets for different tasks.
- TOSICA: TOSICA is a deep learning-based method for one-stop cell type annotation. TOSICA is designed with the self-attention multi-heads transformer without pre-training. It also provides interpretation for researchers about the attention embeddings and uses attention embeddings to perform biological analysis. TOSICA is used for evaluating the Cell-type Annotation task.
- SVM_rej_: SVM_rej_ is a machine learning based classifier for cell-type annotation. This method is designed based on support vector machine with calibration settings. This method was evaluated in [58] and was a top-tier method. SVM_rej_ is used for evaluating the Cell-type Annotation task.
- Gene2vec: Gene2vec is a tool to generate gene embeddings based on Word2vec [136]. It trains a Word2vec model to predict the gene-gene interaction using known datasets, and generates gene embeddings of these genes after training. Gene2vec is used for evaluating the Gene Function Prediction task.
- GEARS: GEARS is a tool for single and multi-gene perturbation prediction based on single-cell RNA sequencing datasets. It combines gene-gene interaction network as prior information and uses a cross-gene neural network with a graph neural network to predict gene expression after perturbation.
- Tangram: Tangram is a toolbox for spatial transcriptomic data analysis based on neural networks. The key idea behind Tangram is using neural networks to find a good mapping function from single-cell data space to spatial data space. After the mapping process, by integrating the information from the single-cell level and spatial level, it can perform several downstream tasks, including data imputation, cell-type deconvolution, and others. Tangram is used for evaluating the Imputation task.
- scDesign3: scDesign3 is a model based on Copula distribution [137] to generate different single-cell datasets. Such datasets can be multimodal. Moreover, based on the input parameters and requirements of scDesign3, it can also generate datasets with specific conditions, including batch effect, cell conditions, and the stages of cell differentiation. The data generation of scDesign3 is based on real datasets. scDesign3 is used for evaluating the Simulation task.
- Splatter: Splatter is a model based on joint probabilistic inference. It models scRNA-seq data by estimating certain parameters from known scRNA-seq data. Then Splatter combines the estimated parameters with additional parameters to simulate datasets under different conditions. All the distributions used in Splatter are the known distributions. Splatter is used for evaluating the Simulation task.
- SATURN: SATURN is a model designed for learning cross-species cell embeddings and integrating datasets from different species. It has two stages, including a pretraining stage as well as a fine-tuning stage based on supervised contrastive learning by using the matched cell-type labels. To make a fair comparison for the cross-species cell-type annotation analysis, we utilize the cell embeddings of SATURN from the pre-training stage to annotate the cell types for datasets from different species.
- Novae: Novae is a model pre-trained with large-scale spatial transcriptomics data. It is based on the optimal transport and contrastive learning to learn the spot representations in a joint space. We utilize Novae in evaluating the batch effect correction performance for spatial transcriptomics data.

## F Model Architecture

The model architecture of scGPT, scBERT, CellLM, Geneformer, tGPT, SCimilarity, UCE, CellPLM and scFoundation are provided in the Supplementary file 5.

## G Data availability

No new sequencing data were generated for this current study. Supplementary file 6 provides the sources and download links for the datasets used in each task. These datasets come from the following papers:

Batch Effect Correction: [48, 61, 108, 138–140].

Cell-type Annotation: [23, 48, 61, 108, 141, 142].

Gene Function Prediction: [45].

Multi-omic Data Integration: [143–146].

Perturbation Prediction: [60].

Imputation: [73].

Gene Network Analysis: [61, 108].

Simulation Analysis: [108].

Model Scaling: [108, 141, 147].

## H Running statistics

We summarized the running time, peak CPU memory usage and peak GPU memory usage, in Supplementary file 7.

