## Supplementary files. for "Evaluating the Utilities of Foundation Models in Single-cell Data Analysis": Supplementary figures.pdf

### I Supplementary Figures

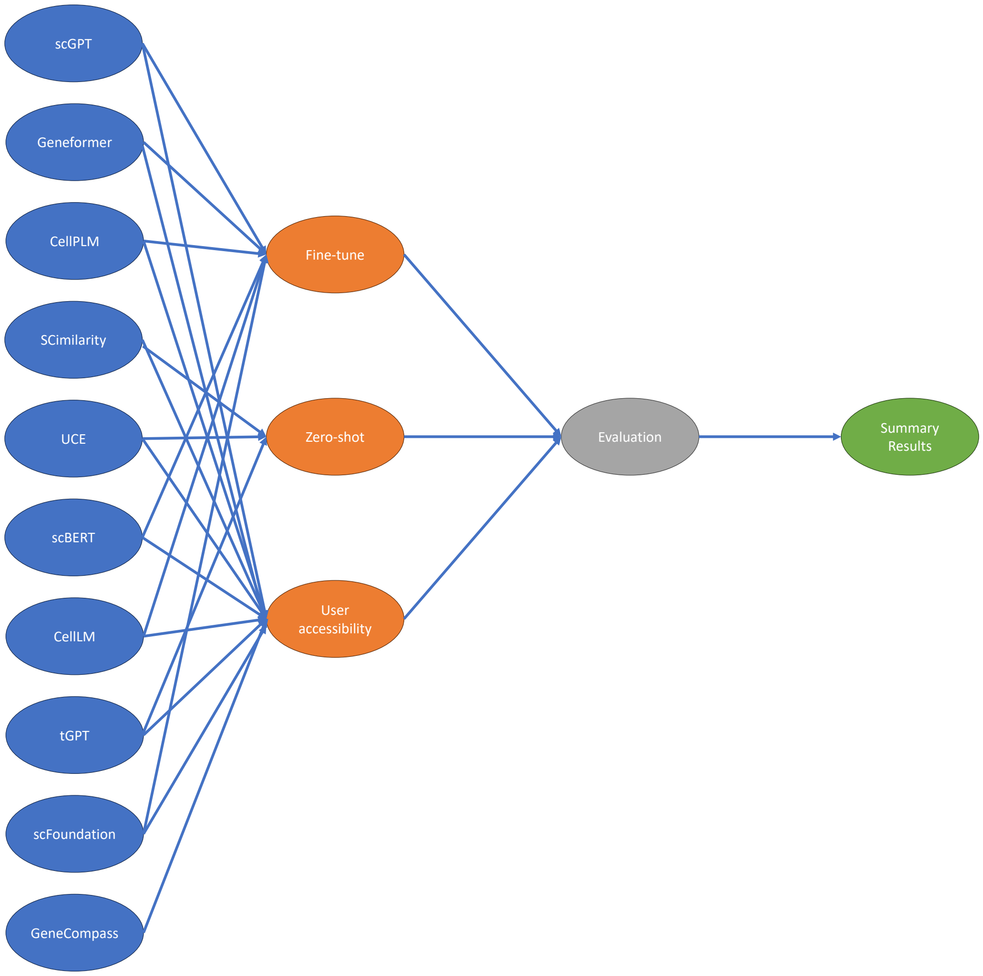


**Extended Data Fig. 1**: Workflow of our evaluation. Here we considered ten single-cell FMs and generated their outputs based on their default usage settings. We also consider the evaluation of user accessibility for both open-source and closed-source tools. We then evaluated the outputs with task-specific metrics. Finally, we summarized our discoveries in this manuscript.

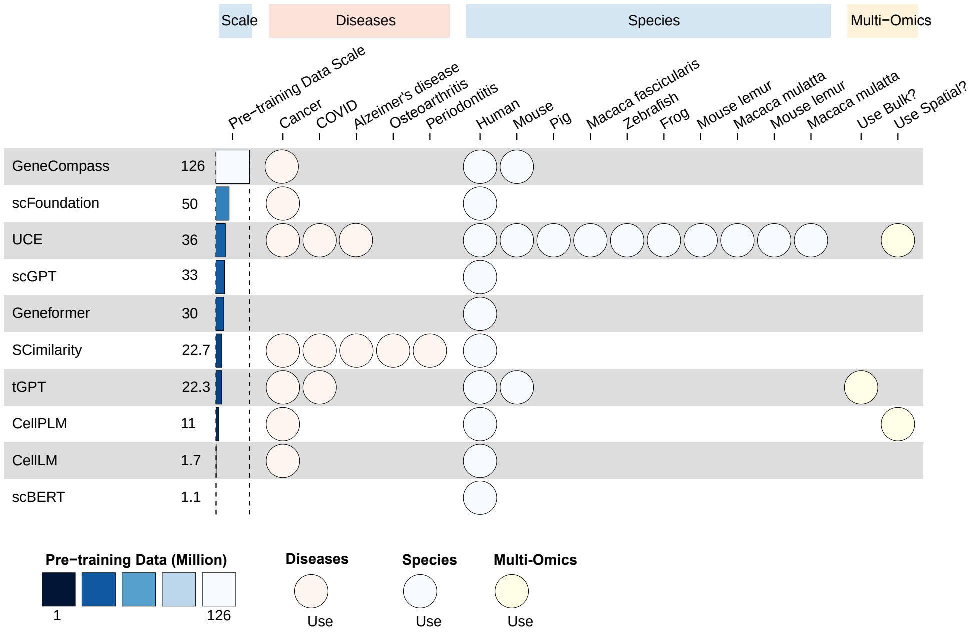


**Extended Data Fig. 2**: Statistics of pre-training datasets for different single-cell FMs. Here we consider summarizing the scale of pre-training datasets, the overlap for major diseases in the pre-training datasets, the overlap for major species in the pre-training datasets, and the overlap of multi-omics in the pre-training datasets, across all the methods. The scale of pre-training dataset is a continuous variable, whether other statistics are recorded based on a binary variable.

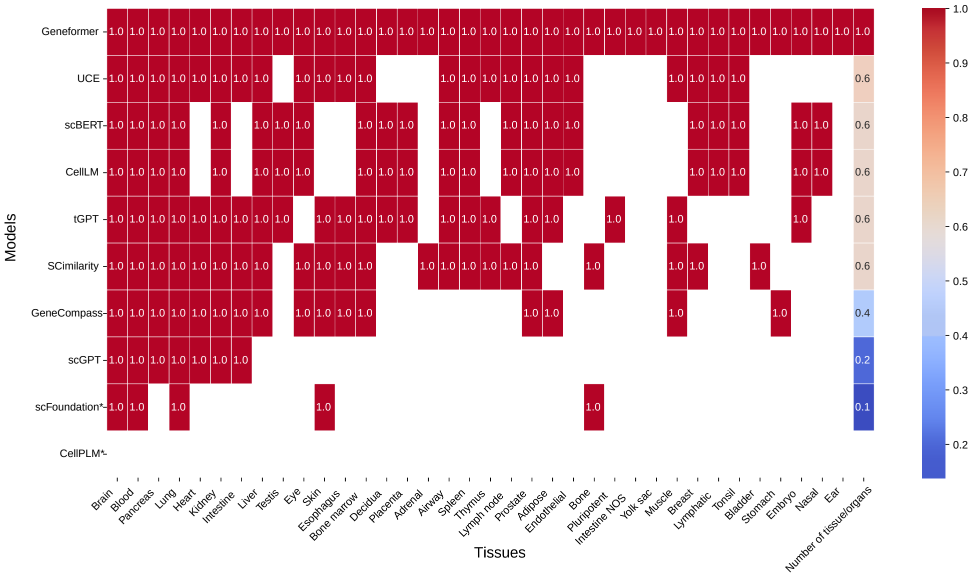


**Extended Data Fig. 3**: The overlap of major human tissues or organs across different single-cell FMs' pre-training datasets. Here we choose tissues or organs included by Geneformer as a baseline because it is published and it includes most of the major information for the cells from human. *: For scFoundation, we only record the major tissues or organs based on its manuscript since we did not find the data sources description from its manuscript and supplementary files. For CellPLM, we did not record the major tissues or organs since we did not find the data source description from its manuscript and supplementary files.

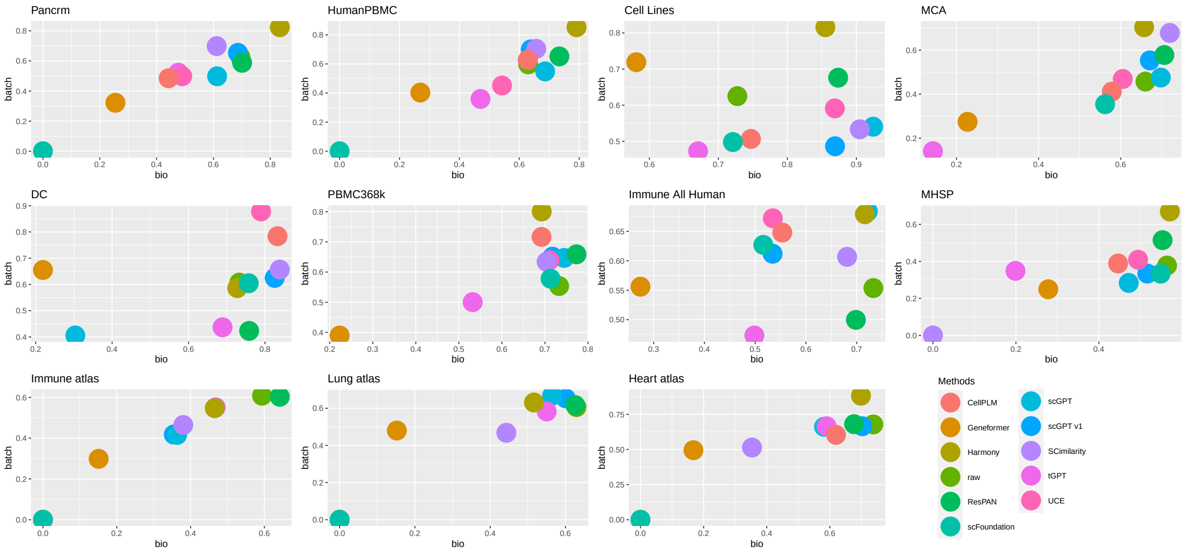


**Extended Data Fig. 4**: Visualization of batch effect removal score and biology variation conservation score in the evaluation of batch effect correction task for each dataset.

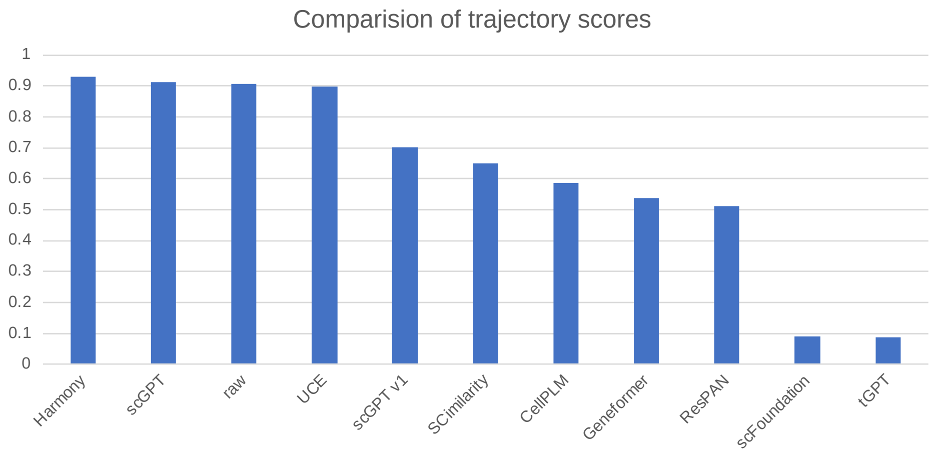


**Extended Data Fig. 5**: Comparisons for different methods on preserving the trajectory information based on the trajectory score.

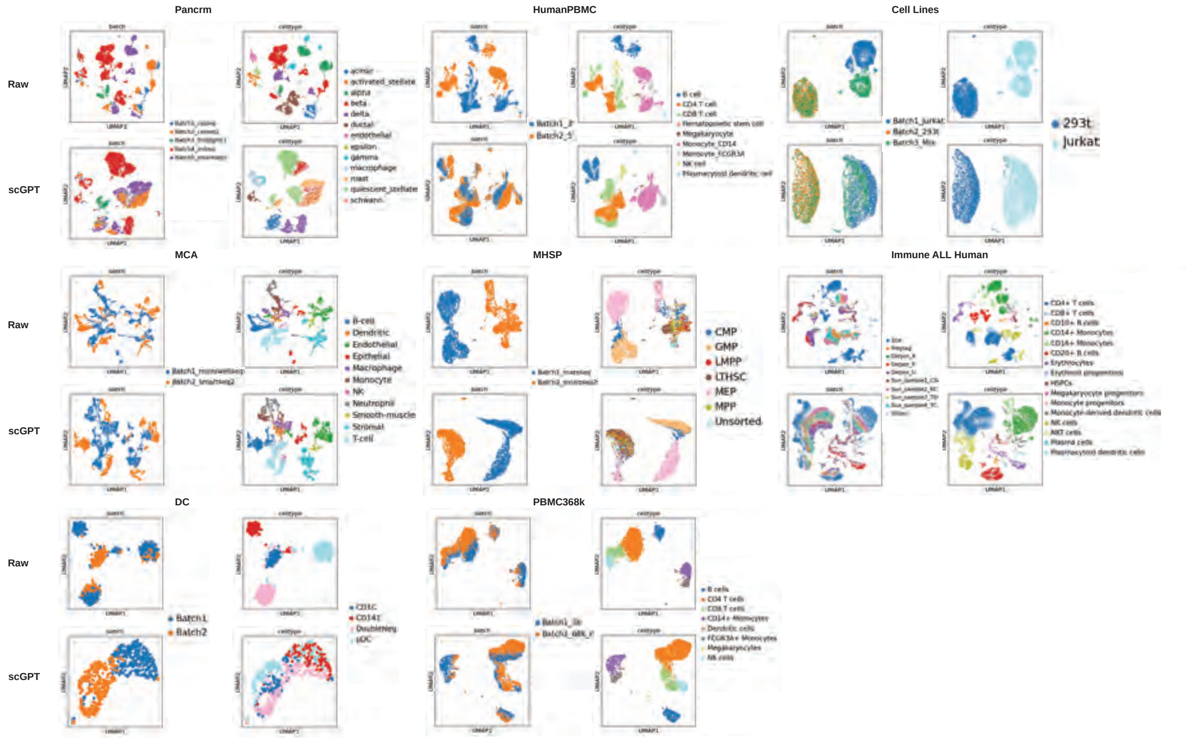


**Extended Data Fig. 6**: UMAPs for raw data and embeddings of scGPT after batch effect correction.

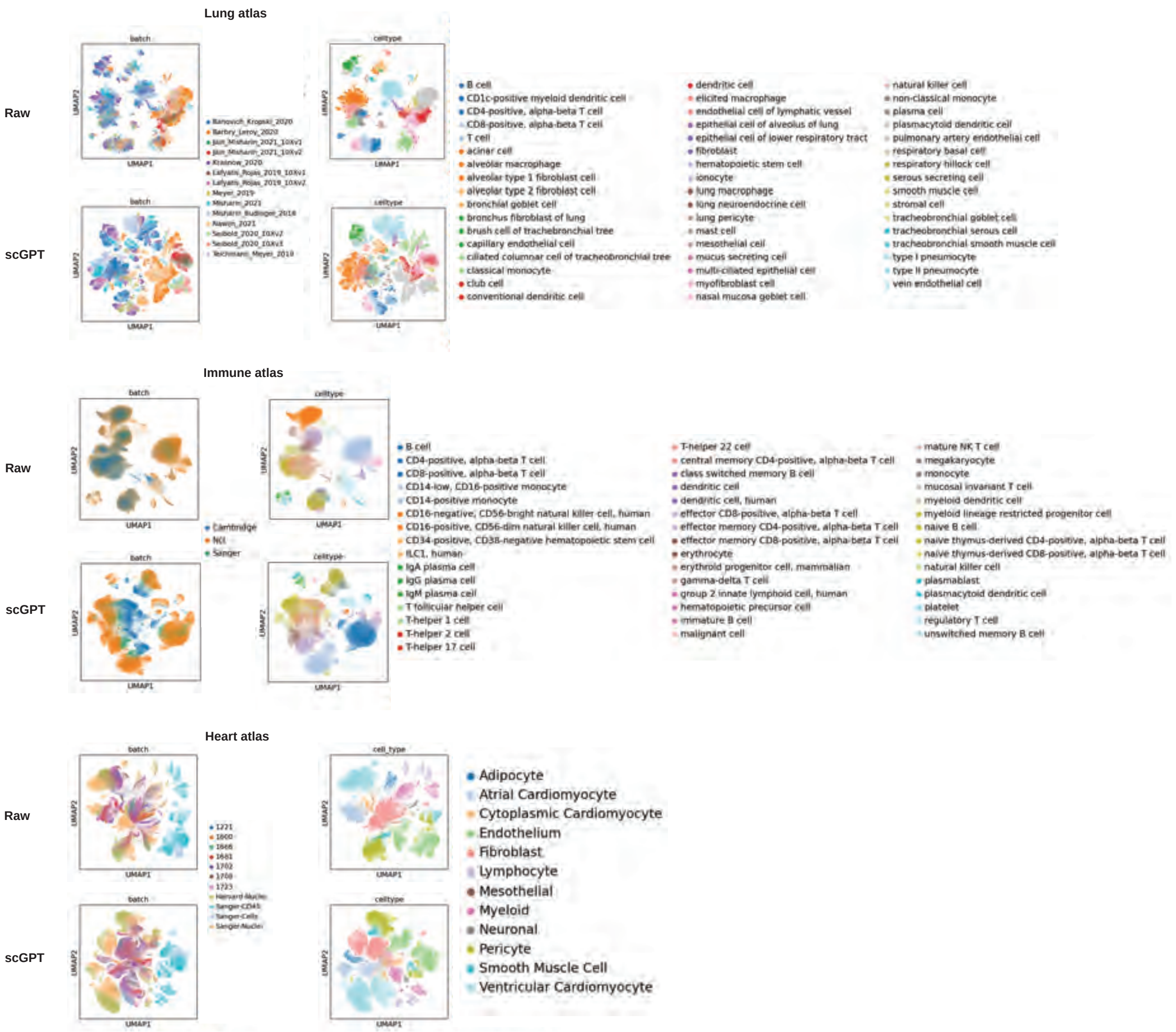


**Extended Data Fig. 7**: UMAPs for raw data and embeddings of scGPT after batch effect correction.

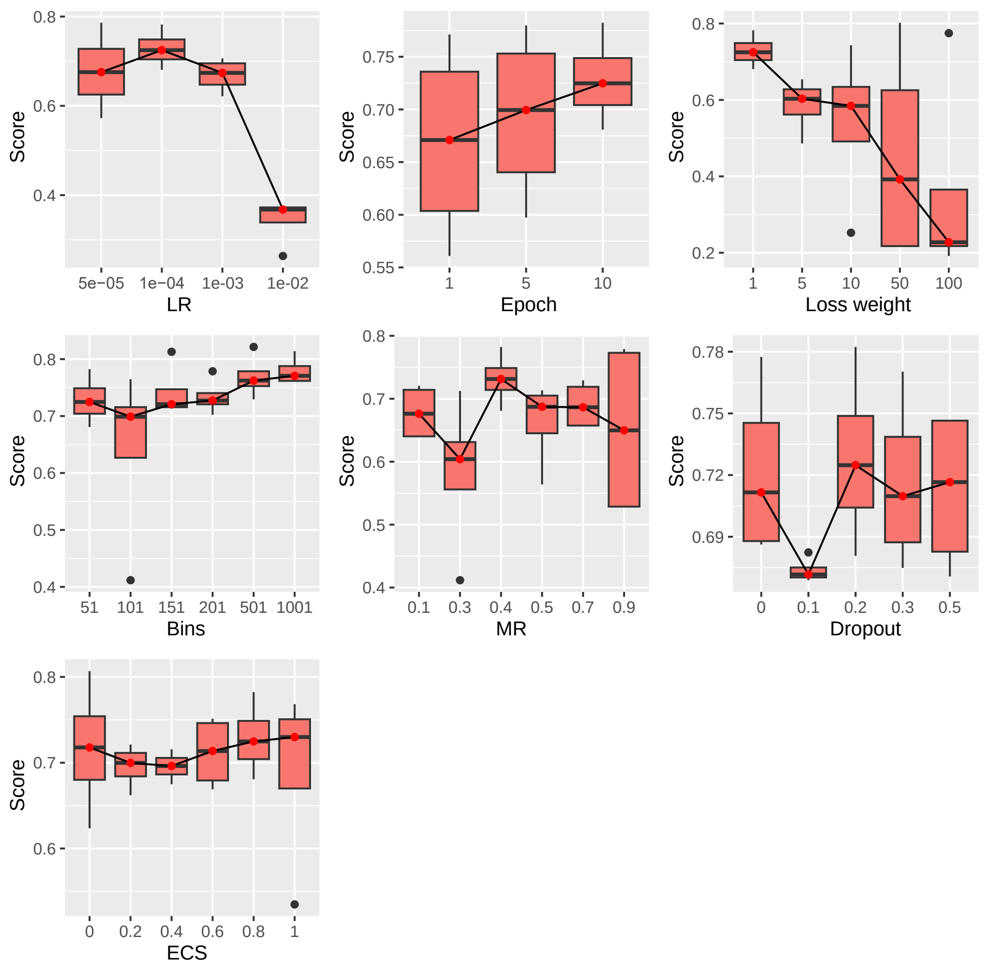


**Extended Data Fig. 8**: Benchmarking results for different hyper-parameters for batch effect correction.

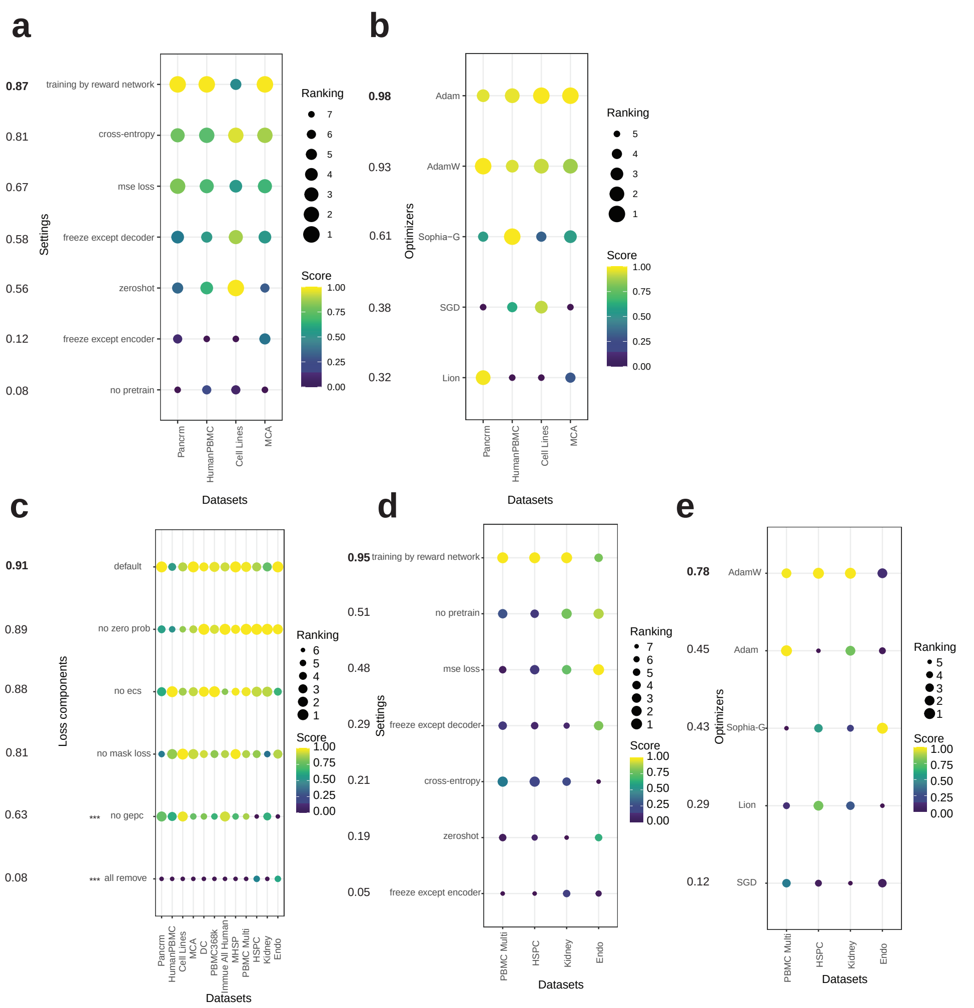


**Extended Data Fig. 9**: Overall evaluation of different components of scGPT. (a): The final score of different settings across different datasets for batch effect correction. (b): The final score of different optimizers across different datasets for batch effect correction. (c): The final score of including different loss function terms across different datasets. The number of stars represents the significance level ($***: p - value < 0.005, **: p - value < 0.05, *: p - value < 0.1$). The numbers on the left side of each sub-figure represent the average score across different datasets for one condition. We combine the datasets for batch effect correction and multi-omic data integration for testing. (d): The final score of different settings across different datasets for multi-omic data integration. (e): The final score of different optimizers across different datasets for multi-omic data integration.

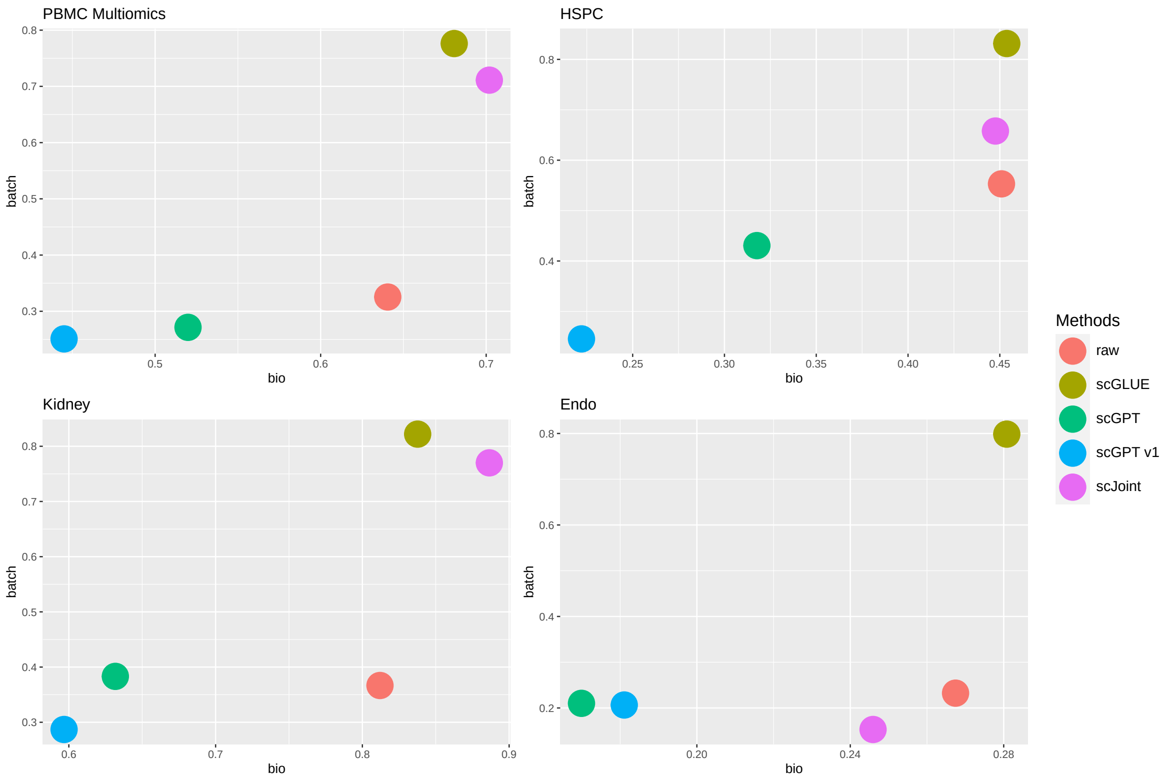


**Extended Data Fig. 10**: Visualization of batch effect removal score and biology variation conservation score in the evaluation of multi-omic data integration task for each dataset.

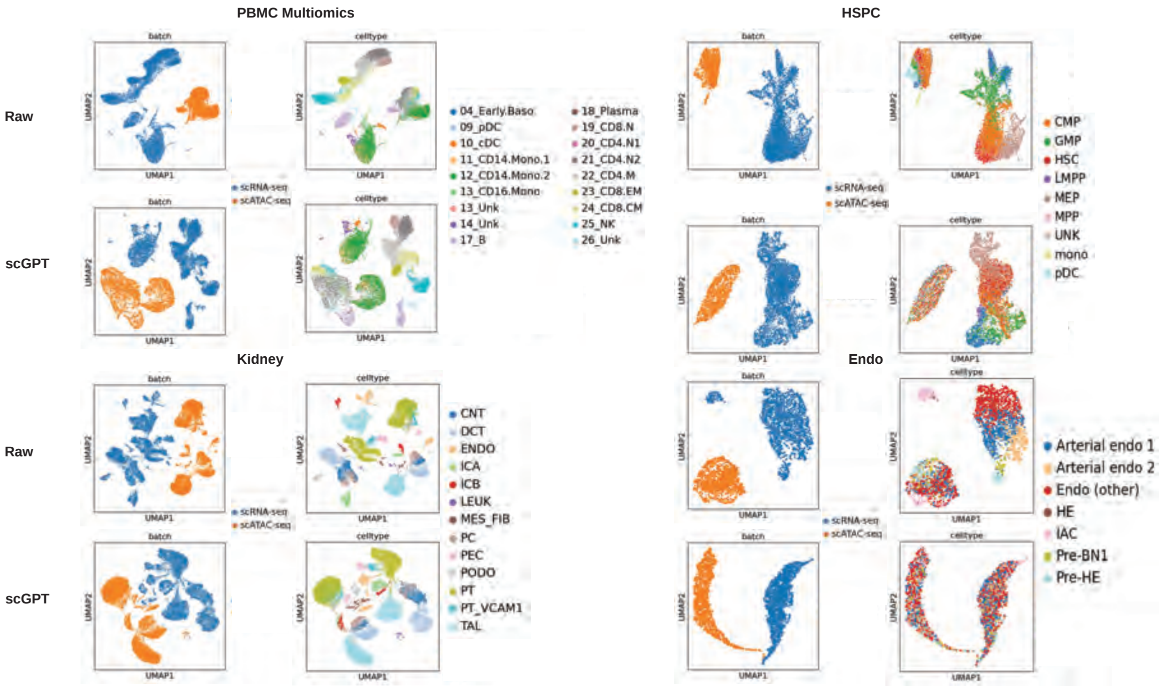


**Extended Data Fig. 11**: UMAPs for raw data and embeddings of scGPT after multi-omic data integration.

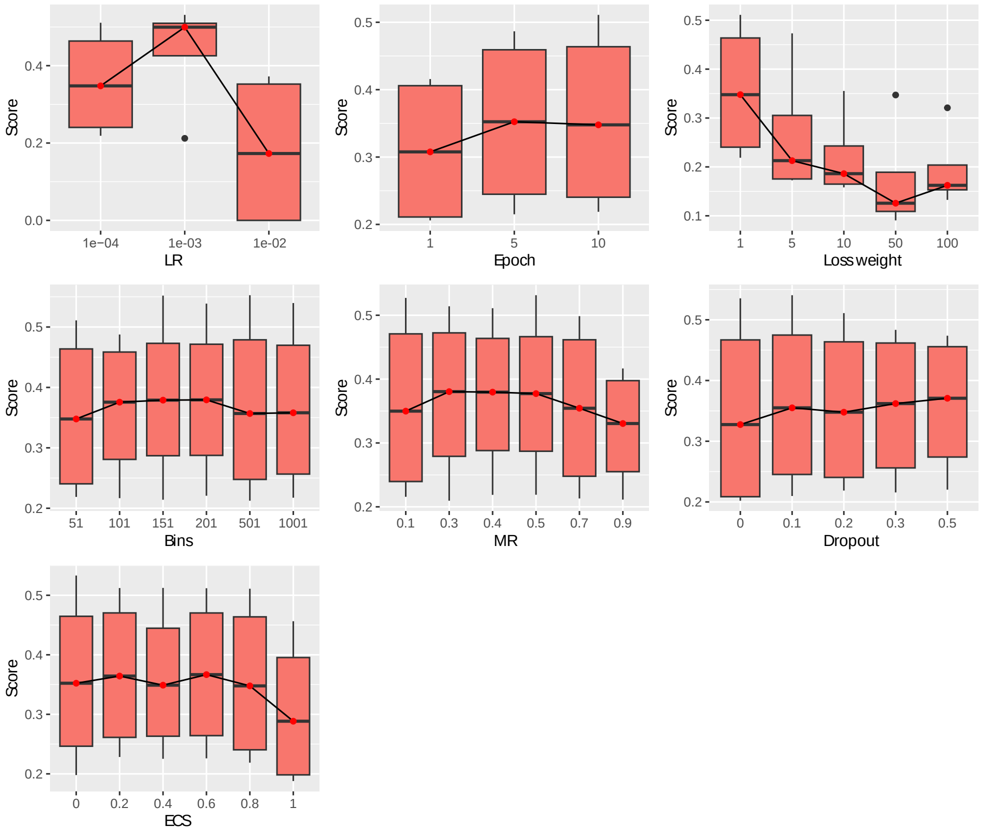


**Extended Data Fig. 12**: Tuning parameters for multi-omic data integration. Subfigures represent the score of scGPT under different hyper-parameters after training.

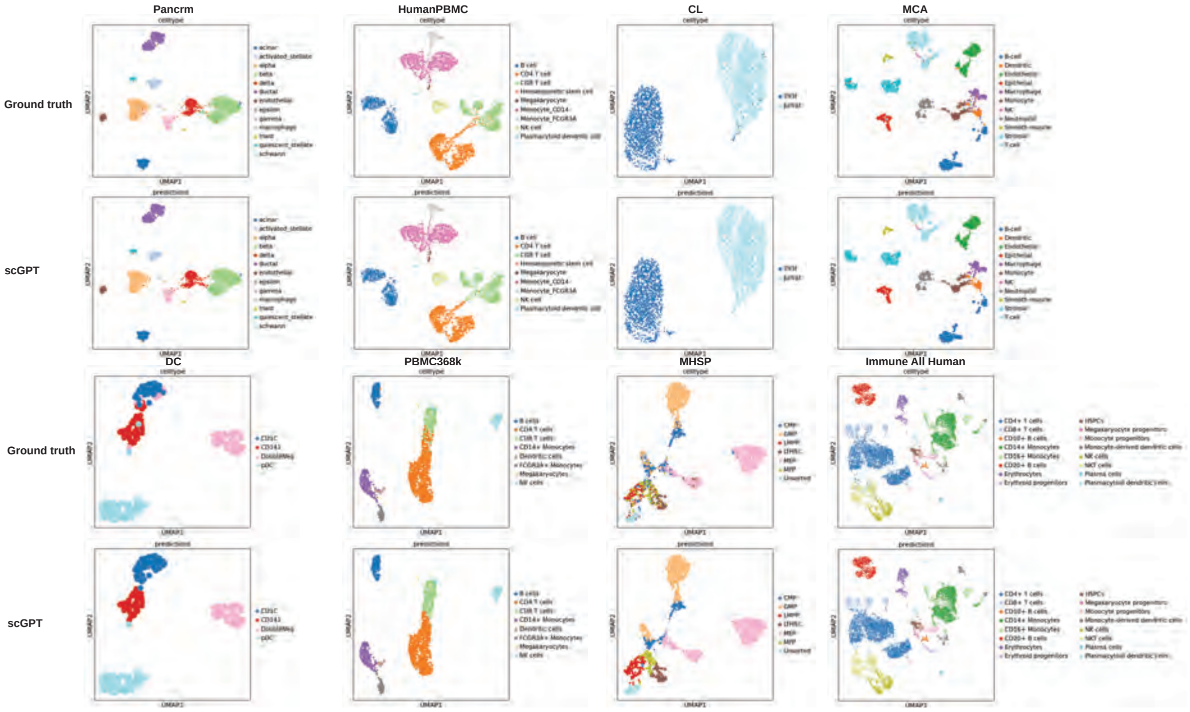


**Extended Data Fig. 13**: UMAPs for ground truth cell types and prediction results based on scGPT.

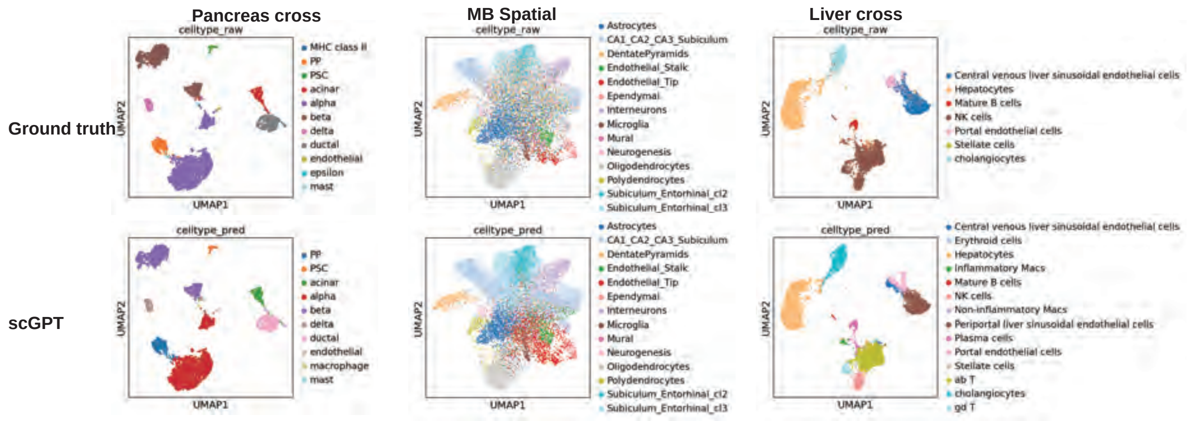


**Extended Data Fig. 14**: UMAPs for ground truth cell types and prediction results based on scGPT.

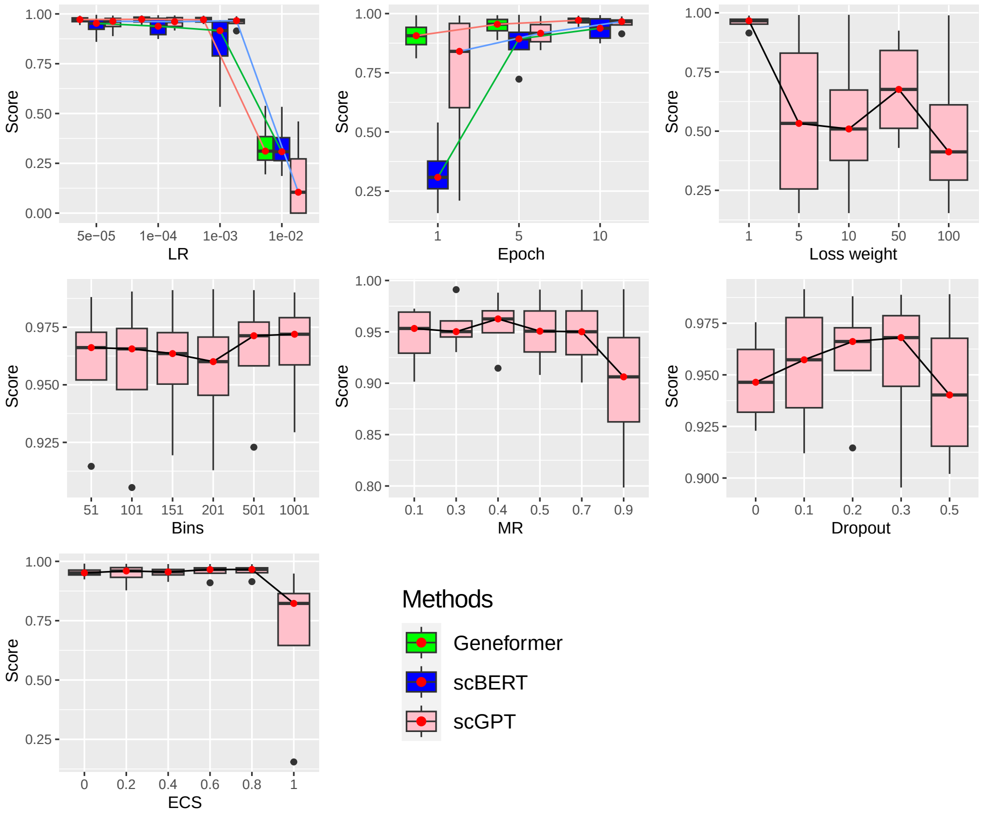


**Extended Data Fig. 15**: Tuning parameters for cell-type annotation. Sub-figures represent the score of scGPT under different hyper-parameters after training.

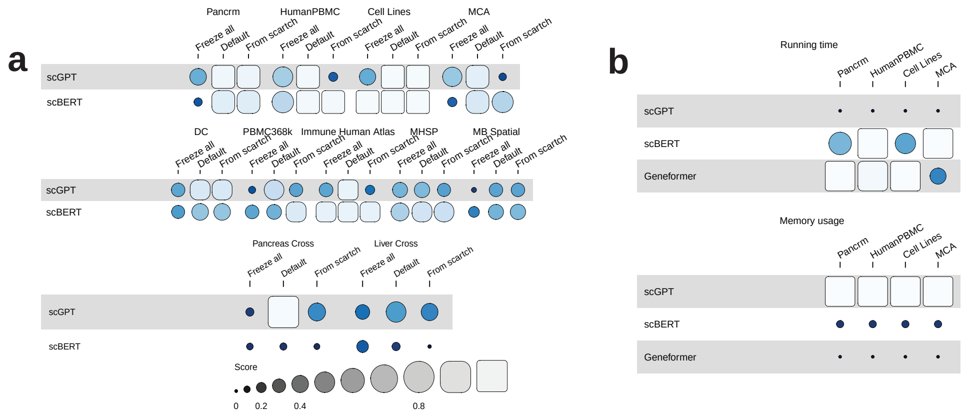


**Extended Data Fig. 16**: Results of different settings, running time, and memory usage for cell-type annotation task. (a): Accuracy of scGPT and scBERT for the Cell-type Annotation task across different datasets. (b): Scaled running time (up) and scaled memory usage (down) statistics for all three single-cell FMs.

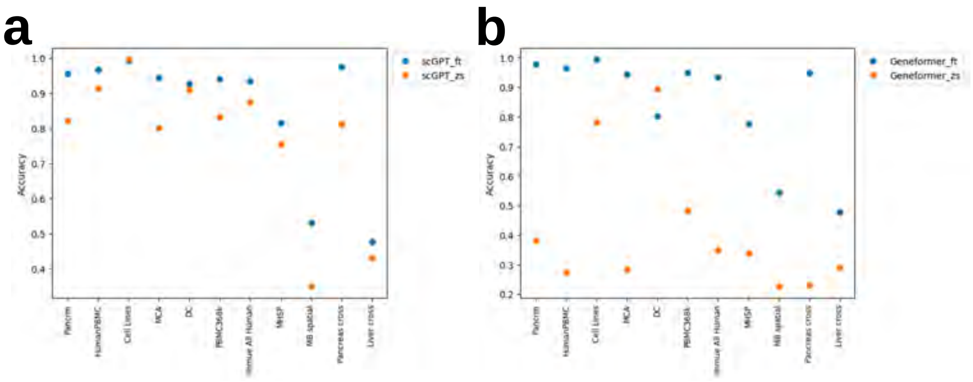


**Extended Data Fig. 17**: Results of cell-type annotation with the different modes of scGPT and Geneformer. (a) The performance of cell-type annotation of scGPT based on both fine-tuning mode and zero-shot learning mode. (b) The performance of cell-type annotation of Geneformer based on both fine-tuning mode and zero-shot learning mode.

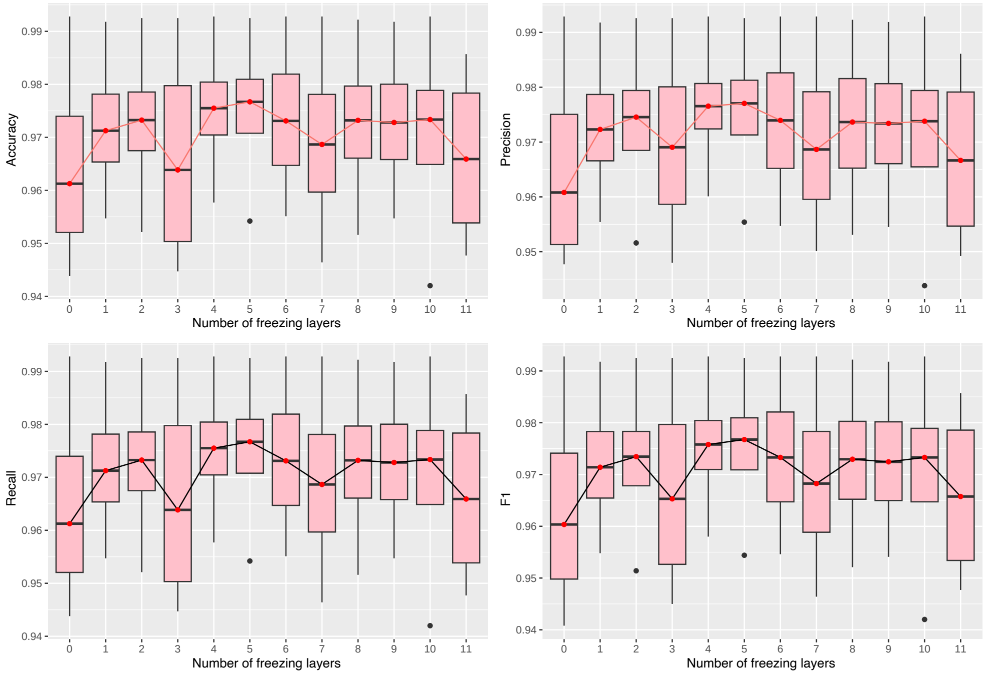


**Extended Data Fig. 18**: Results of cell-type annotation with the different number of freezing layers based on scGPT. The x-axis represents the number of freezing layers and the y-axis represents the value of specific metrics.

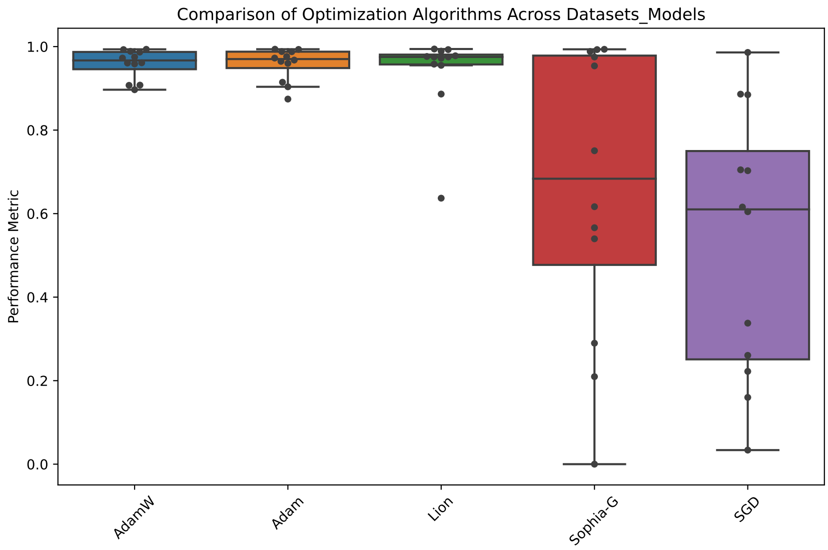


**Extended Data Fig. 19**: Benchmarking results for different optimizers for cell-type annotation. Here each box contains the scores of fine-tuned scFMs across different datasets.

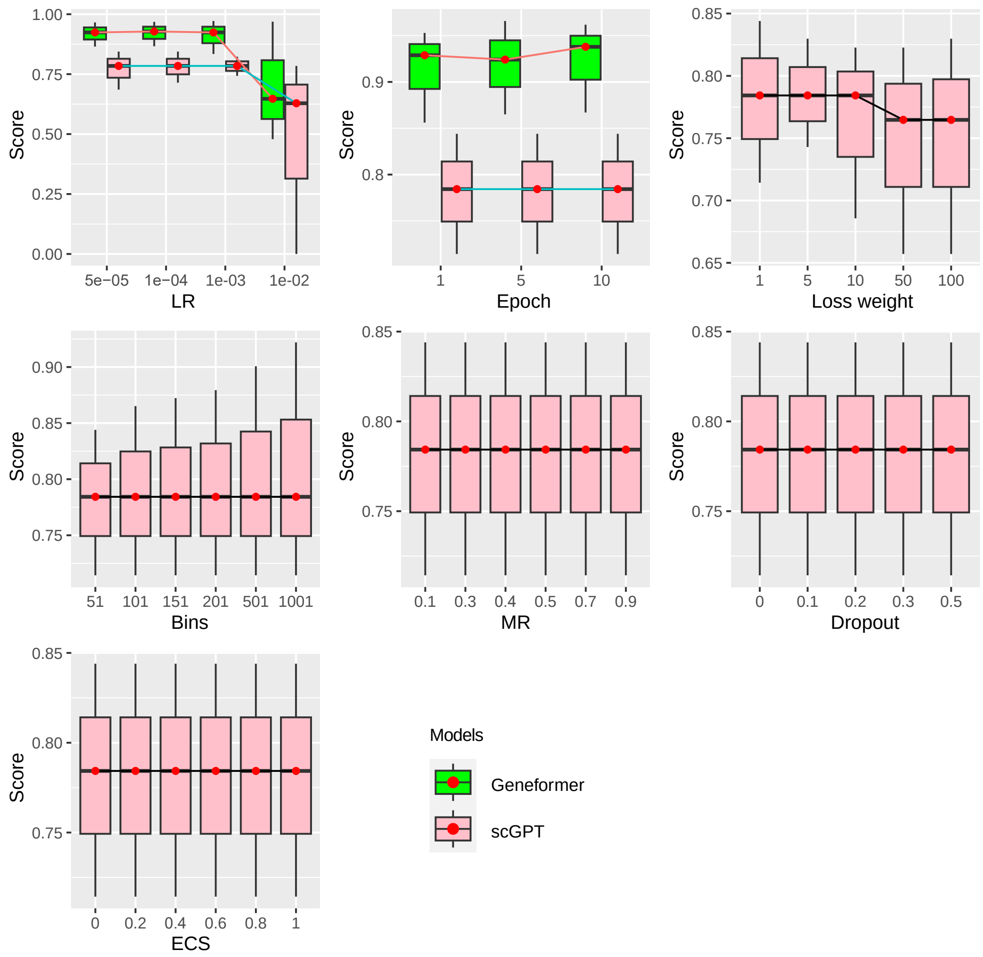


**Extended Data Fig. 20**: Tuning hyper-parameters for gene function prediction. Subfigures represent the score of scGPT under different hyper-parameters after training.

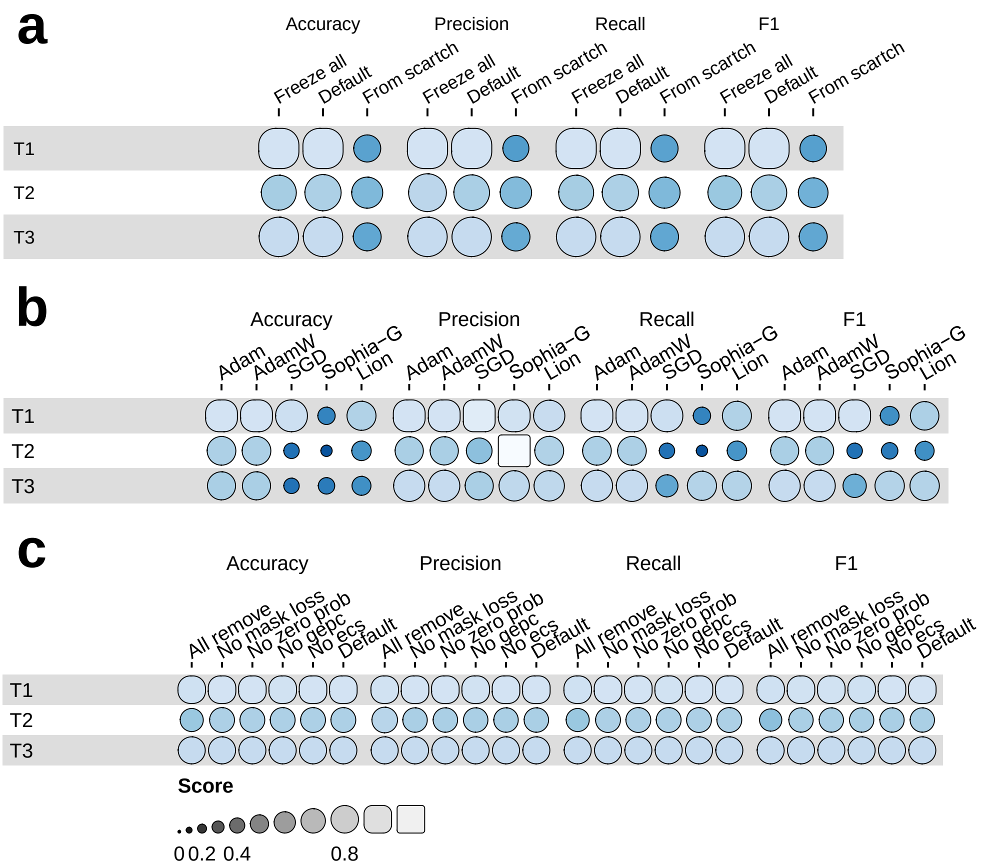


**Extended Data Fig. 21**: Benchmarking results of different initial settings, optimizers, and loss components for gene function prediction. T1-T3 represent different gene prediction cases. (a): Results of adjusting initial settings. (b): Results of adjusting optimizers. (c): Ablation tests based on different loss components.

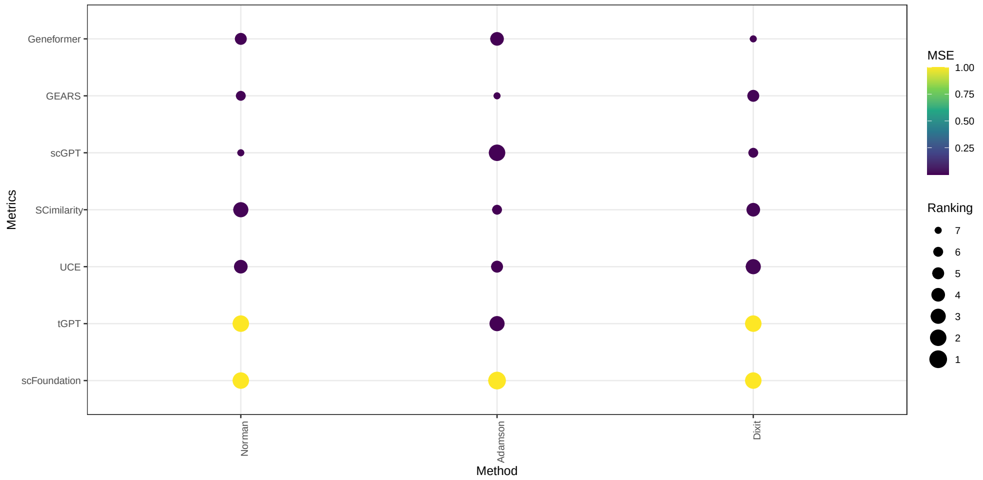


**Extended Data Fig. 22**: MSE across different models to evaluate the performances for perturbation prediction. The maximal value for displayed is 1 for better visualisation, while the true MSE of tGPT is larger than 1e5 for both Norman dataset and Dixit dataset. scFoundation meets the OOT error.

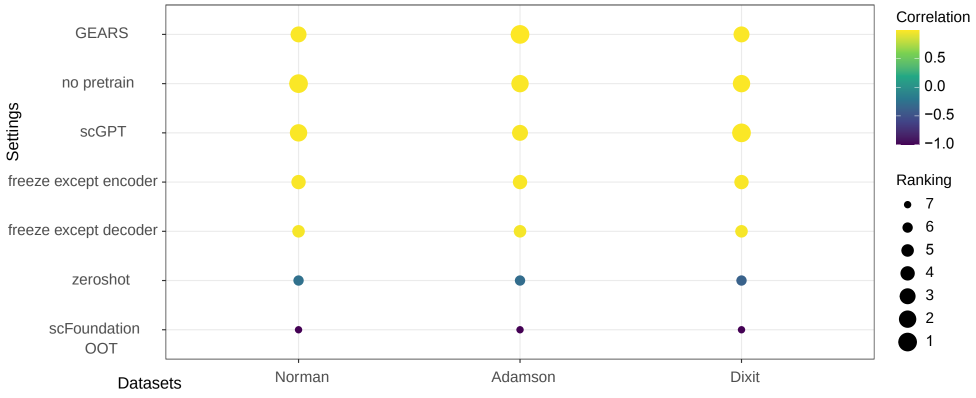


**Extended Data Fig. 23**: MPC across different fine-tuned-based models and settings of scGPT to evaluate the performances for perturbation prediction.

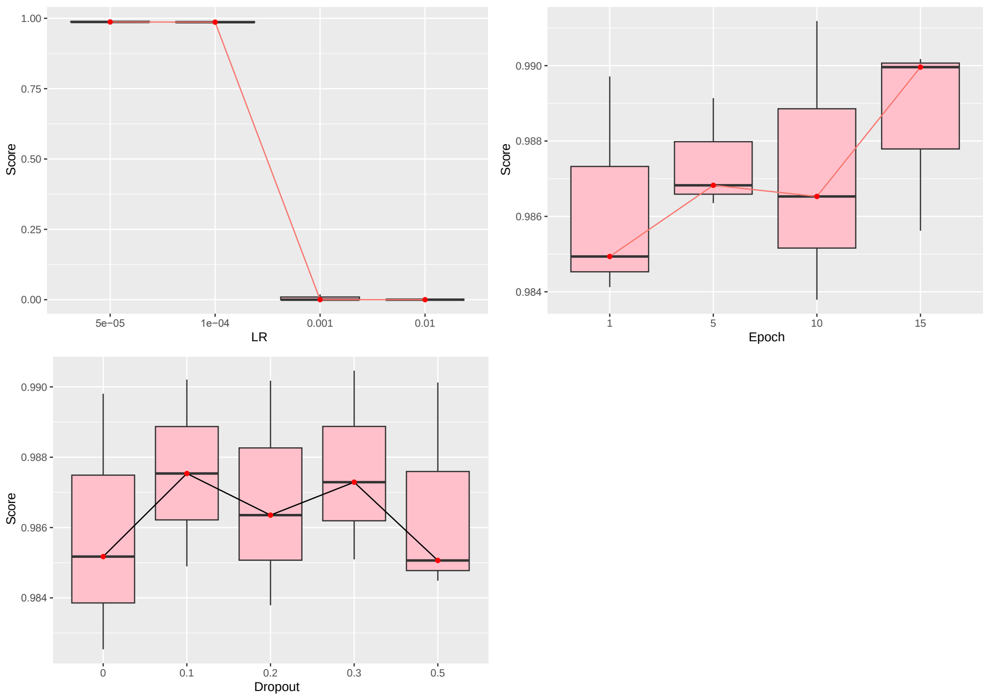


**Extended Data Fig. 24**: Tuning hyper-parameters for perturbation prediction. Sub-figures represent the score of scGPT under different hyper-parameters after training.

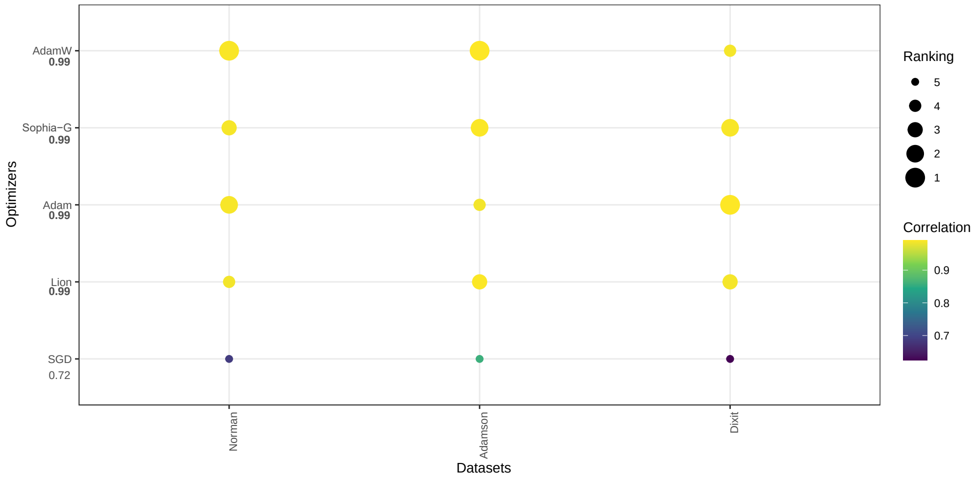


**Extended Data Fig. 25**: Benchmarking results of different optimizers for perturbation prediction.

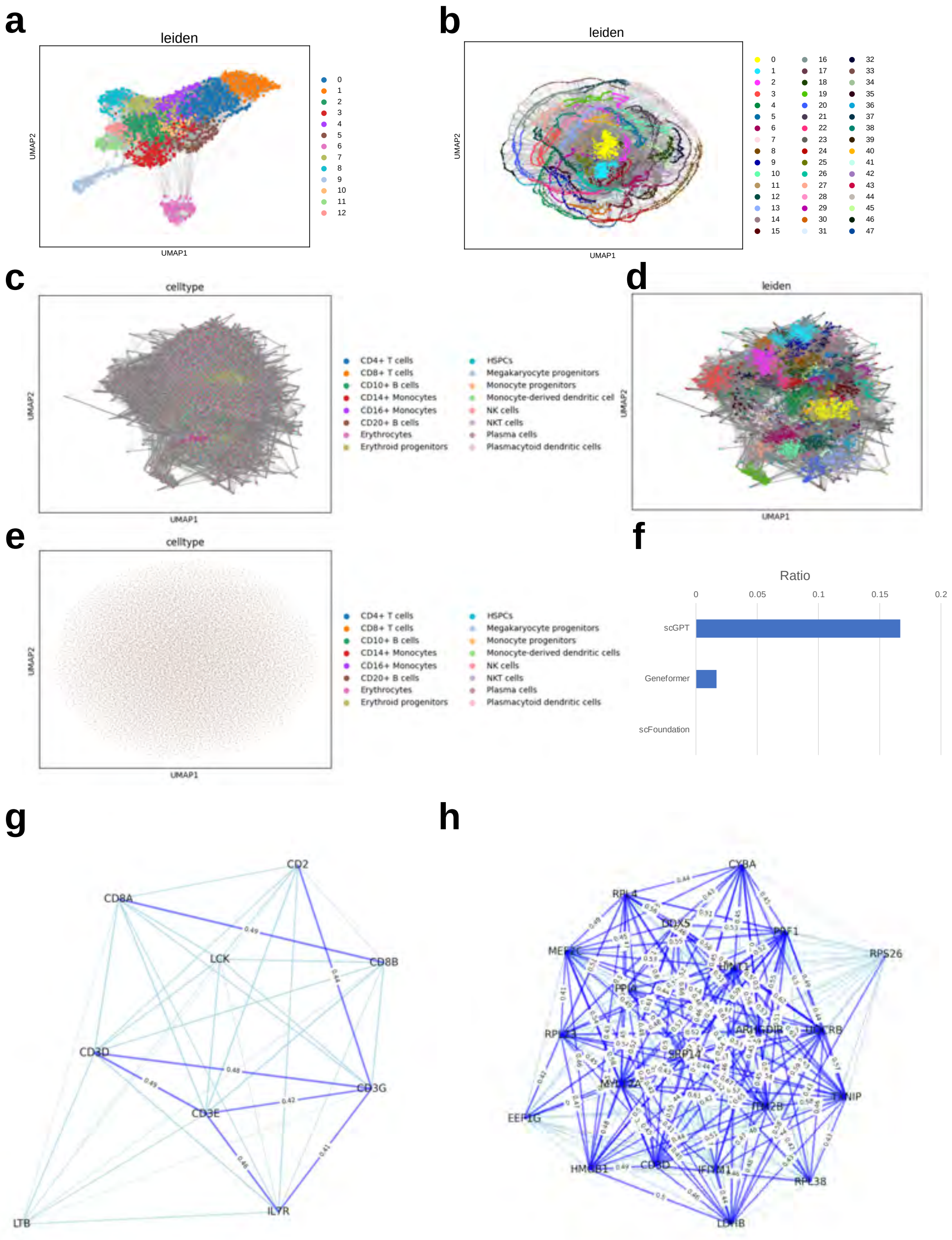


**Extended Data Fig. 26**: Examples of GCN inference results for the Immune Human Atlas (IHA) dataset. (a): Dataset-level gene embeddings from scGPT colored by Leiden clusters. (b): Dataset-level gene embeddings from Geneformer colored by Leiden clusters. (c): Cell-type-level gene embeddings from scGPT colored by the cell types. (d): Leiden cluster results based on the cell-type-level gene embeddings from scGPT. (e): Cell-type-level gene embeddings from Geneformer colored by the cell types. We omitted plotting the Leiden cluster results of Geneformer because of the plotting size limit. (f): Comparison of significant pathways ratio between scGPT and Geneformer for CD3-related gene sets. (g): An example of GCN for IHA based on scGPT. It is a network with CD3-related genes as major nodes. (h): An example of GCN for the IHA dataset based on Geneformer.

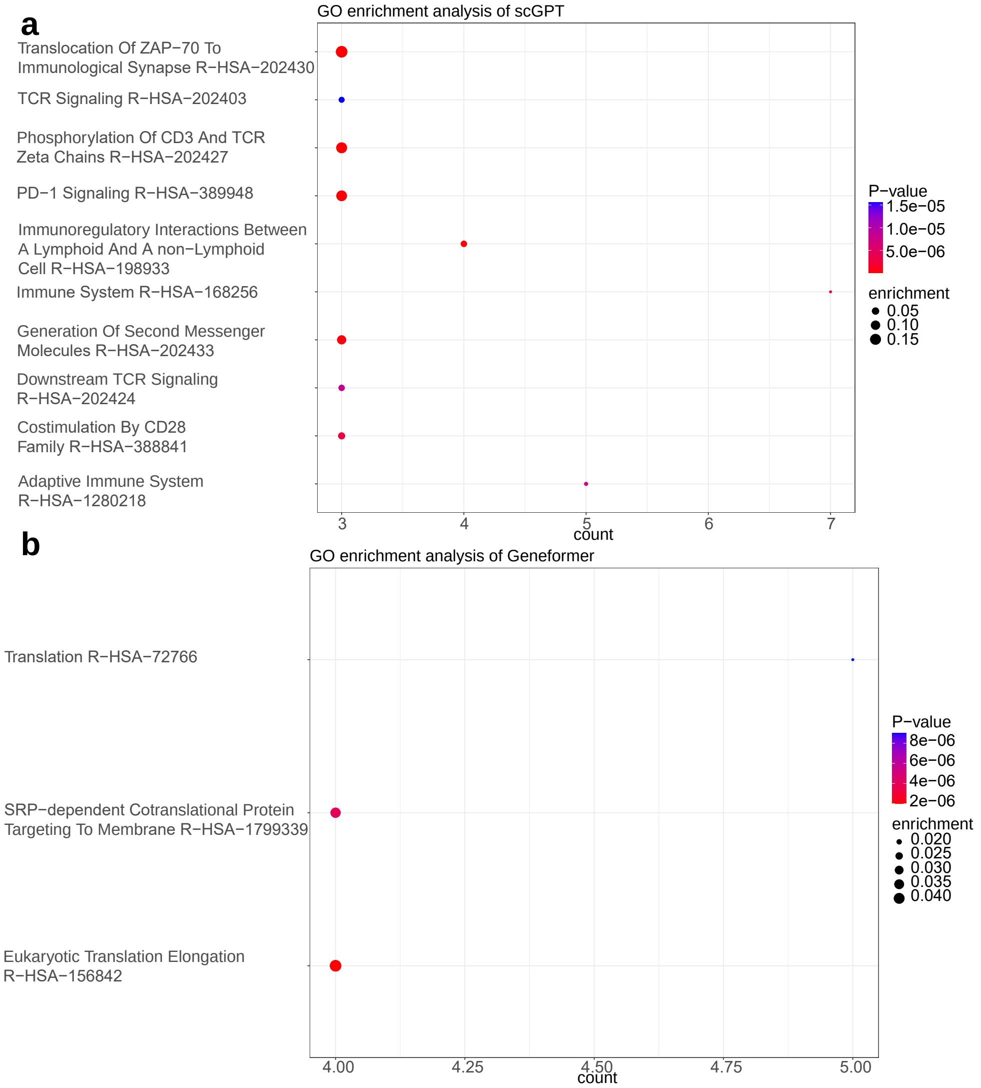


**Extended Data Fig. 27**: Detailed pathway information. (a) The pathway enrichment information from scGPT. (b) The pathway enrichment information from Geneformer.

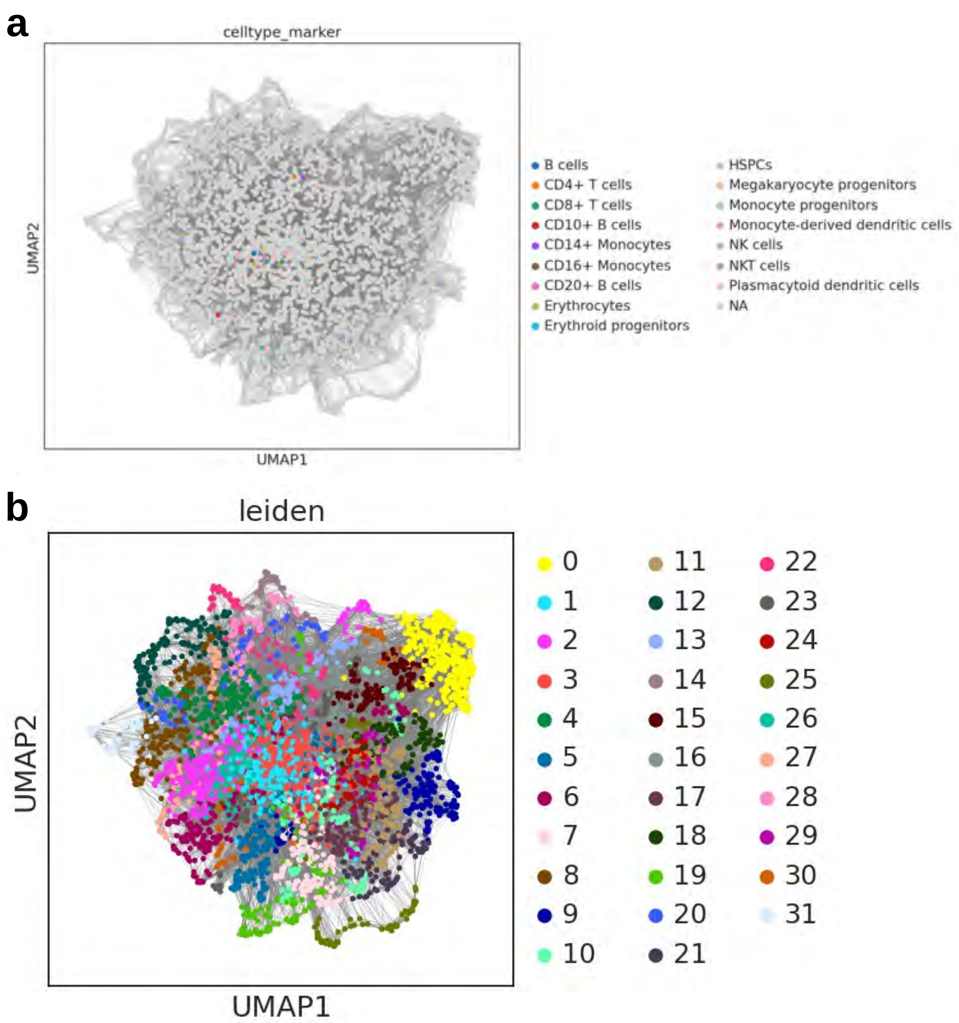


**Extended Data Fig. 28**: Examples of dataset-level GCNs from Geneformer based on HVGs. (a) Dataset-level gene embeddings from Geneformer colored by sources of marker genes. (b) Dataset-level gene embeddings from Geneformer colored by Leiden clusters.

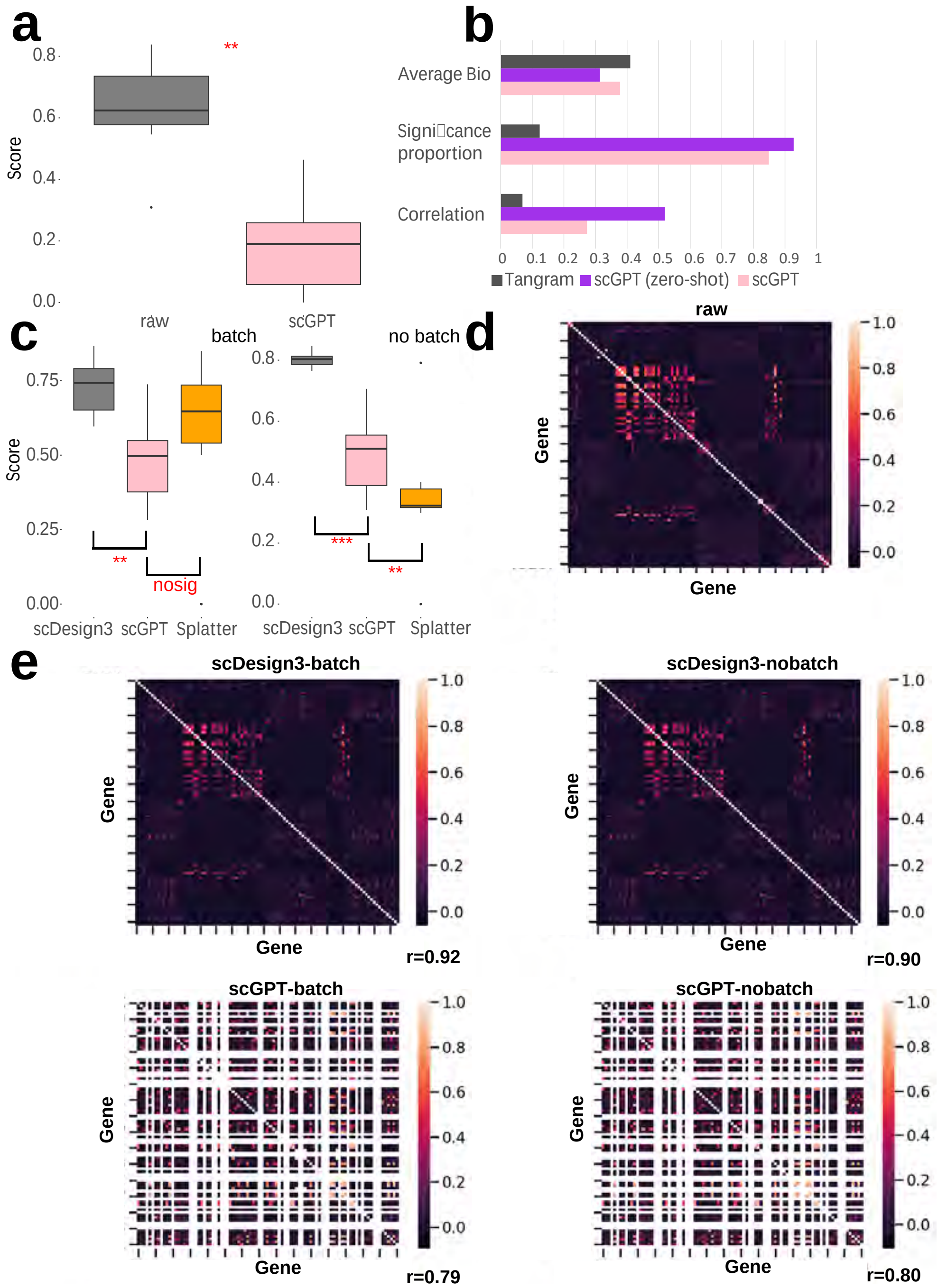


**Extended Data Fig. 29**: Experimental results of the Imputation task and the Simulation task. The significance level was computed based on paired Student's t-test. The number of stars represents the significance level ($* * * : p - value < 0.005, ** : p - value < 0.05$). (a): Comparison of the average bio score between the raw data and imputed data by scGPT in the scRNA-seq imputation task. (b): Comparison of the average bio score, average correlation score, and average significance level score among Tangram, scGPT, and scGPT (zero-shots) in spatial transcriptomics imputation task. (c): Comparison of the average bio score among scDesign3, Splatter, and scGPT for simulation. (d): Gene-gene correlation heatmap from the raw HumanPBMC dataset. We select the subset of the top 100 highly variable genes. (e): Comparison of different simulation methods by correlation. The heatmap represents the top 100 highly variable genes (for raw and scDesign3) or the subset of the top 100 highly variable genes (for scGPT) based on the HumanPBMC dataset. The correlation "r" represents the Pearson correlation between the gene correlation of raw data and the gene correlation of simulation data. The white section represents NaN values caused by the problematic model outputs.

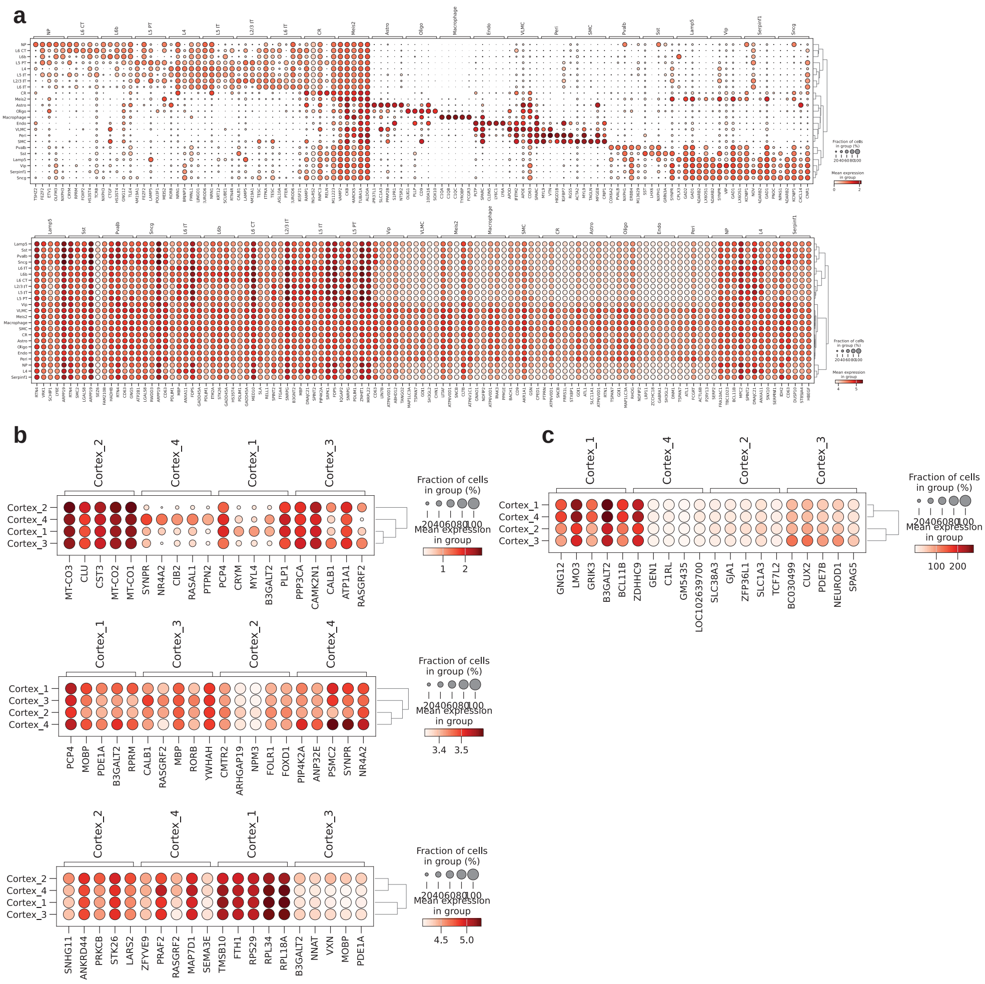


**Extended Data Fig. 30**: Deferentially expressed gene discovery based on results before imputation and after imputation. We used the Mouse scRNA-seq and the Mouse spatial transcriptomic datasets as examples. (a): Deferentially expressed genes by cell types for scRNA-seq data based on pre-imputation data (top) and post-imputation data (bottom). (b): Deferentially expressed genes by cluster types for spatial transcriptomic data based on pre-imputation data (top), post-imputation data based on zero-shot learning (middle), and post-imputation data based on fine-tuning (bottom). (c): Deferentially expressed genes by cluster types for spatial transcriptomic data based on Tangram.

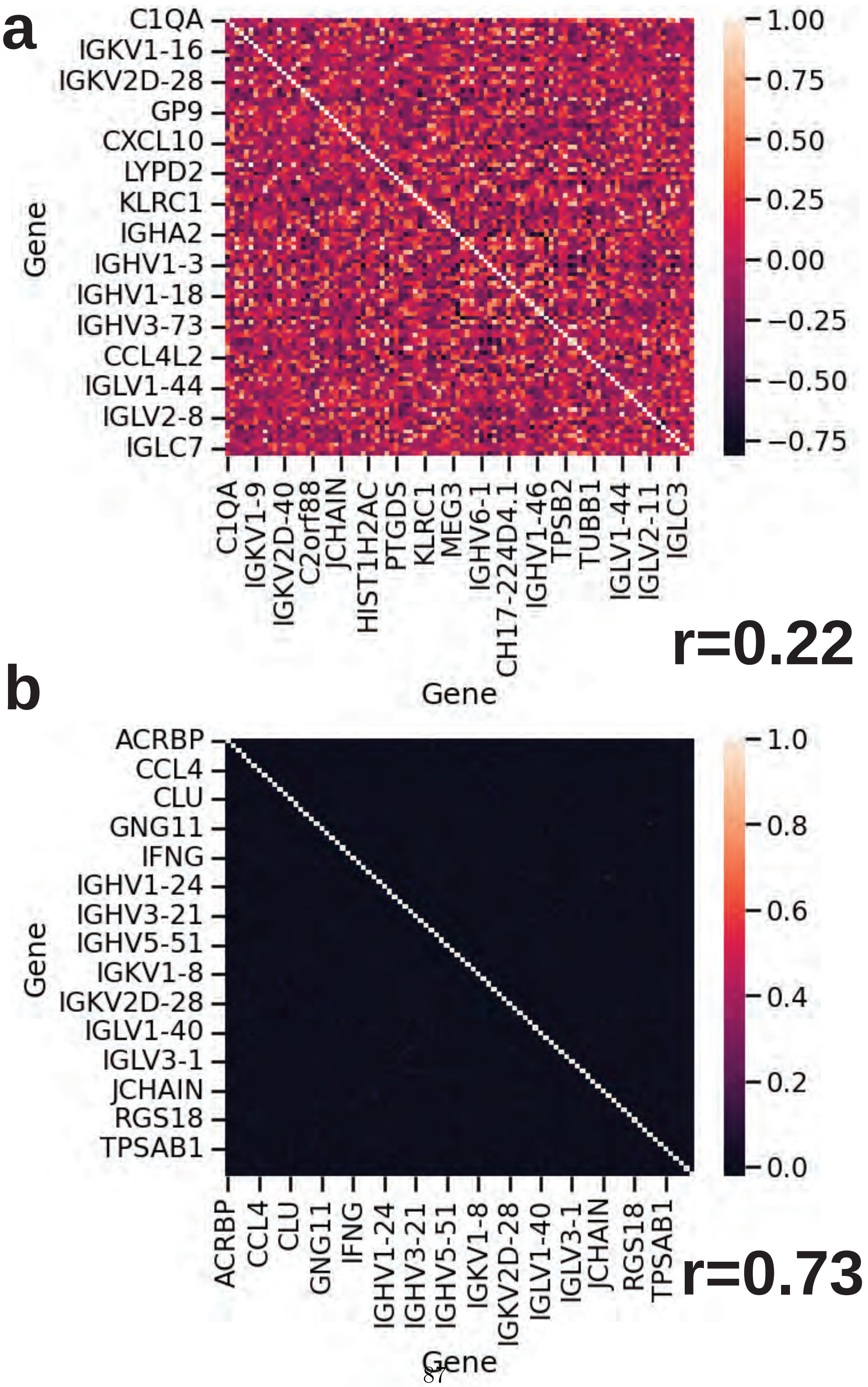




**Extended Data Fig. 31**: Heatmaps for the gene-gene correlation of datasets simulated by Splatter, using the HumanPBMC dataset as an example. The correlation "r" represents the Pearson correlation between the gene correlation of raw data and the gene correlation of simulation data. (a): The heatmap for the simulation dataset with batch effect. (b): The heatmap for the simulation dataset without batch effect.

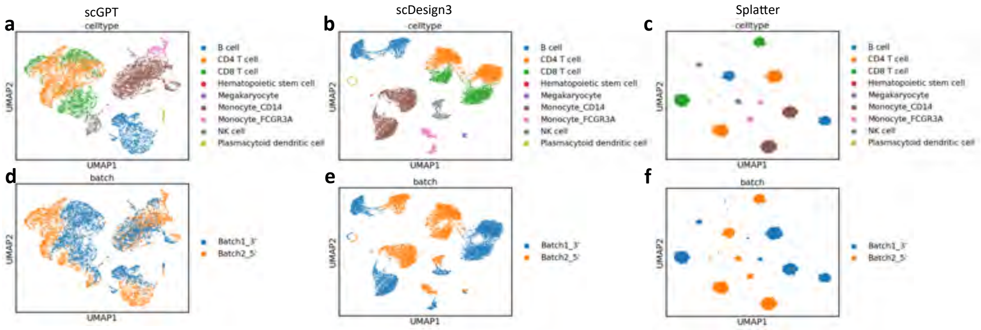


**Extended Data Fig. 32**: UMAPs for the simulation results with batch effect, using the HumanPBMC dataset as an example.

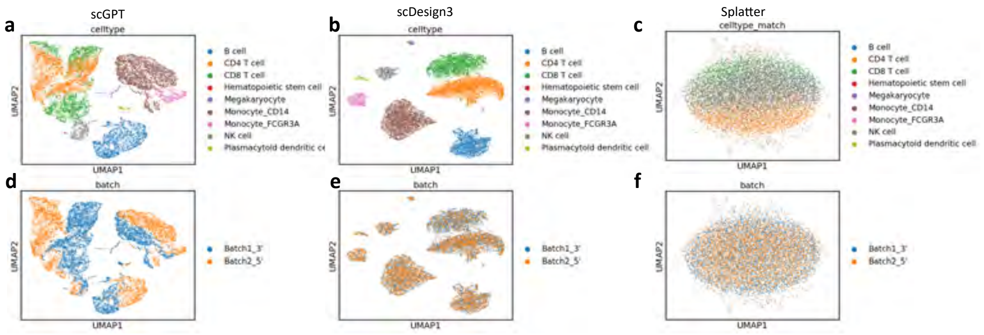


**Extended Data Fig. 33**: UMAPs for the simulation results without batch effect, using the HumanPBMC dataset as an example.

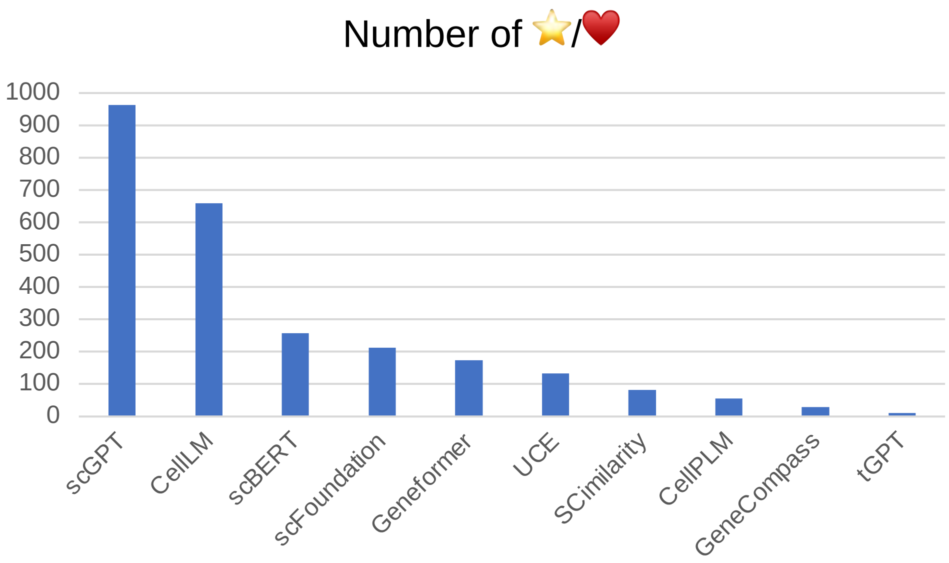


**Extended Data Fig. 34**: The number of stars in GitHub or likes in Huggingface for different single-cell FMs.

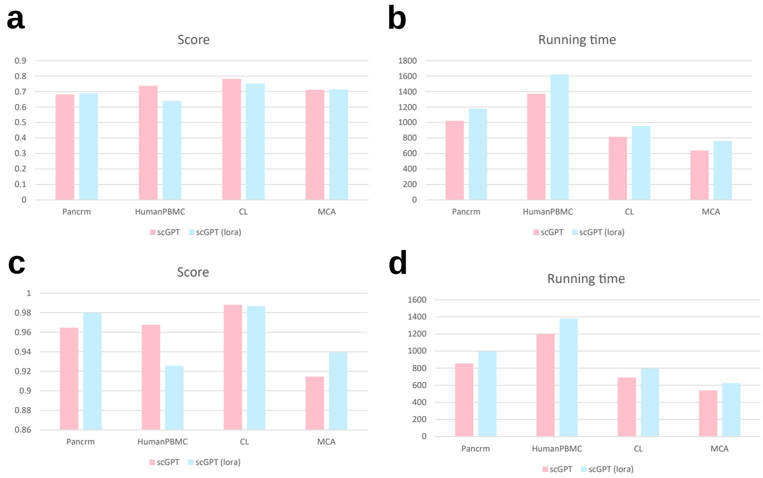


**Extended Data Fig. 35**: Preliminary results of parameter-efficient fine-tuning based on LoRA for Batch Effect Correction (the upper two figures) and Cell-type Annotation (the bottom two figures). (a): The comparison of scores for scGPT with/without LoRA. (b): The running time for scGPT with/without LoRA. (c): The comparison of scores for scGPT with/without LoRA. (d): The running time for scGPT with/without LoRA.

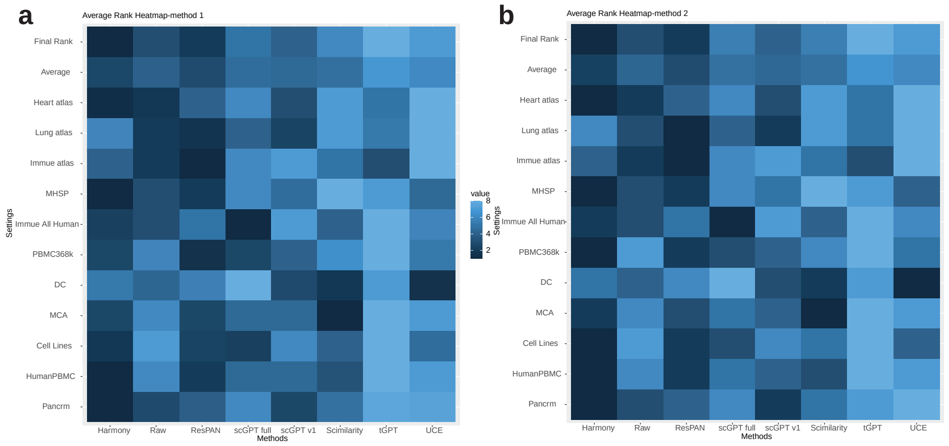


**Extended Data Fig. 36**: Heatmaps for the ranks of different models across different datasets. (a): The heatmap of ranks based on the first method, known as adjusting the weights and computing the ranks. (b): The heatmap of ranks based on the second method, known as using the default weights for $S_{bio}$ and $S_{batch}$.
