## Supplementary files. for "Evaluating the Utilities of Foundation Models in Single-cell Data Analysis": Supplementary file 1 (Keywords in AI).pdf

Glossary of common terms in Artificial Intelligence (AI) and Machine Learning (ML):

1. **Neural Network**: Neural network represents a system for computing inspired by the mechanism of biological neurons. The nodes in each layer of a neural network can communicate and generate the output. One neural network can be optimized by back-propagation.
2. **Deep learning:** Deep learning is a type of machine learning based on neural networks containing multiple layers. Each layer contains many nodes or perceptions. Transformer is a type of neural network. The input of neural networks can be either structured data (for example, biological networks) or unstructured data (for example, gene expression profiles and DNA sequence).
3. **Large language model (LLM):** LLM is a type of machine learning models containing multiple transformer layers. Typically, LLM is trained based on large-scale corpus datasets for Natural Language Processing (NLP) tasks. Moreover, the scale and size of LLM are very large and the training of LLM is based on large-scale datasets.
4. **Foundation model (FM):** FM is a type of machine learning models trained based on broad datasets and can support a diverse range of use cases. LLM is a type of FMs and we can have different FMs for different subjects or domains.
5. **Attention:** Attention is used in transformer layer to learn the representation of one sequence based on the relation of different positions of the same sequence. It is inspired by the visual attention mechanism of animals.
6. **Encoder-decoder architecture:** The encoder-decoder architecture is a common LLM/FM design. Such LLM contains one encoder component, which compresses high-dimensional data into low-dimensional representations (also known as embeddings); and one decoder component, which extends the low-dimensional representations to a high-dimensional space. Single-cell LLMs all follow this design, and the high-dimensional space represents the gene expression profile.
7. **Model size/model scale:** This term represents the number of parameters in an ML model.
8. **Parameters:** Parameters are variables used in LLM/FM training to maximize the performance.
9. **Hyper-parameters:** Hyper-parameters are settings for users to optimize the performance prior the model training, including learning rate, epochs, dropout rate, among others.
10. **Pre-training:** Pre-training is a technique used in LLM/FM using the self-supervised framework to train a LLM/FM without specific downstream application tasks. Based on the pre-trained model, we can train the model based on the datasets for specific tasks with small epochs (known as the fine-tuning process).
11. **Few-shot learning:** Few-shot learning means the fine-tuning process that only requires a few initial examples of the task.
12. **Zero-shot learning:** Without fine-tuning or other training processes, zero-shot learning directly applies the pre-trained model for specific tasks.
13. **Ablation tests:** Performing an ablation test means to evaluate the contribution of one component of the model by removing this component in the training process.
14. **Emergent ability:** Emergent ability is a property of pre-trained LLMs, which represents the ability of pre-trained LLMs that cannot be found or explained on small-scale models.
15. **LLM hallucination:** When the output of LLMs does not follow faithfulness or factualness, we declare that LLMs suffer from hallucinations. We can define the LLM hallucination for biological application as the output of LLMs that does not follow biological assumptions or is different from biological experiment results.

16. **Model attack:** The meaning of model attack is to verify the stability of the model or to find most effective attack approach by introducing human interference during the training of the model. An example is data poisoning, which means we manipulate the training datasets by injecting data with wrong information (for example, cells with wrong cell type labels) to control the behavior of model training.
