## Supplementary files. for "Evaluating the Utilities of Foundation Models in Single-cell Data Analysis": Supplementary file 5 (LLM structure).pdf

### Structure of scGPT:

```
TransformerModel(
  (encoder): GeneEncoder(
    (embedding): Embedding(36574, 512, padding_idx=36571)
    (enc_norm): LayerNorm((512,), eps=1e-05, elementwise_affine=True)
  )
  (value_encoder): ContinuousValueEncoder(
    (dropout): Dropout(p=0.2, inplace=False)
    (linear1): Linear(in_features=1, out_features=512, bias=True)
    (activation): ReLU()
    (linear2): Linear(in_features=512, out_features=512, bias=True)
    (norm): LayerNorm((512,), eps=1e-05, elementwise_affine=True)
  )
  (batch_encoder): BatchLabelEncoder(
    (embedding): Embedding(1, 512)
    (enc_norm): LayerNorm((512,), eps=1e-05, elementwise_affine=True)
  )
  (dsbn): DomainSpecificBatchNorm1d(
    (bns): ModuleList(
      (0): BatchNorm1d(512, eps=6.1e-05, momentum=0.1, affine=False, track_running_stats=True)
    )
  )
  (transformer_encoder): TransformerEncoder(
    (layers): ModuleList(
      (0): FlashTransformerEncoderLayer(
        (self_attn): FlashMHA(
          (Wqkv): Linear(in_features=512, out_features=1536, bias=True)
          (inner_attn): FlashAttention()
          (out_proj): Linear(in_features=512, out_features=512, bias=True)
        )
        (linear1): Linear(in_features=512, out_features=512, bias=True)
        (dropout): Dropout(p=0.2, inplace=False)
        (linear2): Linear(in_features=512, out_features=512, bias=True)
        (norm1): LayerNorm((512,), eps=1e-05, elementwise_affine=True)
        (norm2): LayerNorm((512,), eps=1e-05, elementwise_affine=True)
        (dropout1): Dropout(p=0.2, inplace=False)
        (dropout2): Dropout(p=0.2, inplace=False)
      )
      (1): FlashTransformerEncoderLayer(
        (self_attn): FlashMHA(
          (Wqkv): Linear(in_features=512, out_features=1536, bias=True)
          (inner_attn): FlashAttention()
          (out_proj): Linear(in_features=512, out_features=512, bias=True)
        )
        (linear1): Linear(in_features=512, out_features=512, bias=True)
        (dropout): Dropout(p=0.2, inplace=False)
        (linear2): Linear(in_features=512, out_features=512, bias=True)
        (norm1): LayerNorm((512,), eps=1e-05, elementwise_affine=True)
        (norm2): LayerNorm((512,), eps=1e-05, elementwise_affine=True)
        (dropout1): Dropout(p=0.2, inplace=False)
        (dropout2): Dropout(p=0.2, inplace=False)
      )
      (2): FlashTransformerEncoderLayer(
```

```
    (self_attn): FlashMHA(
      (Wqkv): Linear(in_features=512, out_features=1536, bias=True)
      (inner_attn): FlashAttention()
      (out_proj): Linear(in_features=512, out_features=512, bias=True)
    )
    (linear1): Linear(in_features=512, out_features=512, bias=True)
    (dropout): Dropout(p=0.2, inplace=False)
    (linear2): Linear(in_features=512, out_features=512, bias=True)
    (norm1): LayerNorm((512,), eps=1e-05, elementwise_affine=True)
    (norm2): LayerNorm((512,), eps=1e-05, elementwise_affine=True)
    (dropout1): Dropout(p=0.2, inplace=False)
    (dropout2): Dropout(p=0.2, inplace=False)
  )
  (3): FlashTransformerEncoderLayer(
    (self_attn): FlashMHA(
      (Wqkv): Linear(in_features=512, out_features=1536, bias=True)
      (inner_attn): FlashAttention()
      (out_proj): Linear(in_features=512, out_features=512, bias=True)
    )
    (linear1): Linear(in_features=512, out_features=512, bias=True)
    (dropout): Dropout(p=0.2, inplace=False)
    (linear2): Linear(in_features=512, out_features=512, bias=True)
    (norm1): LayerNorm((512,), eps=1e-05, elementwise_affine=True)
    (norm2): LayerNorm((512,), eps=1e-05, elementwise_affine=True)
    (dropout1): Dropout(p=0.2, inplace=False)
    (dropout2): Dropout(p=0.2, inplace=False)
  )
  (4): FlashTransformerEncoderLayer(
    (self_attn): FlashMHA(
      (Wqkv): Linear(in_features=512, out_features=1536, bias=True)
      (inner_attn): FlashAttention()
      (out_proj): Linear(in_features=512, out_features=512, bias=True)
    )
    (linear1): Linear(in_features=512, out_features=512, bias=True)
    (dropout): Dropout(p=0.2, inplace=False)
    (linear2): Linear(in_features=512, out_features=512, bias=True)
    (norm1): LayerNorm((512,), eps=1e-05, elementwise_affine=True)
    (norm2): LayerNorm((512,), eps=1e-05, elementwise_affine=True)
    (dropout1): Dropout(p=0.2, inplace=False)
    (dropout2): Dropout(p=0.2, inplace=False)
  )
  (5): FlashTransformerEncoderLayer(
    (self_attn): FlashMHA(
      (Wqkv): Linear(in_features=512, out_features=1536, bias=True)
      (inner_attn): FlashAttention()
      (out_proj): Linear(in_features=512, out_features=512, bias=True)
    )
    (linear1): Linear(in_features=512, out_features=512, bias=True)
    (dropout): Dropout(p=0.2, inplace=False)
    (linear2): Linear(in_features=512, out_features=512, bias=True)
    (norm1): LayerNorm((512,), eps=1e-05, elementwise_affine=True)
    (norm2): LayerNorm((512,), eps=1e-05, elementwise_affine=True)
    (dropout1): Dropout(p=0.2, inplace=False)
```

```
    (dropout2): Dropout(p=0.2, inplace=False)
  )
  (6): FlashTransformerEncoderLayer(
    (self_attn): FlashMHA(
      (Wqkv): Linear(in_features=512, out_features=1536, bias=True)
      (inner_attn): FlashAttention()
      (out_proj): Linear(in_features=512, out_features=512, bias=True)
    )
    (linear1): Linear(in_features=512, out_features=512, bias=True)
    (dropout): Dropout(p=0.2, inplace=False)
    (linear2): Linear(in_features=512, out_features=512, bias=True)
    (norm1): LayerNorm((512,), eps=1e-05, elementwise_affine=True)
    (norm2): LayerNorm((512,), eps=1e-05, elementwise_affine=True)
    (dropout1): Dropout(p=0.2, inplace=False)
    (dropout2): Dropout(p=0.2, inplace=False)
  )
  (7): FlashTransformerEncoderLayer(
    (self_attn): FlashMHA(
      (Wqkv): Linear(in_features=512, out_features=1536, bias=True)
      (inner_attn): FlashAttention()
      (out_proj): Linear(in_features=512, out_features=512, bias=True)
    )
    (linear1): Linear(in_features=512, out_features=512, bias=True)
    (dropout): Dropout(p=0.2, inplace=False)
    (linear2): Linear(in_features=512, out_features=512, bias=True)
    (norm1): LayerNorm((512,), eps=1e-05, elementwise_affine=True)
    (norm2): LayerNorm((512,), eps=1e-05, elementwise_affine=True)
    (dropout1): Dropout(p=0.2, inplace=False)
    (dropout2): Dropout(p=0.2, inplace=False)
  )
  (8): FlashTransformerEncoderLayer(
    (self_attn): FlashMHA(
      (Wqkv): Linear(in_features=512, out_features=1536, bias=True)
      (inner_attn): FlashAttention()
      (out_proj): Linear(in_features=512, out_features=512, bias=True)
    )
    (linear1): Linear(in_features=512, out_features=512, bias=True)
    (dropout): Dropout(p=0.2, inplace=False)
    (linear2): Linear(in_features=512, out_features=512, bias=True)
    (norm1): LayerNorm((512,), eps=1e-05, elementwise_affine=True)
    (norm2): LayerNorm((512,), eps=1e-05, elementwise_affine=True)
    (dropout1): Dropout(p=0.2, inplace=False)
    (dropout2): Dropout(p=0.2, inplace=False)
  )
  (9): FlashTransformerEncoderLayer(
    (self_attn): FlashMHA(
      (Wqkv): Linear(in_features=512, out_features=1536, bias=True)
      (inner_attn): FlashAttention()
      (out_proj): Linear(in_features=512, out_features=512, bias=True)
    )
    (linear1): Linear(in_features=512, out_features=512, bias=True)
    (dropout): Dropout(p=0.2, inplace=False)
    (linear2): Linear(in_features=512, out_features=512, bias=True)
```

```
      (norm1): LayerNorm((512,), eps=1e-05, elementwise_affine=True)
      (norm2): LayerNorm((512,), eps=1e-05, elementwise_affine=True)
      (dropout1): Dropout(p=0.2, inplace=False)
      (dropout2): Dropout(p=0.2, inplace=False)
    )
    (10): FlashTransformerEncoderLayer(
      (self_attn): FlashMHA(
        (Wqkv): Linear(in_features=512, out_features=1536, bias=True)
        (inner_attn): FlashAttention()
        (out_proj): Linear(in_features=512, out_features=512, bias=True)
      )
      (linear1): Linear(in_features=512, out_features=512, bias=True)
      (dropout): Dropout(p=0.2, inplace=False)
      (linear2): Linear(in_features=512, out_features=512, bias=True)
      (norm1): LayerNorm((512,), eps=1e-05, elementwise_affine=True)
      (norm2): LayerNorm((512,), eps=1e-05, elementwise_affine=True)
      (dropout1): Dropout(p=0.2, inplace=False)
      (dropout2): Dropout(p=0.2, inplace=False)
    )
    (11): FlashTransformerEncoderLayer(
      (self_attn): FlashMHA(
        (Wqkv): Linear(in_features=512, out_features=1536, bias=True)
        (inner_attn): FlashAttention()
        (out_proj): Linear(in_features=512, out_features=512, bias=True)
      )
      (linear1): Linear(in_features=512, out_features=512, bias=True)
      (dropout): Dropout(p=0.2, inplace=False)
      (linear2): Linear(in_features=512, out_features=512, bias=True)
      (norm1): LayerNorm((512,), eps=1e-05, elementwise_affine=True)
      (norm2): LayerNorm((512,), eps=1e-05, elementwise_affine=True)
      (dropout1): Dropout(p=0.2, inplace=False)
      (dropout2): Dropout(p=0.2, inplace=False)
    )
  )
)
(decoder): ExprDecoder(
  (fc): Sequential(
    (0): Linear(in_features=1024, out_features=512, bias=True)
    (1): LeakyReLU(negative_slope=0.01)
    (2): Linear(in_features=512, out_features=512, bias=True)
    (3): LeakyReLU(negative_slope=0.01)
    (4): Linear(in_features=512, out_features=1, bias=True)
  )
  (zero_logit): Sequential(
    (0): Linear(in_features=1024, out_features=512, bias=True)
    (1): LeakyReLU(negative_slope=0.01)
    (2): Linear(in_features=512, out_features=512, bias=True)
    (3): LeakyReLU(negative_slope=0.01)
    (4): Linear(in_features=512, out_features=1, bias=True)
  )
)
(cls_decoder): ClsDecoder(
  (_decoder): ModuleList(
```

```
      (0): Linear(in_features=512, out_features=512, bias=True)
      (1): ReLU()
      (2): LayerNorm((512,), eps=1e-05, elementwise_affine=True)
      (3): Linear(in_features=512, out_features=512, bias=True)
      (4): ReLU()
      (5): LayerNorm((512,), eps=1e-05, elementwise_affine=True)
    )
    (out_layer): Linear(in_features=512, out_features=1, bias=True)
  )
  (mvc_decoder): MVCDecoder(
    (gene2query): Linear(in_features=512, out_features=512, bias=True)
    (query_activation): Sigmoid()
    (W): Linear(in_features=512, out_features=1024, bias=False)
    (W_zero_logit): Linear(in_features=512, out_features=1024, bias=True)
  )
  (grad_reverse_discriminator): AdversarialDiscriminator(
    (_decoder): ModuleList(
      (0): Linear(in_features=512, out_features=512, bias=True)
      (1): LeakyReLU(negative_slope=0.01)
      (2): LayerNorm((512,), eps=1e-05, elementwise_affine=True)
      (3): Linear(in_features=512, out_features=512, bias=True)
      (4): LeakyReLU(negative_slope=0.01)
      (5): LayerNorm((512,), eps=1e-05, elementwise_affine=True)
    )
    (out_layer): Linear(in_features=512, out_features=1, bias=True)
  )
  (sim): Similarity(
    (cos): CosineSimilarity()
  )
  (creterion_cce): CrossEntropyLoss()
)
```

### Structure of scBERT:

```
PerformerLM(
  (token_emb): Embedding(7, 200)
  (pos_emb): Gene2VecPositionalEmbedding(
    (emb): Embedding(16907, 200)
  )
  (layer_pos_emb): Always()
  (dropout): Dropout(p=0.0, inplace=False)
  (performer): Performer(
    (net): SequentialSequence(
      (layers): ModuleList(
        (0-5): 6 x ModuleList(
          (0): PreLayerNorm(
            (norm): LayerNorm((200,), eps=1e-05, elementwise_affine=True)
            (fn): SelfAttention(
              (fast_attention): FastAttention(
                (kernel_fn): ReLU()
              )
              (to_q): Linear(in_features=200, out_features=640, bias=False)
              (to_k): Linear(in_features=200, out_features=640, bias=False)
              (to_v): Linear(in_features=200, out_features=640, bias=False)
              (to_out): Linear(in_features=640, out_features=200, bias=True)
              (dropout): Dropout(p=0.0, inplace=False)
            )
          )
          (1): PreLayerNorm(
            (norm): LayerNorm((200,), eps=1e-05, elementwise_affine=True)
            (fn): Chunk(
              (fn): FeedForward(
                (w1): Linear(in_features=200, out_features=800, bias=True)
                (act): GELU(approximate='none')
                (dropout): Dropout(p=0.0, inplace=False)
                (w2): Linear(in_features=800, out_features=200, bias=True)
              )
            )
          )
        )
      )
    )
  )
  (norm): LayerNorm((200,), eps=1e-05, elementwise_affine=True)
  (to_out): Linear(in_features=200, out_features=7, bias=True)
)
```

### Structure of Geneformer:

```
BertForTokenClassification(
  (bert): BertModel(
    (embeddings): BertEmbeddings(
      (word_embeddings): Embedding(25426, 256, padding_idx=0)
      (position_embeddings): Embedding(2048, 256)
      (token_type_embeddings): Embedding(2, 256)
      (LayerNorm): LayerNorm((256,), eps=1e-12, elementwise_affine=True)
      (dropout): Dropout(p=0.02, inplace=False)
    )
    (encoder): BertEncoder(
      (layer): ModuleList(
        (0-5): 6 x BertLayer(
          (attention): BertAttention(
            (self): BertSelfAttention(
              (query): Linear(in_features=256, out_features=256, bias=True)
              (key): Linear(in_features=256, out_features=256, bias=True)
              (value): Linear(in_features=256, out_features=256, bias=True)
              (dropout): Dropout(p=0.02, inplace=False)
            )
            (output): BertSelfOutput(
              (dense): Linear(in_features=256, out_features=256, bias=True)
              (LayerNorm): LayerNorm((256,), eps=1e-12, elementwise_affine=True)
              (dropout): Dropout(p=0.02, inplace=False)
            )
          )
          (intermediate): BertIntermediate(
            (dense): Linear(in_features=256, out_features=512, bias=True)
            (intermediate_act_fn): ReLU()
          )
          (output): BertOutput(
            (dense): Linear(in_features=512, out_features=256, bias=True)
            (LayerNorm): LayerNorm((256,), eps=1e-12, elementwise_affine=True)
            (dropout): Dropout(p=0.02, inplace=False)
          )
        )
      )
    )
  )
  (dropout): Dropout(p=0.02, inplace=False)
  (classifier): Linear(in_features=256, out_features=2, bias=True)
)
```

### Structure of CellLM:

```
CTCModel(
  (encoder): PerformerLM_CellLM(
    (token_emb): Embedding(8, 512)
    (pos_emb): Gene2VecPositionalEmbedding(
      (emb): Embedding(19381, 512)
    )
    (layer_pos_emb): Always()
    (dropout): Dropout(p=0.0, inplace=False)
    (performer): Performer(
      (net): SequentialSequence(
        (layers): ModuleList(
          (0-9): 10 x ModuleList(
            (0): PreLayerNorm(
              (norm): LayerNorm((512,), eps=1e-05, elementwise_affine=True)
              (fn): SelfAttention(
                (fast_attention): FastAttention(
                  (kernel_fn): ReLU()
                )
                (to_q): Linear(in_features=512, out_features=1024, bias=False)
                (to_k): Linear(in_features=512, out_features=1024, bias=False)
                (to_v): Linear(in_features=512, out_features=1024, bias=False)
                (to_out): Linear(in_features=1024, out_features=512, bias=True)
                (dropout): Dropout(p=0.0, inplace=False)
              )
            )
            (1): PreLayerNorm(
              (norm): LayerNorm((512,), eps=1e-05, elementwise_affine=True)
              (fn): Chunk(
                (fn): FeedForward(
                  (w1): Linear(in_features=512, out_features=2048, bias=True)
                  (act): GELU(approximate='none')
                  (dropout): Dropout(p=0.0, inplace=False)
                  (w2): Linear(in_features=2048, out_features=512, bias=True)
                )
              )
            )
          )
        )
      )
    )
    (norm): LayerNorm((512,), eps=1e-05, elementwise_affine=True)
    (to_out): Linear(in_features=512, out_features=8, bias=True)
  )
  (conv): Conv2d(1, 1, kernel_size=(1, 512), stride=(1, 1))
  (pred_head): Sequential(
    (0): ReLU()
    (1): Linear(in_features=6000, out_features=512, bias=True)
    (2): ReLU()
    (3): Dropout(p=0, inplace=False)
    (4): Linear(in_features=512, out_features=128, bias=True)
    (5): ReLU()
    (6): Dropout(p=0, inplace=False)
```

```
    (7): Linear(in_features=128, out_features=4, bias=True)
  )
)
```

### Structure of CellPLM:

```
OmicsFormer(
  (embedder): OmicsEmbeddingLayer(
    (act): ReLU()
    (norm0): GroupNorm(4, 1024, eps=1e-05, affine=True)
    (dropout): Dropout(p=0.5, inplace=False)
    (extra_linear): Sequential(
      (0): Linear(in_features=1024, out_features=1024, bias=True)
      (1): ReLU()
      (2): Dropout(p=0.5, inplace=False)
      (3): GroupNorm(4, 1024, eps=1e-05, affine=True)
    )
    (pe_enc): Sinusoidal2dPE(
      (pe_enc): Embedding(10000, 1024)
    )
    (feat_enc): OmicsEmbedder()
  )
  (mask_model): MaskBuilder()
  (encoder): TransformerEncoder(
    (layers): ModuleList(
      (0): FlowformerLayer(
        (self_attn): Flow_Attention(
          (query_projection): Linear(in_features=1024, out_features=1024, bias=True)
          (key_projection): Linear(in_features=1024, out_features=1024, bias=True)
          (value_projection): Linear(in_features=1024, out_features=1024, bias=True)
          (out_projection): Linear(in_features=1024, out_features=1024, bias=True)
          (dropout): Dropout(p=0.01, inplace=False)
        )
        (_ff_block): Sequential(
          (0): Linear(in_features=1024, out_features=2048, bias=True)
          (1): GELU(approximate='none')
          (2): Dropout(p=0.5, inplace=False)
          (3): Linear(in_features=2048, out_features=1024, bias=True)
          (4): Dropout(p=0.5, inplace=False)
        )
        (dropout1): Dropout(p=0.5, inplace=False)
        (norm1): LayerNorm((1024,), eps=1e-05, elementwise_affine=True)
        (norm2): LayerNorm((1024,), eps=1e-05, elementwise_affine=True)
      )
      (1): FlowformerLayer(
        (self_attn): Flow_Attention(
          (query_projection): Linear(in_features=1024, out_features=1024, bias=True)
          (key_projection): Linear(in_features=1024, out_features=1024, bias=True)
          (value_projection): Linear(in_features=1024, out_features=1024, bias=True)
          (out_projection): Linear(in_features=1024, out_features=1024, bias=True)
          (dropout): Dropout(p=0.01, inplace=False)
        )
        (_ff_block): Sequential(
          (0): Linear(in_features=1024, out_features=2048, bias=True)
          (1): GELU(approximate='none')
          (2): Dropout(p=0.5, inplace=False)
          (3): Linear(in_features=2048, out_features=1024, bias=True)
          (4): Dropout(p=0.5, inplace=False)
```

```
      )
      (dropout1): Dropout(p=0.5, inplace=False)
      (norm1): LayerNorm((1024,), eps=1e-05, elementwise_affine=True)
      (norm2): LayerNorm((1024,), eps=1e-05, elementwise_affine=True)
    )
    (2): FlowformerLayer(
      (self_attn): Flow_Attention(
        (query_projection): Linear(in_features=1024, out_features=1024, bias=True)
        (key_projection): Linear(in_features=1024, out_features=1024, bias=True)
        (value_projection): Linear(in_features=1024, out_features=1024, bias=True)
        (out_projection): Linear(in_features=1024, out_features=1024, bias=True)
        (dropout): Dropout(p=0.01, inplace=False)
      )
      (_ff_block): Sequential(
        (0): Linear(in_features=1024, out_features=2048, bias=True)
        (1): GELU(approximate='none')
        (2): Dropout(p=0.5, inplace=False)
        (3): Linear(in_features=2048, out_features=1024, bias=True)
        (4): Dropout(p=0.5, inplace=False)
      )
      (dropout1): Dropout(p=0.5, inplace=False)
      (norm1): LayerNorm((1024,), eps=1e-05, elementwise_affine=True)
      (norm2): LayerNorm((1024,), eps=1e-05, elementwise_affine=True)
    )
    (3): FlowformerLayer(
      (self_attn): Flow_Attention(
        (query_projection): Linear(in_features=1024, out_features=1024, bias=True)
        (key_projection): Linear(in_features=1024, out_features=1024, bias=True)
        (value_projection): Linear(in_features=1024, out_features=1024, bias=True)
        (out_projection): Linear(in_features=1024, out_features=1024, bias=True)
        (dropout): Dropout(p=0.01, inplace=False)
      )
      (_ff_block): Sequential(
        (0): Linear(in_features=1024, out_features=2048, bias=True)
        (1): GELU(approximate='none')
        (2): Dropout(p=0.5, inplace=False)
        (3): Linear(in_features=2048, out_features=1024, bias=True)
        (4): Dropout(p=0.5, inplace=False)
      )
      (dropout1): Dropout(p=0.5, inplace=False)
      (norm1): LayerNorm((1024,), eps=1e-05, elementwise_affine=True)
      (norm2): LayerNorm((1024,), eps=1e-05, elementwise_affine=True)
    )
  )
)
(latent): LatentModel(
  (layers): ModuleList(
    (0): PlaceholderLayer()
    (1): MergeLatentLayer(
      (lat_2lat): Sequential(
        (0): Linear(in_features=1024, out_features=512, bias=True)
      )
    )
```

```
    )
  )
  (head): AnnotationHead(
    (ce_loss): CrossEntropyLoss()
    (mlp): Sequential(
      (0): Dropout(p=0.5, inplace=False)
      (1): Linear(in_features=512, out_features=14, bias=True)
    )
  )
  (pre_latent_norm): PreLatentNorm(
    (norm): LayerNorm((1024,), eps=1e-05, elementwise_affine=True)
  )
)
```

### Structure of SCimilarity:

```
Encoder(
  (network): ModuleList(
    (0): Sequential(
      (0): Dropout(p=0.4, inplace=False)
      (1): Linear(in_features=28231, out_features=1024, bias=True)
      (2): BatchNorm1d(1024, eps=1e-05, momentum=0.1, affine=True, track_running_stats=True)
      (3): PReLU(num_parameters=1)
    )
    (1-2): 2 x Sequential(
      (0): Dropout(p=0.5, inplace=False)
      (1): Linear(in_features=1024, out_features=1024, bias=True)
      (2): BatchNorm1d(1024, eps=1e-05, momentum=0.1, affine=True, track_running_stats=True)
      (3): PReLU(num_parameters=1)
    )
    (3): Linear(in_features=1024, out_features=128, bias=True)
  )
)
```

[8]:

### Structure of UCE:

```
TransformerModel(
  (pos_encoder): PositionalEncoding(
    (dropout): Dropout(p=0.05, inplace=False)
  )
  (encoder): Sequential(
    (0): Linear(in_features=5120, out_features=1280, bias=True)
    (1): GELU(approximate='none')
    (2): LayerNorm((1280,), eps=1e-05, elementwise_affine=True)
  )
  (transformer_encoder): TransformerEncoder(
    (layers): ModuleList(
      (0-3): 4 x TransformerEncoderLayer(
        (self_attn): MultiheadAttention(
          (out_proj): NonDynamicallyQuantizableLinear(in_features=1280, out_features=1280, bias=Tru
e)
        )
        (linear1): Linear(in_features=1280, out_features=5120, bias=True)
        (dropout): Dropout(p=0.05, inplace=False)
        (linear2): Linear(in_features=5120, out_features=1280, bias=True)
        (norm1): LayerNorm((1280,), eps=1e-05, elementwise_affine=True)
        (norm2): LayerNorm((1280,), eps=1e-05, elementwise_affine=True)
        (dropout1): Dropout(p=0.05, inplace=False)
        (dropout2): Dropout(p=0.05, inplace=False)
      )
    )
  )
  (decoder): Sequential(
    (0): Sequential(
      (0): Linear(in_features=1280, out_features=1024, bias=True)
      (1): LayerNorm((1024,), eps=1e-05, elementwise_affine=True)
      (2): GELU(approximate='none')
      (3): Dropout(p=0.05, inplace=False)
    )
    (1): Sequential(
      (0): Linear(in_features=1024, out_features=1280, bias=True)
      (1): LayerNorm((1280,), eps=1e-05, elementwise_affine=True)
      (2): GELU(approximate='none')
      (3): Dropout(p=0.05, inplace=False)
    )
    (2): Sequential(
      (0): Linear(in_features=1280, out_features=1280, bias=True)
      (1): LayerNorm((1280,), eps=1e-05, elementwise_affine=True)
      (2): GELU(approximate='none')
      (3): Dropout(p=0.05, inplace=False)
    )
    (3): Linear(in_features=1280, out_features=1280, bias=True)
  )
  (binary_decoder): Sequential(
    (0): Sequential(
      (0): Linear(in_features=2560, out_features=2048, bias=True)
      (1): LayerNorm((2048,), eps=1e-05, elementwise_affine=True)
```

```
      (2): GELU(approximate='none')
      (3): Dropout(p=0.05, inplace=False)
    )
    (1): Sequential(
      (0): Linear(in_features=2048, out_features=512, bias=True)
      (1): LayerNorm((512,), eps=1e-05, elementwise_affine=True)
      (2): GELU(approximate='none')
      (3): Dropout(p=0.05, inplace=False)
    )
    (2): Sequential(
      (0): Linear(in_features=512, out_features=128, bias=True)
      (1): LayerNorm((128,), eps=1e-05, elementwise_affine=True)
      (2): GELU(approximate='none')
      (3): Dropout(p=0.05, inplace=False)
    )
    (3): Linear(in_features=128, out_features=1, bias=True)
  )
  (gene_embedding_layer): Sequential(
    (0): Linear(in_features=5120, out_features=1280, bias=True)
    (1): GELU(approximate='none')
    (2): LayerNorm((1280,), eps=1e-05, elementwise_affine=True)
  )
)
```

### Structure of tGPT:

```
GPT2Model(
  (wte): Embedding(21150, 1024)
  (wpe): Embedding(1024, 1024)
  (drop): Dropout(p=0.1, inplace=False)
  (h): ModuleList(
    (0): GPT2Block(
      (ln_1): LayerNorm((1024,), eps=1e-05, elementwise_affine=True)
      (attn): GPT2Attention(
        (c_attn): Conv1D()
        (c_proj): Conv1D()
        (attn_dropout): Dropout(p=0.1, inplace=False)
        (resid_dropout): Dropout(p=0.1, inplace=False)
      )
      (ln_2): LayerNorm((1024,), eps=1e-05, elementwise_affine=True)
      (mlp): GPT2MLP(
        (c_fc): Conv1D()
        (c_proj): Conv1D()
        (act): NewGELUActivation()
        (dropout): Dropout(p=0.1, inplace=False)
      )
    )
    (1): GPT2Block(
      (ln_1): LayerNorm((1024,), eps=1e-05, elementwise_affine=True)
      (attn): GPT2Attention(
        (c_attn): Conv1D()
        (c_proj): Conv1D()
        (attn_dropout): Dropout(p=0.1, inplace=False)
        (resid_dropout): Dropout(p=0.1, inplace=False)
      )
      (ln_2): LayerNorm((1024,), eps=1e-05, elementwise_affine=True)
      (mlp): GPT2MLP(
        (c_fc): Conv1D()
        (c_proj): Conv1D()
        (act): NewGELUActivation()
        (dropout): Dropout(p=0.1, inplace=False)
      )
    )
    (2): GPT2Block(
      (ln_1): LayerNorm((1024,), eps=1e-05, elementwise_affine=True)
      (attn): GPT2Attention(
        (c_attn): Conv1D()
        (c_proj): Conv1D()
        (attn_dropout): Dropout(p=0.1, inplace=False)
        (resid_dropout): Dropout(p=0.1, inplace=False)
      )
      (ln_2): LayerNorm((1024,), eps=1e-05, elementwise_affine=True)
      (mlp): GPT2MLP(
        (c_fc): Conv1D()
        (c_proj): Conv1D()
        (act): NewGELUActivation()
        (dropout): Dropout(p=0.1, inplace=False)
      )
```

```
)
(3): GPT2Block(
  (ln_1): LayerNorm((1024,), eps=1e-05, elementwise_affine=True)
  (attn): GPT2Attention(
    (c_attn): Conv1D()
    (c_proj): Conv1D()
    (attn_dropout): Dropout(p=0.1, inplace=False)
    (resid_dropout): Dropout(p=0.1, inplace=False)
  )
  (ln_2): LayerNorm((1024,), eps=1e-05, elementwise_affine=True)
  (mlp): GPT2MLP(
    (c_fc): Conv1D()
    (c_proj): Conv1D()
    (act): NewGELUActivation()
    (dropout): Dropout(p=0.1, inplace=False)
  )
)
(4): GPT2Block(
  (ln_1): LayerNorm((1024,), eps=1e-05, elementwise_affine=True)
  (attn): GPT2Attention(
    (c_attn): Conv1D()
    (c_proj): Conv1D()
    (attn_dropout): Dropout(p=0.1, inplace=False)
    (resid_dropout): Dropout(p=0.1, inplace=False)
  )
  (ln_2): LayerNorm((1024,), eps=1e-05, elementwise_affine=True)
  (mlp): GPT2MLP(
    (c_fc): Conv1D()
    (c_proj): Conv1D()
    (act): NewGELUActivation()
    (dropout): Dropout(p=0.1, inplace=False)
  )
)
(5): GPT2Block(
  (ln_1): LayerNorm((1024,), eps=1e-05, elementwise_affine=True)
  (attn): GPT2Attention(
    (c_attn): Conv1D()
    (c_proj): Conv1D()
    (attn_dropout): Dropout(p=0.1, inplace=False)
    (resid_dropout): Dropout(p=0.1, inplace=False)
  )
  (ln_2): LayerNorm((1024,), eps=1e-05, elementwise_affine=True)
  (mlp): GPT2MLP(
    (c_fc): Conv1D()
    (c_proj): Conv1D()
    (act): NewGELUActivation()
    (dropout): Dropout(p=0.1, inplace=False)
  )
)
(6): GPT2Block(
  (ln_1): LayerNorm((1024,), eps=1e-05, elementwise_affine=True)
  (attn): GPT2Attention(
    (c_attn): Conv1D()
```

```
      (c_proj): Conv1D()
      (attn_dropout): Dropout(p=0.1, inplace=False)
      (resid_dropout): Dropout(p=0.1, inplace=False)
    )
    (ln_2): LayerNorm((1024,), eps=1e-05, elementwise_affine=True)
    (mlp): GPT2MLP(
      (c_fc): Conv1D()
      (c_proj): Conv1D()
      (act): NewGELUActivation()
      (dropout): Dropout(p=0.1, inplace=False)
    )
  )
  (7): GPT2Block(
    (ln_1): LayerNorm((1024,), eps=1e-05, elementwise_affine=True)
    (attn): GPT2Attention(
      (c_attn): Conv1D()
      (c_proj): Conv1D()
      (attn_dropout): Dropout(p=0.1, inplace=False)
      (resid_dropout): Dropout(p=0.1, inplace=False)
    )
    (ln_2): LayerNorm((1024,), eps=1e-05, elementwise_affine=True)
    (mlp): GPT2MLP(
      (c_fc): Conv1D()
      (c_proj): Conv1D()
      (act): NewGELUActivation()
      (dropout): Dropout(p=0.1, inplace=False)
    )
  )
 )
 (ln_f): LayerNorm((1024,), eps=1e-05, elementwise_affine=True)
)
```
